# The Mechanism of MICU-Dependent Gating of the Mitochondrial Ca^2+^ Uniporter

**DOI:** 10.1101/2020.04.04.025833

**Authors:** Vivek Garg, Ishan Paranjpe, Tiffany Unsulangi, Junji Suzuki, Lorin S. Milescu, Yuriy Kirichok

**Affiliations:** Department of Physiology, University of California San Francisco, San Francisco, CA, USA; Department of Physiology, University of Maryland, Baltimore, MD, USA; Department of Biology, University of Maryland, College Park, MD, USA

**Author notes:** Correspondence to (YK); (VG).

## Abstract

Mitochondrial Ca^2+^ uniporter (MCU) mediates mitochondrial Ca^2+^ uptake, regulating ATP production and cell death. According to the existing paradigm, MCU is occluded at the resting cytosolic [Ca^2+^] and only opens above an ∼400 nM threshold. This Ca^2+^-dependent gating is putatively conferred by MICUs, EF hand-containing auxiliary subunits that block/unblock the MCU pore depending on cytosolic [Ca^2+^]. Here we provide the first direct, patch-clamp based analysis of the Ca^2+^-dependent MCU gating and the role played by MICUs. Surprisingly, MICUs do not occlude the MCU pore, and MCU is a constitutively active channel without cytosolic [Ca^2+^] activation threshold. Instead, MICUs potentiate MCU activity when cytosolic Ca^2+^ binds to their EF hands. MICUs cause this potentiation by increasing the probability of open state of the MCU channel.

**One Sentence Summary:** Auxiliary MICU subunits do not occlude the mitochondrial Ca^2+^ uniporter (MCU) but increase its activity as cytosolic Ca^2+^ is elevated.

## Main Text

Mitochondrial Ca^2+^ uptake regulates ATP production, shapes intracellular Ca^2+^ transients and plays a crucial role in deciding cell fate (*1-4*). It is mediated by the mitochondrial Ca^2+^ uniporter (MCU) (*3-5*), which upon elevation of cytosolic [Ca^2+^] ([Ca^2+^]_cyto_) allows selective Ca^2+^ permeation into the mitochondrial matrix, down the high electrochemical gradient across the IMM. All Ca^2+^ channels lose their selectivity and become permeable for Na^+^ at low [Ca^2+^], when Ca^2+^ is removed from the pore (*6-8*). MCU also conducts Na^+^ but only when [Ca^2+^] is decreased to low nM range, because the MCU pore has a Ca^2+^ binding site with an exceptionally high affinity (*K*_*d*_ ≤2 nM) (*9-15*). This prevents permeation of abundant cytosolic monovalent cations even at a resting [Ca^2+^]_cyto_ of ∼100 nM, and makes MCU the most selective Ca^2+^ channel known.

MCU activity must be regulated. Insufficient Ca^2+^ uptake would result in deficient ATP production, whereas excessive uptake would lead to mitochondrial Ca^2+^ overload, ΔΨ dissipation, mitochondrial dysfunction and cell death (*16*). A few early studies suggested that MCU activity might be potentiated by cytosolic [Ca^2+^] (*4, 17, 18*). However, the results differed significantly between labs, because MCU activity was assessed indirectly in suspensions of isolated mitochondria and critical experimental conditions could not be reliably controlled (*3, 4*). Thus, such potentiation was controversial and no clear unifying model for Ca^2+^-dependent MCU gating was generated.

Recent molecular characterization established that MCU is a macromolecular complex (fig. S1A). Its pore is formed by the MCU subunit (*19, 20*) and the essential MCU regulator (EMRE) subunit (*21*). EF hand domain-containing auxiliary MICU1−3 subunits are tethered on the cytosolic side of the MCU/EMRE pore (*22, 23*). MICU1 interacts directly with the MCU and EMRE, while MICU2 and MICU3 attach to the MCU complex only by heterodimerizing with MICU1 (*21, 24-26*). MICU3 is a neuronal- and embryonic-specific isoform with little expression in other tissues (*23, 27*).

The understanding of the molecular composition of the MCU complex renewed interest in the MCU gating by cytosolic Ca^2+^. In MICU1 deficiency, when none of the MICU subunits is associated with the MCU/EMRE pore, mitochondrial Ca^2+^ ([Ca^2+^]_mito_) starts to increase at lower [Ca^2+^]_cyto_ both in cells (*24, 25, 28-31*) and isolated mitochondria (*32*). Based on these results, the term “[Ca^2+^]_cyto_ threshold for mitochondrial Ca^2+^ uptake” was coined, and it was postulated that MICU1 (in association with other MICUs) confers the [Ca^2+^]_cyto_ threshold for MCU activation (*28, 29*). Specifically, the current paradigm suggests that at resting [Ca^2+^]_cyto_, MICU1 occludes the MCU pore (*28, 33, 34*), but when [Ca^2+^]_cyto_ increases above ∼400−800 nM and Ca^2+^ binds to the MICU1 EF hands, this occlusion is relieved (*24, 28, 29*) (Fig. 6A). MICU2 is proposed to facilitate this MICU1 function (*24, 25, 35*). In this model, the occlusion of MCU by MICU1/MICU2 at the resting [Ca^2+^]_cyto_ is considered well-established (tables S1 and S2), while the degree to which the occlusion is removed at elevated [Ca^2+^]_cyto_ remains controversial with different groups reporting a wide range of effects (tables S1 and S2). *MICU1*^*-/-*^ mice show profound late embryonic and postnatal lethality (*32, 36*), while loss-of function MICU1 mutations in humans cause fatigue, lethargy, severe myopathy, developmental and learning disabilities, and progressive extrapyramidal movement disorder (*30, 37-39*).

The paradigm that MICUs occlude the MCU pore at resting cytosolic Ca^2+^ and impart [Ca^2+^]_cyto_ activation threshold on MCU has affected the field profoundly. However, it has never been demonstrated by direct measurement of Ca^2+^ currents mediated by MCU. Instead, MCU activity was inferred from the changes in [Ca^2+^] inside or outside of mitochondria, as measured with Ca^2+^ indicators. However, such [Ca^2+^] changes never reflect MCU activity alone but are determined by the balance between mitochondrial Ca^2+^ uptake and efflux mechanisms (*3, 4, 40*). Some of these studies (*34, 36*) used CGP37157 to inhibit the mitochondrial Ca^2+^ efflux associated with the Ca^2+^/Na^+^ exchange mechanism, but this was clearly insufficient to eliminate all mitochondrial Ca^2+^ efflux. Indeed, if the Ca^2+^ efflux was fully eliminated, the free [Ca^2+^]_mito_ (based on the Nernst equation and assuming 100 nM [Ca^2+^]_cyto_ and ΔΨ at −160 mV) would reach an enormous value of ∼25 mM even with residual MCU activity. Other factors such as ΔΨ, the volume of mitochondrial matrix, matrix Ca^2+^ buffering with phosphates (*40*) and pH can further confound indirect assessment of MCU activity using Ca^2+^ indicators.

The numerous pitfalls associated with indirect assessment of MCU activity make direct measurements of MCU currents (*9, 10, 41*) necessary for understanding of MCU regulation and the role of MICU subunits. However, such direct measurements have been considered extremely challenging, especially in the context of structure–function studies of the MCU complex, which require assessment of numerous knockout and mutant models. There have been a few attempts to characterize MICU1-dependent regulation of MCU using direct electrophysiology, but the scope of electrophysiological experiments in these few studies was very limited, and MICU1 function was assessed only at high [Ca^2+^]_cyto_ (*42-44*). These incomplete electrophysiological studies generated very diverse results ranging from inhibition to no effect of MICU1 on the MCU activity, and thus no clarity was achieved (table S1). One electrophysiological study tried to assess the effects of MICU1 and MICU2 at both low and high [Ca^2+^]_cyto_ (table S1) (*25*). Unfortunately, in this study a recombinant MCU subunit was reconstituted in planar lipid bilayers in the absence of EMRE, the subunit essential for both a functional MCU channel and the association of MICU1/MICU2 with the MCU pore (*25*). The observed channel conducted Na^+^ even at μM [Ca^2+^] and failed to replicate the exceptionally high MCU selectivity for Ca^2+^. Thus, the channel activity observed in this EMRE-less system was artifactual.

Therefore, to facilitate a rigorous and systematic insight into the function of MICU1−3, as well as other subunits of the MCU complex, we developed a heterologous expression system for direct patch-clamp analysis of the MCU complex in the native IMM. Using this system, we demonstrate that MICUs do not occlude the MCU pore. We next demonstrate that the actual function of the MICU subunits is to potentiate MCU activity when their EF hands bind cytosolic Ca^2+^. Thus, MCU has no intrinsic [Ca^2+^]_cyto_ activation threshold. It is a constitutively active channel that is potentiated by [Ca^2+^]_cyto_ via the MICU subunits.

## Results

### System for direct structure–function analysis of MCU

Two factors are crucial to the success of the whole-IMM patch-clamp: the size of individual mitoplasts (vesicles of the whole native IMM) and the IMM stability during the electrophysiological experiments. Therefore, we tested various cell lines for the best possible optimization of these two factors. Eventually, we selected a *Drp1*^*-/-*^ MEF cell line (*45*), in which mitochondria form long tubular networks and provide a significantly higher proportion of large isolated mitoplasts that are also remarkably resilient during the whole-IMM electrophysiological experiments. We confirmed that this cell line expresses all principal subunits of the MCU complex (fig. S2A-D). We next generated gene knockouts for all principal subunits of the MCU complex (MCU, EMRE and MICU1−3) using CRISPR-Cas9 in the background of *Drp1*^*-/-*^ MEFs (fig. S1). All knockout cell lines lacked protein expression of the respective subunit (fig. S2A-C).

To explore the cytosolic/mitochondrial Ca^2+^ phenotypes in these MCU complex knockout cell lines, we induced slow elevation of [Ca^2+^]_cyto_ using the SERCA inhibitor thapsigargin (Tg) and observed an associated increase in [Ca^2+^]_mito_ (fig. S2E-J). [Ca^2+^]_cyto_ was measured using Fura-2 while the mitochondrial Ca^2+^ changes were measured using a genetically-encoded Ca^2+^ indicator *Cepia* (*46*) targeted to mitochondria. [Ca^2+^]_cyto_ under resting conditions was maintained ∼75 nM in all cell lines (fig. S2K) and peaked in the range of 400−1000 nM upon addition of Tg (fig. S2L). In cells with the *WT* MCU complex, the [Ca^2+^]_cyto_ increase was followed, after a short delay, by [Ca^2+^]_mito_ elevation (fig. S2E). However, as expected, in *MCU*^*-/-*^ or *EMRE*^*-/-*^ cell lines that have no functional MCU complex (*19-21*), no significant [Ca^2+^]_mito_ elevation was observed (fig. S2F and G). In MICU1−3-deficient cells, the [Ca^2+^]_cyto_ threshold for elevation of [Ca^2+^]_mito_ was altered as compared to that in cells with the *WT* MCU complex (fig. S2H-J, and M). In *MICU1*^*-/-*^ cells, the threshold was drastically decreased (fig. S2H and M), and a significant but less profound decrease was also observed in *MICU2*^*-/-*^ cells (fig. S2I and M). However, *MICU3*^*-/-*^ cells had an increased threshold (fig. S2J and M). Thus, in our cell system, we observed the same [Ca^2+^]_mito_ phenotypes associated with knockout of individual MCU complex subunits as reported previously (*24, 25, 28, 29, 34*).

We next explored how knockouts for various MCU complex subunits affect MCU currents. Importantly, the MCU complex was intact in isolated whole-IMM vesicles (mitoplasts) used in our patch-clamp experiments, and its composition was the same as in intact mitochondria based on MCU-FLAG co-immunoprecipitation experiments (fig. S4A). Mitoplasts isolated from cells with the *WT* MCU complex had a robust whole-IMM Ca^2+^ current (*I*_Ca_). The voltage step from 0 to −160 mV, followed by a voltage ramp to +80 mV, elicited an inwardly rectifying *I*_Ca_ that gradually increased as [Ca^2+^]_cyto_ (bath solution) was elevated (Fig. 1A, *left panel*, and fig. S3A and B). As expected, in a Ca^2+^-free bath solution (control), we only observed an outward Na^+^ current (*I*_Na_, black trace) via MCU, because the pipette solution contained Na^+^ (Fig. 1A, *left panel*, and fig. S3A and B).

**Fig. 1.**
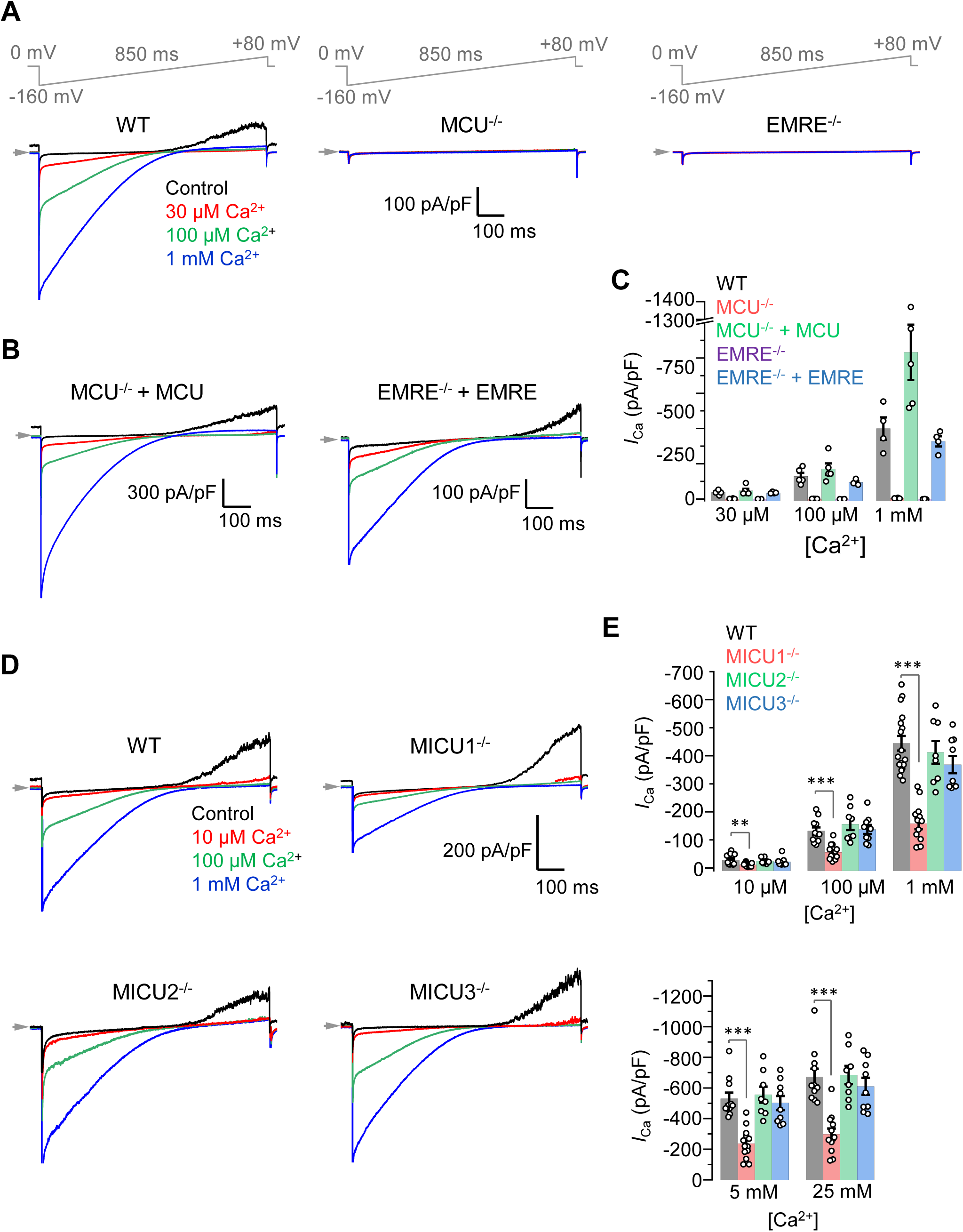
MCU-mediated *I*_Ca_ in *WT* and knockouts of MCU complex subunits. (**A**) Inward *I*_Ca_ elicited by a voltage ramp in *WT, MCU*^*-/-*^ and *EMRE*^*-/-*^ mitoplasts exposed to [Ca^2+^]_cyto_ of 30 μM, 100 μM and 1 mM. In *WT*, also note an outward Na^+^ current via MCU at positive voltages in Ca^2+^-free bath solution (Control). Voltage protocol is indicated on the top. (**B**) *I*_Ca_ is rescued by the recombinant expression of MCU and EMRE in their respective knockout cell lines. (**C**) *I*_Ca_ density measured at −160 mV at different [Ca^2+^]_cyto_ in indicated cell lines. (*n* = 4 to 5 each) Mean ± SEM. (**D**) Inward *I*_Ca_ in *WT, MICU1*^*-/-*^, *MICU2*^*-/-*^ and *MICU3*^*-/-*^ mitoplasts exposed to 10 μM, 100 μM and 1 mM [Ca^2+^]_cyto_.. (**E**) *I*_Ca_ amplitudes measured at −160 mV in mitoplasts at [Ca^2+^]_cyto_ of 10 μM, 100 μM and 1 mM (*upper*), as well as 5 mM and 25 mM (*lower*). (*n* = 8 to 17) Mean ± SEM; one-way ANOVA with post-hoc Tuckey test. \*\**p*< 0.01; \*\*\**p*< 0.001.

Mitoplasts isolated from *MCU*^*-/-*^ and *EMRE*^*-/-*^ lines had no inward *I*_Ca_ or outward *I*_Na_, confirming the essential role of these two subunits for the functional MCU complex (*21, 47, 48*) (Fig. 1A and C). Importantly, even millimolar [Ca^2+^]_cyto_ induced no *I*_Ca_ in *MCU*^*-/-*^ and *EMRE*^*-/-*^, demonstrating that MCU is the only electrogenic mechanism for mitochondrial Ca^2+^ uptake. Heterologous expression of MCU or EMRE in their corresponding knockout cell lines (fig. S4B and C) resulted in restoration of the inward *I*_Ca_ and outward *I*_Na_ (Fig. 1B and C).

Thus, we have identified a system that has robust MCU currents, can be used for heterologous expression of recombinant MCU complex subunits, and significantly improves throughput of whole-IMM patch-clamp recording.

### MICUs are Ca^2+^-dependent MCU potentiators

In contrast to *MCU*^*-/-*^ and *EMRE*^*-/-*^, none of the MICU knockouts (MICU1−3) showed loss of *I*_Ca_ or *I*_Na_ (Fig. 1D), demonstrating that these subunits are not absolutely required for a functional MCU channel. However, among all MICU knockouts, loss of MICU1 resulted in a marked reduction (∼50%) of *I*_Ca_ in both micromolar and millimolar ranges of [Ca^2+^]_cyto_ (Fig. 1D and E, and fig. S5). The same reduction was observed when *I*_Ca_ was measured at both −160 mV (Fig. 1E) and −80 mV (fig. S5C). We next focused on understanding the mechanism by which MICU1 regulates MCU function.

As was suggested previously, MICU1 tethers other MICU subunits to the MCU/EMRE pore (*21, 24-26*). Thus, in *MICU1*^*-/-*^ none of the MICU subunits are associated with the MCU complex. The levels of MCU and MCUb (MCU paralog with no Ca^2+^ transport activity and putative dominant-negative effect on the MCU function) subunits (*49*) were not affected in *MICU1*^*-/-*^, while EMRE expression was significantly reduced (fig. S6A-D), as was also shown previously (*32*). The lower EMRE expression in *MICU1*^*-/-*^ was not a limiting factor for *I*_Ca_, because EMRE overexpression in *MICU1*^*-/-*^ cells did not rescue the *I*_Ca_ reduction (fig. S6D-F). Therefore, the *I*_Ca_ reduction in *MICU1*^*-/-*^ was caused by the lack of MICU1 (and other MICU proteins) in the MCU complex. Because *I*_Ca_ was recorded at [Ca^2+^]_cyto_ ≥ 10 μM, when the EF hands of MICU subunits (*K*_d_ ∼600 nM) (*50*) are occupied by Ca^2+^, we conclude that in the Ca^2+^-bound state MICUs potentiate the MCU current.

We next studied how MICUs affect the MCU current when Ca^2+^ is not bound to their EF hands. Because this requires [Ca^2+^]_cyto_ <60 nM (10-fold less than *Kd*) and *I*_Ca_ cannot be measured reliably under these conditions, we used Na^+^ as the permeating ion. A robust *I*_Na_ via MCU was observed when Ca^2+^ was eliminated on the cytosolic face of the IMM with Ca^2+^ chelators (Fig. 2A, *left panel*). As expected, *I*_Na_ completely disappeared in *MCU*^*-/-*^ and *EMRE*^*-/-*^ (Fig. 2A and B). Interestingly, in a striking contrast to *I*_Ca_, *I*_Na_ was not reduced in *MICU1*^*-/-*^, (Fig. 2C-E, also see Fig. 1D and E). The very presence of a robust *I*_Na_, and the fact that it is not altered in *MICU1*^*-/-*^, argues strongly against the currently accepted paradigm (*33, 34*) in which the MCU/EMRE pore is occluded by MICUs when their EF hands are not occupied by Ca^2+^ (Fig. 6A). In the absence of cytosolic Ca^2+^, a robust *I*_Na_ via MCU was also previously recorded in mitoplasts isolated from COS-7 cells, mouse heart and skeletal muscle (*9, 10*). Nanomolar concentrations of MCU inhibitor ruthenium red (RuR) completely block this *I*_Na_ (*9, 10*). In the absence of divalent cations, a RuR-sensitive, Na^+^-selective MCU-dependent uniport was also reported in intact isolated mitochondria (*51, 52*). Thus, the MCU/EMRE pore is not occluded by MICU proteins when Ca^2+^ is not bound to their EF hands. Moreover, the similarity of *I*_Na_ amplitudes in *WT* and *MICU1*^*-/-*^ (Fig. 2C-E) suggests that in their Ca^2+^-free state MICUs do not affect ion permeation through the MCU/EMRE pore at all.

**Fig. 2.**
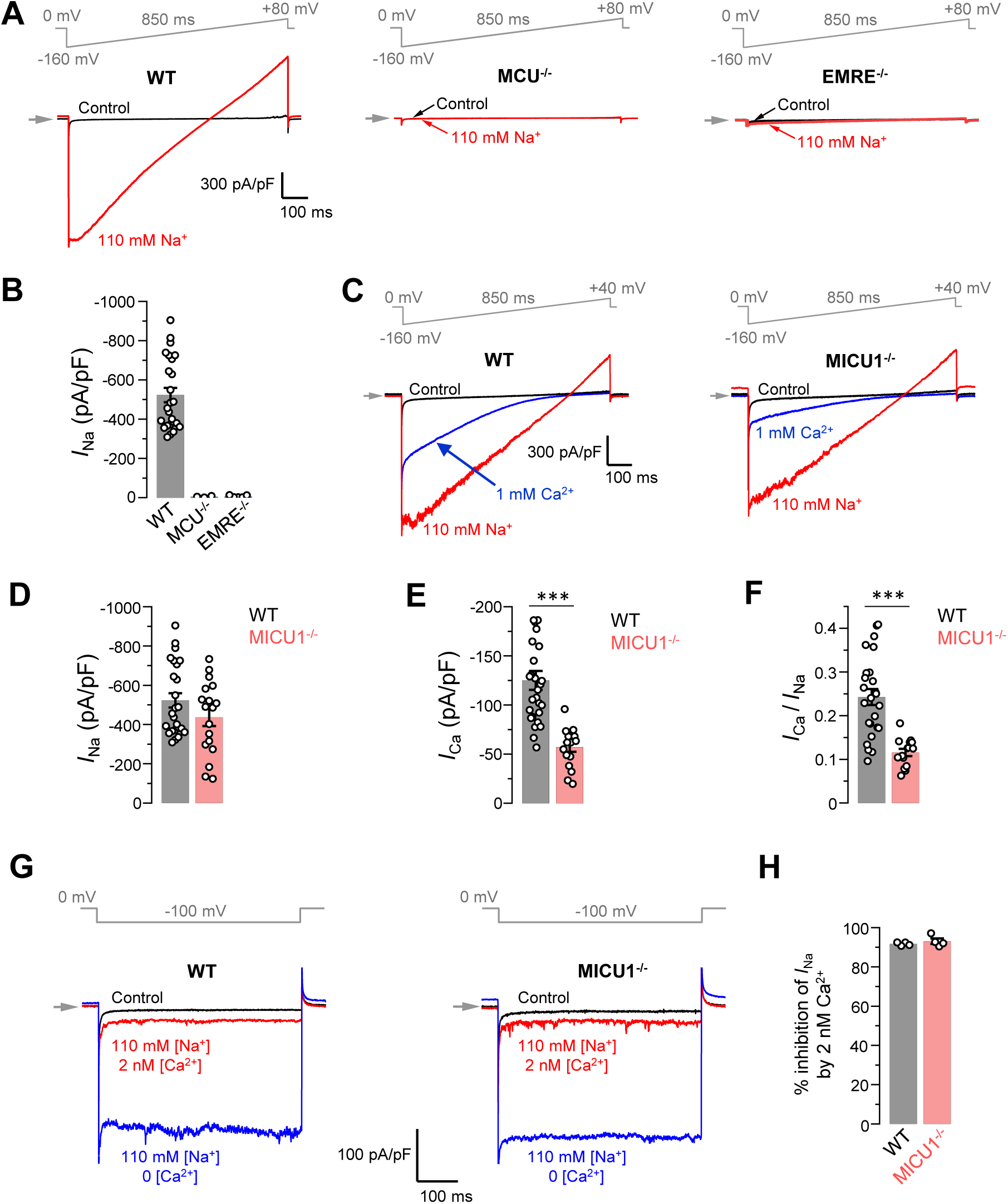
MICU1 is a Ca^2+^-dependent MCU potentiator. (**A**) Representative *I*_Na_ in *WT, MCU*^*-/-*^ and *EMRE*^*-/-*^ mitoplasts at 110 mM [Na^+^]_cyto_. (**B**) *I*_Na_ amplitudes measured at −80 mV in *WT* (*n* = 20), *MCU*^*-/-*^ (*n* = 3) and *EMRE*^*-/-*^ (*n* = 3) mitoplasts. (**C**) Representative *I*_Ca_ (*blue*) and *I*_Na_ (*red*) recorded from the same *WT* and *MICU1*^*-/-*^ mitoplasts exposed to 1 mM [Ca^2+^]_cyto_ or 110 mM [Na^+^]_cyto_. (**D** to **F**), Amplitudes of *I*_Na_ (D) and *I*_Ca_ (E), and the *I*_Ca_/*I*_Na_ ratio in the same mitoplast (F) in *WT* (*n* = 27) and *MICU1*^*-/-*^ (*n* = 18). Current were measured at −80 mV. Mean ± SEM; unpaired t-test, two-tailed. \*\*\**p*< 0.001. (**G**) Inward *I*_Na_ recorded in the absence of cytosolic Ca^2+^ (*blue*) and subsequently at 2 nM [Ca^2+^]_cyto_ (*red*) in *WT* (*left*) and *MICU1*^*-/-*^ (*right*) mitoplasts exposed to 110 mM [Na^+^]_cyto_. (**H**) Inhibition of *I*Na by 2 nM [Ca^2+^]cyto in *WT* and *MICU1*^*-/-*^. Mean ± SEM; unpaired t-test, two-tailed (*n* = 4 each).

The *MICU1*^*-/-*^ phenotypes of *I*_Na_ (no change) and *I*_Ca_ (reduction) suggest that the only function of MICUs is potentiation of the MCU complex activity when their EF hands are occupied by Ca^2+^. To further examine this phenotype, we studied how the ratio of *I*_Ca_ to *I*_Na_, as measured in the same mitoplast, is affected by *MICU1*^*-/-*^. Such *I*_Ca_*/I*_Na_ ratio depends only on the functional properties of the MCU complex, and, in contrast to *I*_Ca_ and *I*_Na_ amplitudes, is independent of the number of MCU complexes in a mitoplast. Thus, an alteration of the *I*_Ca_*/I*_Na_ ratio in *MICU1*^*-/-*^ can be directly attributed to altered functional properties of the MCU complex, and would not depend on any associated changes in MCU/EMRE expression affecting the number of MCU complexes.

The *I*_Ca_/*I*_Na_ ratio was dramatically reduced in *MICU1*^*-/-*^ mitoplasts (Fig. 2F), which means that the loss of MICUs is directly responsible for the reduction of *I*_Ca_ as compared to *I*_Na_. The loss of MICUs can cause such reduction in the *I*_Ca_/*I*_Na_ ratio by either altering the channel gating or affecting the relative affinities for Ca^2+^ and Na^+^ binding in the selectivity filter. The reduction in *I*_Ca_/*I*_Na_ ratio in *MICU1*^*-/-*^ could not be explained by altered relative affinities for Ca^2+^ and Na^+^ binding in the selectivity filter, because *I*_Na_ was inhibited to the same extent by 2 nM [Ca^2+^]_cyto_ in both *WT* and *MICU1*^*-/-*^ mitoplasts (Fig. 2G and H). Thus, a reduced *I*_Ca_/*I*_Na_ ratio in *MICU1*^*-/-*^ is caused by a disrupted MICU-dependent gating mechanism. This gating mechanism potentiates MCU currents in a Ca^2+^-dependent fashion.

The above experiments were all performed in cell lines with disrupted *Drp1*. However, *Drp1* is not a part of the MCU complex, and therefore *Drp1* loss is not expected to affect MCU currents as measured directly with patch-clamp electrophysiology. Indeed, in our experiments *Drp1* knockout did not affect the amplitudes of *I*_Ca_ or *I*_Na_ mediated by the MCU complex (fig. S7A and B). However, we still confirmed that the observed *MICU1*^*-/-*^ current phenotypes were the same, irrespective of the *Drp1* background. Similar to MICU1 knockout in *Drp1*^*-/-*^ MEFs, MICU1 knockout in *Drp1*^*+/+*^ MEFs did not affect *I*_Na_ while markedly reduced *I*_Ca_ (fig. S7C-E). Additionally, *MICU1*^*-/-*^ reduced the *I*_Ca_/*I*_Na_ ratio, as measured in the same mitoplast, to the similar extent in *Drp1*^*+/+*^ MEFs (fig. S7F). Thus, as expected, *Drp1* presence or absence does not affect currents mediated by the MCU complex or the *MICU1*^*-/-*^ phenotypes.

To conclude, MICU subunits do not occlude the MCU/EMRE pore or impart a [Ca^2+^]_cyto_ activation threshold on the MCU complex. Instead, MCU is a constitutively active channel, and the actual function of MICU subunits is to potentiate MCU currents as [Ca^2+^]_cyto_ is elevated.

### Role of EF hands of MICUs in *I*_Ca_ potentiation

To confirm that Ca^2+^ binding to the EF hands of MICUs is responsible for the Ca^2+^-dependent potentiation of MCU, we recombinantly expressed MICU1–3 or MICU1–3 with mutated EF hands (mut-EF-MICU, to disable Ca^2+^ binding (*24*)) in their respective knockout cell lines and examined the changes in *I*_Ca_ (Fig. 3A).

**Fig. 3.**
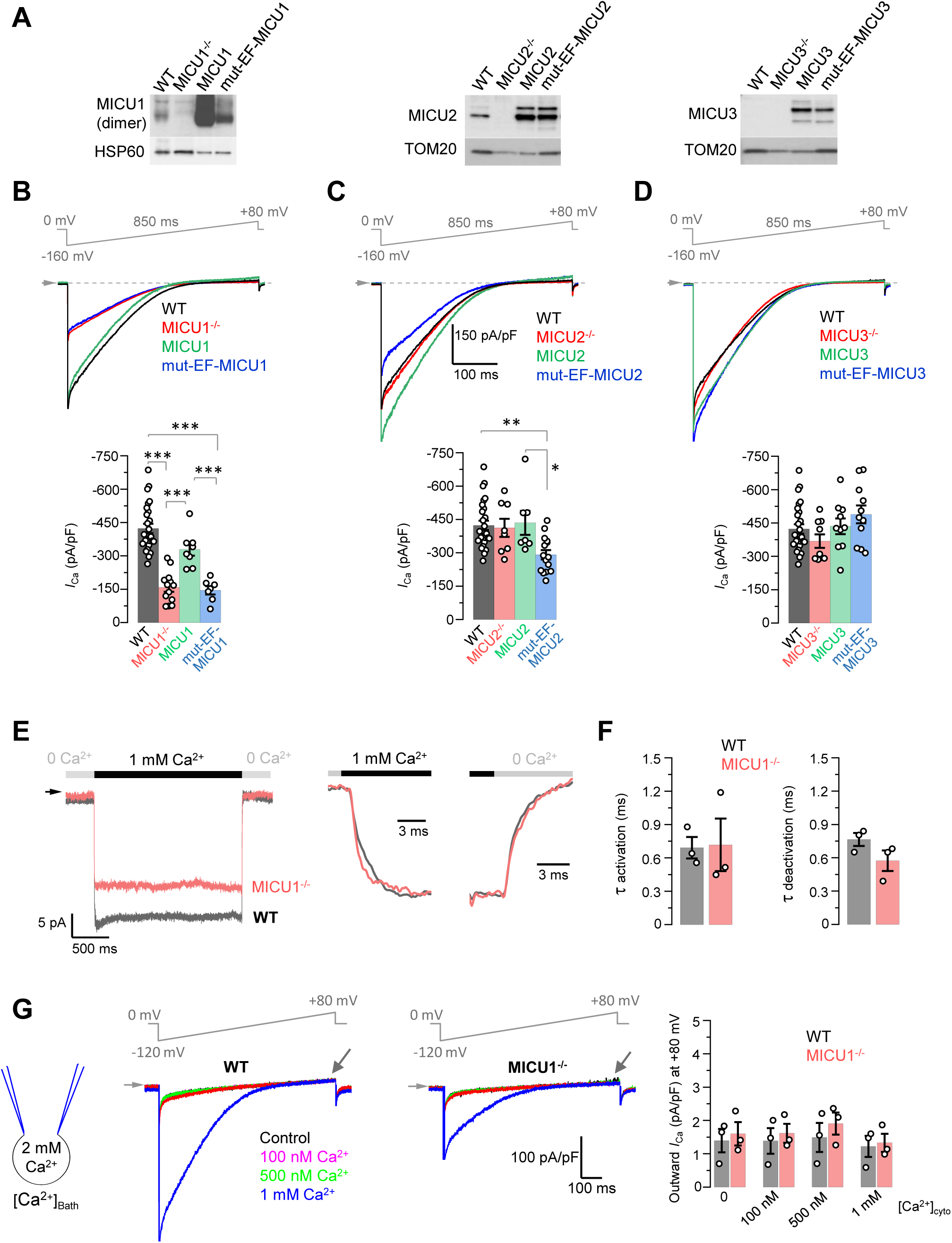
Effects of MICU proteins and their EF hands on the amplitude, kinetics and rectification of *I*_Ca_. (**A**) Western blots showing overexpression of MICU proteins or MICU proteins with non-functional EF hands (mut-EF-MICU) in their respective knockout background (*left, MICU1*^*-/-*^; *middle, MICU2*^*-/-*^ and; *right, MICU3*^*-/-*^). (**B** to **D**) *Upper panels: I*_Ca_ in *MICU1*^*-/-*^ (B), *MICU2*^*-/-*^ (C) and *MICU3*^*-/-*^ (D) before and after overexpression of a corresponding MICU subunit or its EF hand mutant, as compared to *WT*. To simplify comparison, representative *I*_Ca_ traces recorded from the mitoplasts of different backgrounds in 1 mM [Ca^2+^]_cyto_ are shown together in a single panel. *Lower panels:* quantification of *I*_Ca_ amplitudes from the upper panel at −160 mV. The same *WT* and knockout data were used as in Fig. 1e. Mean ± SEM; one-way ANOVA with post-hoc Tuckey test (*n* = 7 to 26). \**p*< 0.05; \*\**p*< 0.01; \*\*\**p*< 0.001. (**E**) *Left panel*: *I*_Ca_ measured at a holding voltage of −100 mV while [Ca^2+^]_cyto_ was rapidly (τ ∼0.4 ms, see Methods) switched from virtual zero to 1 mM and then back to virtual zero in *WT* (*grey*) and *MICU1*^*-/-*^ (*red*) mitoplasts. *Right panel, I*_Ca_ kinetics within ∼10 ms after the fast [Ca^2+^]_cyto_ elevation and subsequent decrease in *WT* (*grey*) and *MICU1*^*-/-*^ (*red*) mitoplasts from the left panel. *I*_Ca_ traces were normalized to the maximal amplitude to facilitate comparison of kinetics in *WT* and *MICU1*^*-/-*^. (**F**) *Left*: *I*_Ca_ activation time constant (*τ*_*a*_) in *WT* and *MICU1*^*-/-*^; *Right*: *I*_Ca_ deactivation time constant (*τ*_*d*_) in *WT* and *MICU1*^*-/-*^. Mean ± SEM (*n* = 3, each). (**G**) *I*_Ca_ at [Ca^2+^]_mito_ = 2 mM and indicated [Ca^2+^]_cyto_ in *WT* and *MICU1*^*-/-*^. Arrows point out where the amplitude of outward *I*_Ca_ was measured. Bar-graph shows the amplitude of outward *I*_Ca_ measured at +80 mV. *n* = 3, each [Ca^2+^]_cyto_.

In *MICU1*^*−/−*^, expression of MICU1 was able to restore *I*_Ca_ to the *WT* level, but mut-EF-MICU1 expression failed to do so (Fig. 3B). Expression levels of both the recombinant MICU1 and mut-EF-MICU1 were significantly higher as compared to MICU1 expression in the cells with *WT* MCU complex (Fig. 3A). This confirms our hypothesis that Ca^2+^ binding to the EF hands of MICU1 is indispensable for *I*_Ca_ potentiation.

In *MICU2*^*−/−*^, *I*_Ca_ was not significantly affected (Fig. 3C, and 1D and E), because the loss of MICU2 appeared to be compensated with increased MICU1 expression and replacement of MICU1/MICU2 heterodimer with MICU1/MICU1 homodimer (fig. S8A-C). Therefore, overexpression of recombinant MICU2 in *MICU2*^*-/-*^ and preferential conversion of MICU1/MICU1 homodimers back into MICU1/MICU2 heterodimers also did not alter the *I*_Ca_ amplitude (Fig. 3C). In contrast, mut-EF-MICU2 overexpression displaced MICU1 from MICU1/MICU1 homodimers in favor of MICU1/mut-EF-MICU2 heterodimer, leading to a dominant-negative effect and a significant decrease in MICU-dependent *I*_Ca_ potentiation (Fig. 3C). These functional data, combined with biochemical evidence for MICU1/MICU2 heterodimers (*25, 53, 54*), suggest that MICU2, along with MICU1, is responsible for allosteric potentiation of MCU upon binding of cytosolic Ca^2+^ to their EF hands.

The effect of *MICU1*^*-/-*^ on *I*_Ca_ was more profound as compared to that of *MICU2*^*-/-*^ (Fig. 1D and E), because MICU1 could compensate for MICU2. However, the reverse compensation was impossible, because only MICU1 tethers the MICU1/MICU2 heterodimer to the MCU/EMRE pore. The composition of MICU dimers can also be affected by MICU3 that similar to MICU2 was proposed to interact and form heterodimers with MICU1 (*27*). However, in our experiments, we did not observe robust current phenotypes associated with MICU3.

Specifically, *I*_Ca_ was not affected in *MICU3*^*-/-*^ mitoplasts, and overexpression of recombinant MICU3 or mut-EF-MICU3 in *MICU3*^*-/-*^ also had no effect on *I*_Ca_ (Fig. 3D). It has been suggested that MICU3 is a minor protein as compared to MICU1 and 2 in the majority of tissues and cell lines assessed (*27*). Although in our system the amount of MICU3 mRNA appeared to be comparable with that of other MICU subunits (fig. S2D), and the MICU3 protein was expressed (Fig. 3A *right panel*, and fig. S2C), the relative abundance of MICU3 vs other MICUs is not clear. Moreover, in contrast to MICU1, MICU3 is not upregulated in *MICU2*^*-/-*^ cells (fig. S8B), and thus, MICU3 expression does not appear to be linked to the level of MICU2. Therefore, although MICU3 could in principle support the Ca^2+^-dependent potentiation of MCU by forming dimers with MICU1 (*27*), the exact role of MICU3 and its interaction with other MCU complex subunits remains to be established.

Ca^2+^ binding to the EF hands of MICU subunits and a subsequent conformational change that potentiates the MCU complex activity require a finite time and may delay *I*_Ca_ activation/deactivation in response to changes in [Ca^2+^]_cyto_. Such delayed *I*_Ca_ kinetics can profoundly affect [Ca^2+^]_mito_, because in situ MCU takes up Ca^2+^ from Ca^2+^ microdomains (*55*) that exist in the cytosol only for a few milliseconds (*56*). Therefore, we examined *I*_Ca_ activation and deactivation kinetics in response to rapid changes in [Ca^2+^]_cyto_ and tested whether they depend on MICUs. *I*_Ca_ activation upon rapid elevation of [Ca^2+^]_cyto_ from virtually Ca^2+^-free to 1 mM was immediate, with kinetics comparable to the rate of solution exchange (τ ∼0.4 ms) achieved by our piezoelectric fast application system (Fig. 3E). Importantly, the kinetics of the *I*_Ca_ rapid response was not altered in *MICU1*^*-/-*^ (Fig. 3E and F). The deactivation kinetics was similarly fast and not dependent on MICU1 (Fig. 3E and F). The result of these experiments correspond to the previous observation that EF hands of calmodulin bind Ca^2+^ with a μs time constant (*57*). The conclusion from these experiments is that the kinetics of Ca^2+^ binding to the MICU’s EF hands, and the resultant conformational change in the MCU complex, are extremely fast, and thus MICU-dependent potentiation of the MCU activity should occur instantaneously upon elevation of the [Ca^2+^]_cyto_. This is perhaps true even within Ca^2+^ microdomains, but it has to be taken into account that in our experiments we used somewhat higher [Ca^2+^]_cyto_ (1 mM) as compared to the maximal [Ca^2+^]_cyto_ achieved in the microdomains (100 μM).

A phenomenon of Ca^2+^-induced mitochondrial Ca^2+^ release (mCICR) by which mitochondria release Ca^2+^ into cytosol in response to elevations of [Ca^2+^]_cyto_ has been observed (*4, 58, 59*). mCICR required mitochondrial depolarization and was proposed to be mediated by MCU (*60, 61*) and/or the permeability transition pore (PTP) (*58, 59*). Therefore, we tested whether MCU can mediate Ca^2+^-dependent Ca^2+^ efflux at depolarized membrane voltages and whether such efflux is dependent on MICUs. We measured outward *I*_Ca_ at positive voltages with 2 mM [Ca^2+^]_mito_ (the pipette solution), as [Ca^2+^]_cyto_ was gradually elevated from virtual zero to 1 mM. Remarkably, such [Ca^2+^]_cyto_ elevation failed to induce any outward *I*_Ca_. However, as expected, it caused a robust inward *I*_Ca_ (Fig. 3G). This experiment also demonstrates, as was also suggested previously (*9*), that MCU has a strong inward rectification (unidirectional Ca^2+^ permeation into the matrix). This strong inward rectification of MCU under various [Ca^2+^]_cyto_ remained unaltered in *MICU1*^*-/-*^ (Fig. 3G). Thus, MCU has a strong preference for conducting Ca^2+^ into mitochondria and is unlikely to mediate Ca^2+^ release via mCICR.

It has also been suggested that MCU is regulated by matrix [Ca^2+^] (*62*). Specifically, *I*_Ca_ was shown to be profoundly reduced at [Ca^2+^]_mito_ ∼400 nM, as compared to that at both lower (Ca^2+^-free) and higher (high μM) [Ca^2+^]_mito_ (*62*). The authors also proposed that the reduction of the MCU current at [Ca^2+^]_mito_ ∼400 nM is MICU1-dependent. However, in contrast to this previous observation, in our experiments *I*_Ca_ amplitude remained unaltered when [Ca^2+^]_mito_ was set at Ca^2+^-free, 400 nM, or 400 μM (fig. S9). Thus, the MCU complex is not regulated by matrix Ca^2+^, and MICUs only impart the regulation of the MCU complex by cytosolic Ca^2+^. It should also be mentioned that the authors proposed a membrane topology of EMRE (*62*) that is reverse to that determined in the recent biochemical and structural studies (*14, 26*).

Taken together, these data indicate that MICU proteins allosterically potentiate MCU-mediated Ca^2+^ influx when cytosolic Ca^2+^ binds to their EF hands.

### MICUs increase the open probability of MCU

To investigate the mechanism by which Ca^2+^-bound MICU proteins potentiate *I*_Ca_, we examined the activity of single MCU channels in inside-out (matrix-side out) IMM patches. Because the unitary Ca^2+^ current (*i*_Ca_, current via a single MCU channel) is very small (<1 pA), it must be recorded at high [Ca^2+^]=105 mM to improve resolution (*9*). At this [Ca^2+^], EF hands of MICU subunits are fully saturated with Ca^2+^.

MCU exhibits multiple levels of single channel conductance. These subconductances can be observed at all tested voltages (−40, −80, and −120 mV), but their resolution improves markedly as transmembrane voltage becomes more negative. At −120 mV, in addition to what appears to be a fully open *i*_Ca_, subconductances at ∼0.8 and ∼0.6 of the amplitude of the fully open *i*_Ca_ can be easily detected (Fig. 4A and C). Because similar amplitude levels were observed in all the patches, we conclude that these events represent genuine subconductances in the MCU channel.

**Fig. 4.**
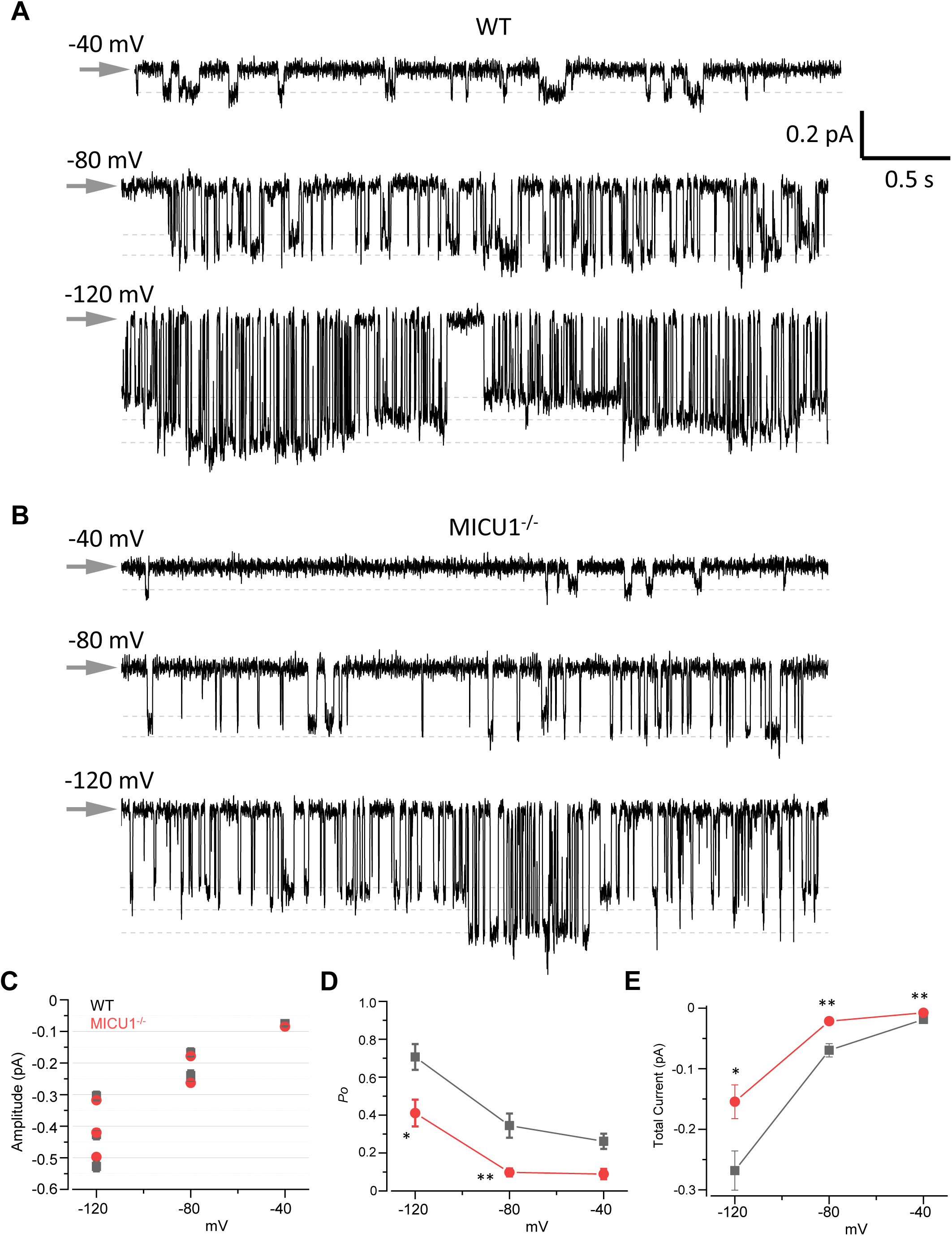
Open probability of the MCU channel is decreased in *MICU1*^*-/-*^. (**A** and **B**) MCU single-channel currents (*i*_Ca_) from inside-out IMM patches in *WT* (A) and *MICU1*^*-/-*^ (B) recorded at indicated potentials in symmetrical 105 mM Ca^2+^, and low-pass filtered at 0.3 kHz for display purposes. Arrows indicate closed state level, and downward deflections are the open state events. Multiple subconductance levels are clearly visible at −80 and −120 mV. (**C** to **E**) Single-channel amplitudes (C), open probability (*P*_o_) (D), and time-averaged unitary current (E) (see Methods) in *WT* and *MICU1*^*-/-*^ at indicated potentials. Mean ± SEM; unpaired t-test, two-tailed; *n* = 5−6, each. \**p*< 0.05; \*\**p*< 0.01.

There was no difference in the single channel amplitude between control and *MICU1*^*-/-*^ mitoplasts (Fig. 4A-C). However, we found that the single-channel open probability (*P*o) was significantly decreased ∼2–3 fold in *MICU1*^*-/-*^ versus *WT* mitoplasts, depending on the transmembrane voltage (Fig. 4A, B, D and fig. S10A-C). As a result, the time-averaged current contributed by a single MCU channel differs significantly between control and *MICU1*^*-/-*^ mitoplasts (Fig. 4E), thus mirroring and explaining the effect of MICU1 knockout on the amplitude of the whole-mitoplast *I*_Ca_ (Fig. 1E).

These results demonstrate that the potentiating effect of MICU proteins on the MCU/EMRE pore is not associated with an increased single-channel conductance. Rather, when their EF hands bind Ca^2+^, MICUs increase MCU currents by causing an increase in the open probability of the MCU/EMRE pore.

### MICU1 does not affect the Mn^2+^ vs Ca^2+^ permeability of MCU

While Mn^2+^ is essential for the proper function of several mitochondrial enzymes, its excessive accumulation inhibits oxidative phosphorylation and causes toxicity (*63*). MCU appears to be the primary pathway for Mn^2+^ entry into mitochondria (*4*). Recently, it has been suggested that MICU1 is responsible for the relatively low permeability of MCU for Mn^2+^ as compared to Ca^2+^, and MICU1 deficiency or loss-of-function MICU1 mutations in patients can lead to excessive mitochondrial Mn^2+^ accumulation and cellular toxicity (*42, 64*). These observations were explained within the paradigm in which MICU1 occludes the MCU/EMRE pore. It was postulated that Mn^2+^ binds to MICU1 EF hands but, in contrast to Ca^2+^, cannot induce the MICU1 conformation change necessary to unblock the MCU pore. Thus, MICU1 prevents Mn^2+^ permeation via MCU and ensures selective Ca^2+^ permeation (*42, 64*).

We recorded the inward Mn^2+^ current (*I*_Mn_) in the presence of 5 mM [Mn^2+^]_cyto_. *I*_Mn_ disappeared in *MCU*^*-/-*^ and *EMRE*^*-/-*^, confirming that it was solely mediated by MCU (Fig. 5A-D). As was also shown previously (*9*), *I*_Mn_ was indeed significantly smaller (∼7-fold) than *I*_Ca_ at 5 mM [Mn^2+^]_cyto_ and 5 mM [Ca^2+^]_cyto_, respectively (Fig. 5E). However, we also observed that *I*_Mn_ and *I*_Ca_ were reduced to a similar extent in *MICU1*^*-/-*^ (Fig. 5F-H). This result was in a striking contrast to the current MICU1-based model for Mn^2+^ vs Ca^2+^ selectivity of MCU (*42, 64*), under which *I*_Mn_ would be increased but *I*_Ca_ not affected under our experimental conditions. Moreover, even the ratio between *I*_Mn_ and *I*_Ca_ calculated from the same mitoplast (*I*_Mn_/*I*_Ca_) was not affected in *MICU1*^*-/-*^ (Fig. 5I), although it is expected to be decreased as per the MICU1-based model for Mn^2+^ vs Ca^2+^ selectivity of MCU (*42, 64*). Finally, Mn^2+^ inhibited *I*_Ca_ to the same extent in *WT* and *MICU1*^*-/-*^ mitoplasts (Fig. 5F and J), indicating that Ca^2+^ and Mn^2+^ are likely to compete in the selectivity filter of the MCU/EMRE pore.

**Fig. 5.**
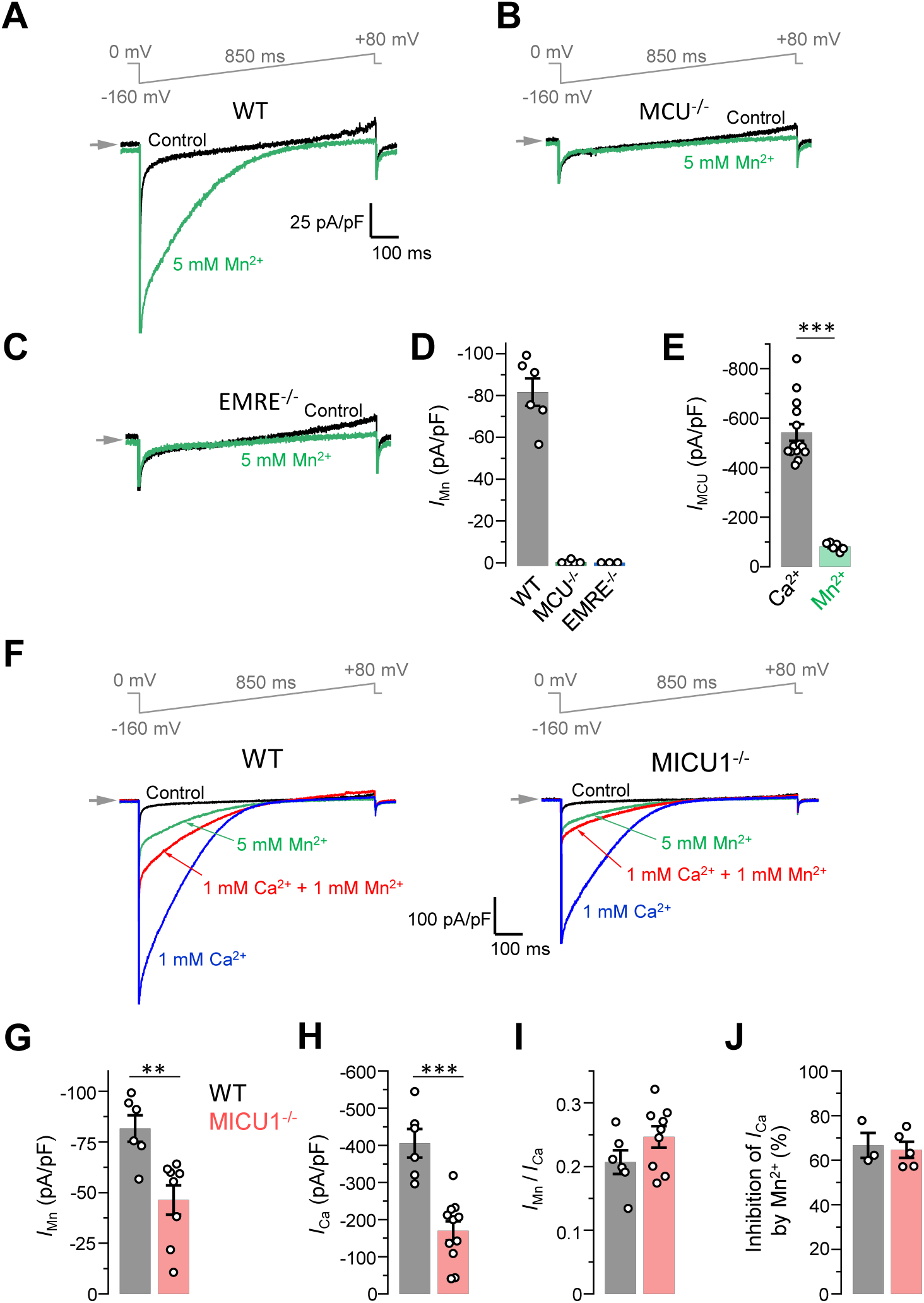
*I*_Mn_ is reduced in *MICU1*^*-/-*^ to the similar extent as *I*_Ca_. (**A** to **C**) Representative inward *I*_Mn_ in *WT* (A), *MCU*^*-/-*^ (B) and *EMRE*^*-/-*^ (C) mitoplasts at 5 mM [Mn^2+^]_cyto_. (**D**) *I*_Mn_ measured at −160 mV from *WT* (*n* = 6), *MCU*^*-/-*^ (*n* = 5) and *EMRE*^*-/-*^ (*n* = 3) mitoplasts. Mean ± SEM. (**E**) *I*_MCU_ amplitudes at 5 mM [Ca^2+^]_cyto_ and 5 mM [Mn^2+^]_cyto_ in *WT* mitoplasts. Currents were measured at −160 mV. Mean ± SEM; unpaired t-test, two-tailed; *n* = 6−14; \*\*\**p*< 0.001. (**F**) Representative *I*_Ca_ (*blue*, [Ca^2+^]_cyto_=1 mM), *I*_Mn_ (*green*, [Mn^2+^]_cyto_=5 mM) and inhibition of *I*_Ca_ by Mn^2+^ (*red*, [Ca^2+^]_cyto_=1 mM and [Mn^2+^]_cyto_=1 mM) in *WT* and *MICU1*^*-/-*^ mitoplasts. (**G** to **J**) *I*_Mn_ (G), *I*_Ca_ (H), *I*_Mn_/*I*_Ca_ ratio (I, measured in the same mitoplasts), and inhibition of *I*_Ca_ by 1 mM [Mn^2+^]_cyto_ (J) in *WT* (*n* = 3−6) and *MICU1*^*-/-*^ (*n* = 5−11). Mean ± SEM; unpaired t-test, two-tailed. \*\**p*< 0.01; \*\*\**p*< 0.001.

Thus, the *I*_Ca_ and *I*_Mn_ phenotypes of *MICU1*^*-/-*^ are the same, and MICU1 does not determine the preference of MCU for Ca^2+^ over Mn^2+^. Permeation of both Ca^2+^ and Mn^2+^ is enhanced, rather than inhibited by MICU1. Instead of MICU1, the selectivity of the MCU complex for Ca^2+^ over Mn^2+^ (and for any other ion) should be determined by the selectivity filter located in the pore (*11-13, 15*), exactly as in other ion channels. Thus, the properties of Mn^2+^ permeation via MCU cannot be explained within the paradigm in which MICUs occlude the MCU/EMRE pore, nor it can be used to validate it.

## Discussion

In summary, the direct patch-clamp analysis presented here argues for a significant revision of the current paradigm for the gating of the MCU complex, its control by [Ca^2+^]_cyto_, and the role played by MICU subunits (Fig. 6).

**Fig. 6.**
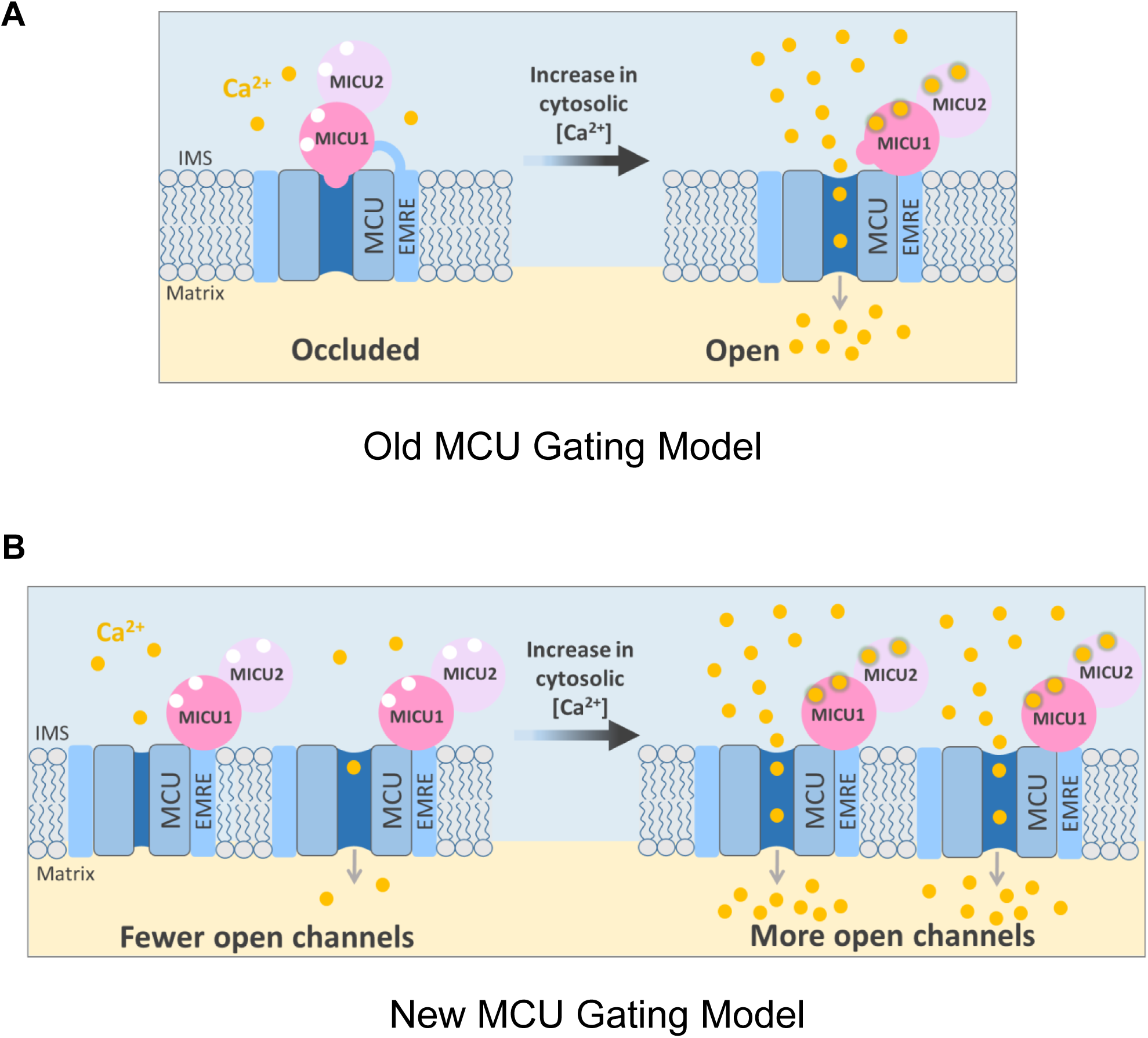
Gating models of the MCU complex. (**A**) Current model of the MCU complex gating and the role of MICU subunits. The MCU complex has two states: MICU-occluded and open. At low [Ca^2+^]_cyto_, MICU subunits occlude the MCU pore and inhibit Ca^2+^ influx. As [Ca^2+^]_cyto_ is increased, Ca^2+^ binds to the EF hands of MICU subunits, the MICU-mediated occlusion is relieved, and the MCU pore is open. (**B**) New model of the MCU complex gating and the role of MICU subunits. The MCU complex is a constitutively active channel. The level of the MCU activity is determined by spontaneous transitions between the open and closed states and the equilibrium between them. At low [Ca^2+^]_cyto_, this equilibrium is such that the probability of the open and closed states are comparable. As [Ca^2+^]_cyto_ is increased and Ca^2+^ binds to the EF hands of MICU subunits, MICUs strongly shift the equilibrium to the open state, which leads to a significant increase in the probability of the open state (*Po*) and a robust increase in the MCU activity.

In contrast to the existing model, we demonstrated that at low [Ca^2+^]_cyto_, when EF hands of MICU subunits are Ca^2+^-free, the MCU/EMRE pore is not occluded by MICUs and conducts robust Na^+^ current regardless of MICU’s presence. Thus, the MCU complex is a constitutively active channel. We further demonstrated that the real function of MICU subunits is to potentiate the activity of the MCU complex as cytosolic Ca^2+^ is elevated and binds to MICU’s EF hands (Fig. 6B). MICU subunits potentiate MCU activity by increasing the open state probability of the MCU/EMRE pore. MICUs are likely to achieve this effect by interacting with EMRE that is predicted to control the gating of the MCU/EMRE pore (*14*). The recent cryo-EM structures of the complete MCU complex clarify this mechanism further and suggests that MICU1/MICU2 dimers connect (at the cytosolic side) EMREs of two different MCU/EMRE pores and could control MCU gating by pulling on these EMREs.

MICU1/MICU2 dimer binds cytosolic Ca^2+^ with *K*_d_ ∼600 nM (*50*). To understand the Ca^2+^-dependent function of the MICU1/MICU2 dimer, the effects of MICUs on the MCU/EMRE pore must be studied at the two extremes of the EF-hand Ca^2+^ titration range – when all MICUs are essentially Ca^2+^-free and when they are fully occupied by Ca^2+^. This can be achieved by measuring the two types of currents via the MCU complex - *I*_Na_ and *I*_Ca._

As we demonstrated previously (*9, 10*) and also elaborate in this work, the MCU complex can conduct both Na^+^ and Ca^2+^. This is because Ca^2+^ and Na^+^ have very similar radii and both can bind to and permeate through the narrowest Ca^2+^ binding site in the MCU selectivity filter, similar to other Ca^2+^ channels (*6, 11-13*). *I*_Na_ and *I*_Ca_ have been instrumental in understanding the MCU channel and its exceptionally high Ca^2+^ selectivity (*9, 10*). *I*_Na_ is measurable only when [Ca^2+^]_cyto_ ≤ 2 nM (Fig. 2G), while *I*_Ca_ can only be measured at [Ca^2+^]_cyto_ ≥ 10 μM (Fig. 1D). In between 2 nM and 10 μM, lies a [Ca^2+^]_cyto_ range where MCU currents are extremely small and cannot be measured reliably. Such “no-current” range is not a unique property of MCU, but is a characteristic property of all Ca^2+^-selective channels and is explained by the anomalous mole-fraction effect, a phenomenon of binding and competition between two different ions (Na^+^ and Ca^2+^) in the selectivity filter (*65*). However, by measuring *I*_Na_ and *I*_Ca_, the patch-clamp electrophysiology can reliably establish the effect of MICUs on the MCU/EMRE pore at the two extremes of the EF-hand Ca^2+^ titration range – when all EF hands are either in the Ca^2+^-free state or in the Ca^2+^-occupied state. It is important to understand that such direct measurement of *I*_Na_ and *I*_Ca_ is the only reliable way to assess the function of MICU subunits within the MCU complex.

It is tempting to assume that optical methods can assess the MCU complex activity and the MICU function continuously over a wide [Ca^2+^]_cyto_ range, starting from high nM. This perceived “high sensitivity” of optical methods is achieved by integration of the net mitochondrial Ca^2+^ influx (even if it is very slow) over a period of time, resulting in a measurable [Ca^2+^]_mito_ change. However, it must be realized that the optical methods do not measure MCU activity directly or in isolation from other Ca^2+^ transport mechanisms, and do not permit adequate control over the experimental conditions. When [Ca^2+^]_cyto_ is in the high nM range (around the resting levels), the MCU-mediated Ca^2+^ uptake is very slow and exists in an equilibrium with the mitochondrial Ca^2+^ efflux mechanisms (*40*). Thus, any measured changes in [Ca^2+^]_mito_ cannot be assigned to MCU exclusively. When [Ca^2+^]_cyto_ is elevated into the μM range, the MCU activity becomes high and overwhelms not only the Ca^2+^ efflux machinery but also the electron transport chain, resulting in a decreased driving force for Ca^2+^ and underestimation of the MCU activity (*3, 4*). Because of all these technical limitations, measuring the effect of MICU knockouts on *I*_Na_ at nM [Ca^2+^]_cyto_ and on *I*_Ca_ at μM [Ca^2+^]_cyto_ with direct patch-clamp electrophysiology is the only way to reliably study the MICU function.

Here we demonstrate that the MCU complex is constitutively active and has no intrinsic [Ca^2+^]_cyto_ threshold. A recent report also suggested no apparent [Ca^2+^]_cyto_ threshold for MCU in heart and skeletal muscle (*66*). Thus, the [Ca^2+^]_cyto_ threshold for elevation of [Ca^2+^]_mito_ is simply determined by the equilibrium between the MCU-dependent Ca^2+^ uptake and the mitochondrial Ca^2+^ efflux mechanisms. Such a simple equilibrium-based [Ca^2+^]_cyto_ threshold for [Ca^2+^]_mito_ elevation was proposed previously and was termed the “set point” (*40*).

Assuming that the *K*_d_ for Ca^2+^ binding to MICU EF hands is ∼600 nM (*50*), MICUs would start potentiating the MCU complex activity already in the high nanomolar range of [Ca^2+^]_cyto_, around the resting levels. Thus, MICUs should help MCU to overcome the mitochondrial Ca^2+^ efflux machinery and decrease the set point.

It is therefore paradoxical that the optical studies report not an increase but a decrease in “threshold” for mitochondrial Ca^2+^ uptake in *MICU1*^*-/-*^. However, it simply illustrates that the results of the optical experiments should be interpreted with a caution not only at the level of MCU but also at the level of the whole organelle. The set point for mitochondrial Ca^2+^ accumulation is affected not only by the MCU-mediated uptake but also by mitochondrial Ca^2+^ efflux and numerous other factors such as Δψ, matrix pH, permeability of the outer mitochondrial membrane, and mitochondria-ER interface, to mention the most obvious. These factors can also change and overcompensate for the reduced MCU activity in *MICU1*^*-/-*^, resulting in a decreased set point. In contrast to the *MICU1*^*-/-*^, such overcompensation, however, cannot correct the phenotype of MCU and EMRE knockouts because the mitochondrial Ca^2+^ uptake is completely eliminated. To further illustrate these points, in species other than mammals, where the compensatory mechanisms induced by MICU1 knockout may be different, the [Ca^2+^]_cyto_ “threshold” for [Ca^2+^]_mito_ elevation is affected in a different way, although the composition of the MCU complex (including EMRE and MICU1) is similar to mammals. Specifically, in *Trypanosoma cruzi*, MICU1 knockout causes an increase in the Ca^2+^ uptake “threshold” and a marked decrease in Ca^2+^ uptake capacity at all [Ca^2+^]_cyto_ (*67*). The possibility that MICU proteins have other functions (*68*) beyond being a part of the MCU complex can further complicate the interpretation of the *MICU1*^*-/-*^ phenotype as assessed by optical methods. In *Drosophila*, a lethal phenotype of MICU1 knockout was not rescued when combined with either MCU or EMRE knockouts (the MCU and EMRE knockouts themselves had mild phenotypes), suggesting functions for MICU proteins beyond the MCU complex (*68*). Thus, the results obtained with optical methods must be interpreted with due consideration to direct electrophysiological and structural data on the MCU complex. Otherwise, not only the properties of the MCU complex, but also mitochondrial Ca^2+^ homeostasis in general will be misunderstood.

In summary, we demonstrate that MICUs are Ca^2+^-dependent MCU potentiators. They are likely to exert their potentiating effect over a range of [Ca^2+^]_cyto_ from resting to high micromolar. By doing so, MICUs can control both the [Ca^2+^]_cyto_ set point for [Ca^2+^]_mito_ elevation and the maximum [Ca^2+^]_mito_ reached during intracellular Ca^2+^ signaling. Importantly, the potentiation of MCU by MICUs could help to reduce the number of MCU channels required for adequate Ca^2+^-dependent stimulation of mitochondrial ATP production. Without MICUs, the number of MCU channels per mitochondrion would have to be ∼2 times higher, which would also increase futile Ca^2+^ cycling at resting [Ca^2+^]_cyto_.

## ACKNOWLEDGEMENTS

We thank Drs. Katsuyoshi Mihara (Kyushu University, Japan) and David C. Chan (Caltech, USA) for sending us *Drp1*^*-/-*^ MEFs, and Dr. Toren Finkel (University of Pittsburgh, USA) for sending the MEFs with intact Drp1 (*WT* and *MICU1*^*-/-*^ MEFs). We thank the Nikon Microscopy Core (DeLaine Larsen, Kari Herrington) and Lab for Cell Analysis (Sarah Elms) at UCSF for help with use of microscopy and FACS equipment. We thank all members of the Y.K. lab for helpful discussions;

## Author contributions

V.G. and Y.K. conceived the project and designed all experiments. V.G. performed all experiments. V.G. and T.U. performed Western blot experiments. J.S. consulted on Ca^2+^ imaging experiments. I.P. helped with the analysis of Ca^2+^ imaging experiments. L.S.M. consulted on single-channel analysis using QuB. V.G and Y.K. discussed the results and wrote the manuscript. All authors commented on the manuscript;

## Data and materials availability

All data is available in the main text or the supplementary materials. Further information and requests for reagents may be directed to the lead contact Yuriy Kirichok.

## SUPPLEMENTARY MATERIALS

Materials and Methods Figure S1 – S10 Tables S1 – S2 References

## Supplementary Materials

### MATERIALS AND METHODS

#### Cell culture and recombinant gene expression

All mouse embryonic fibroblast (MEF) cells with (*32*) or without Drp1 (*45*), and all knockout clones were grown in low glucose (5.6 mM) Dulbecco’s modified Eagle’s medium (DMEM) supplemented with 10% FBS, 100 U/ml penicillin, and 100 U/ml streptomycin at 37°C, 5% CO_2_. Cells were maintained by splitting every 48-72 hours at a ratio of 1:5 to 1:10.

We used third generation lentiviral (bi-cistronic) vectors containing the ORF for gene of interest with or without a selection marker (EGFP, mCherry or puromycin, Supplementary Fig. 1 and 2). The vectors were generated by VectorBuilder, Inc. (Chicago, IL, USA), and their sequences were confirmed independently by the company and by us. Recombinant cDNA expressing cells were enriched using multiple rounds of FACS or antibiotic selection. In some cases, EGFP was targeted to mitochondria (using a mitochondrial targeting sequence from COX8) to identify mitoplasts expressing the recombinant protein of interest during patch clamp experiments.

#### Quantitative Real-Time PCR Analysis

qPCR was performed by Syd Labs (Natick, MA, USA). Total RNA was isolated from cells using the RNAeasy Minikit (QIAGEN), and reverse transcribed using the First Strand cDNA Synthesis Kit (Syd Labs). qPCR reactions were performed with the following gene-specific primers (generated by Integrated DNA Technologies):

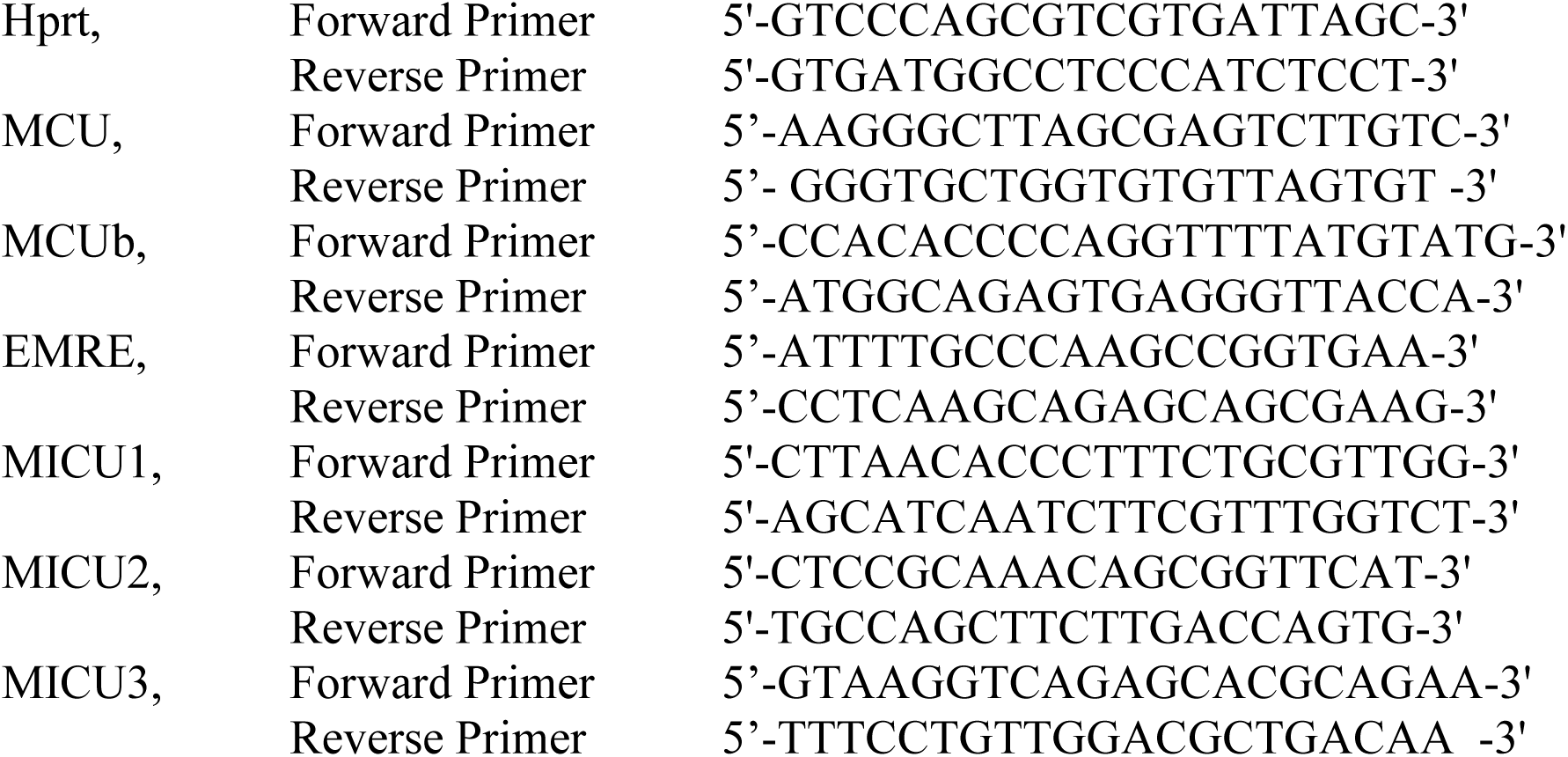

cDNA (100 ng, calculated from initial RNA) samples were pre-amplified for 12 cycles using ABsolute qPCR SYBR Green Low ROX Mix (ThermoFisher). qPCR reactions were performed using an Agilent MX3000 (Fluidigm) with 40 cycles of amplification (15 s at 95°C, 5 s at 70°C, and 60 s at 60°C). Ct values were calculated by the Real-Time PCR Analysis Software (Fluidigm). Relative gene expression was determined by the ΔCt method. Hprt was selected as the reference gene.

#### Generation of knockout cell lines by the CRISPR/Cas9 method

Knockout MEF cell lines were generated using the CRISPR/Cas9 method (*69*). All knockouts (except the *MCU*^*-/-*^ line) were generated by Alstem LLC (Richmond, CA, USA). Either one sgRNA or a pair of two adjacent sgRNAs were used to create a point indel or a truncate indel, respectively (Fig. S1).

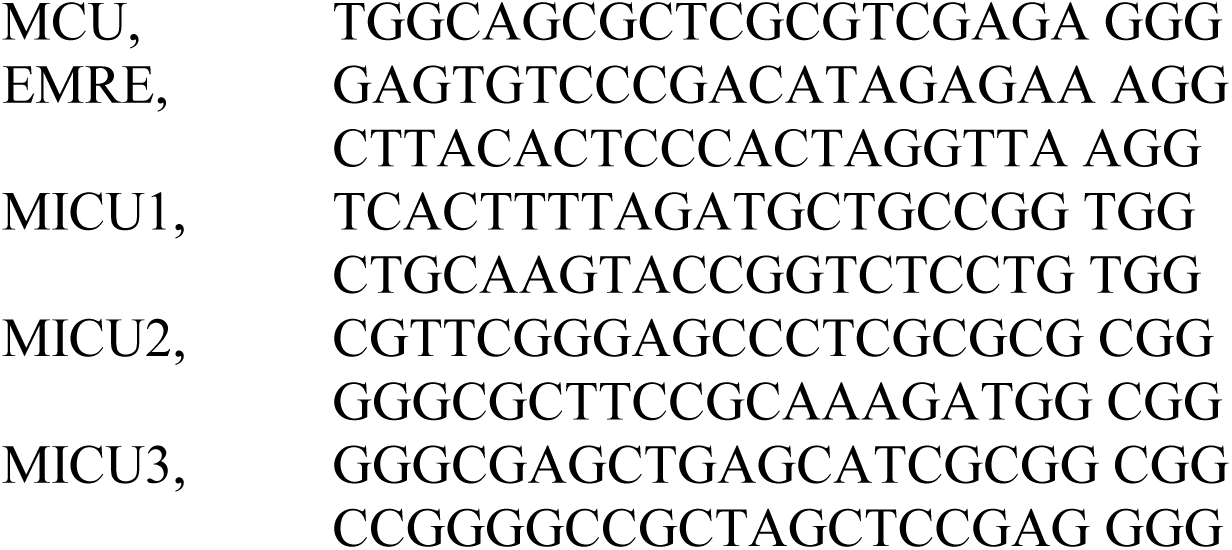

MEFs were transfected with the Cas9 gRNA vector (Addgene: PX459) via electroporation (Invitrogen Neon transfection system) using the following parameters: 1×10^6^ cells and 1 μg of two different gRNA-Cas9 plasmids. Puromycin was used for enrichment of transfected cells, and serial dilution was performed to select single-cell clones. A stable homozygous knockout cell line was confirmed by PCR amplification of the targeted region, cloning into a pUC19 vector, and sequencing showing that either a frameshift or large deletion had occurred in the targeted region of the gene (Fig. S1). All knockout clones were further validated by Western blotting (Fig. S2). The primers used for amplification of genomic sites and cloning into pUC19 sequencing vector were as follows:

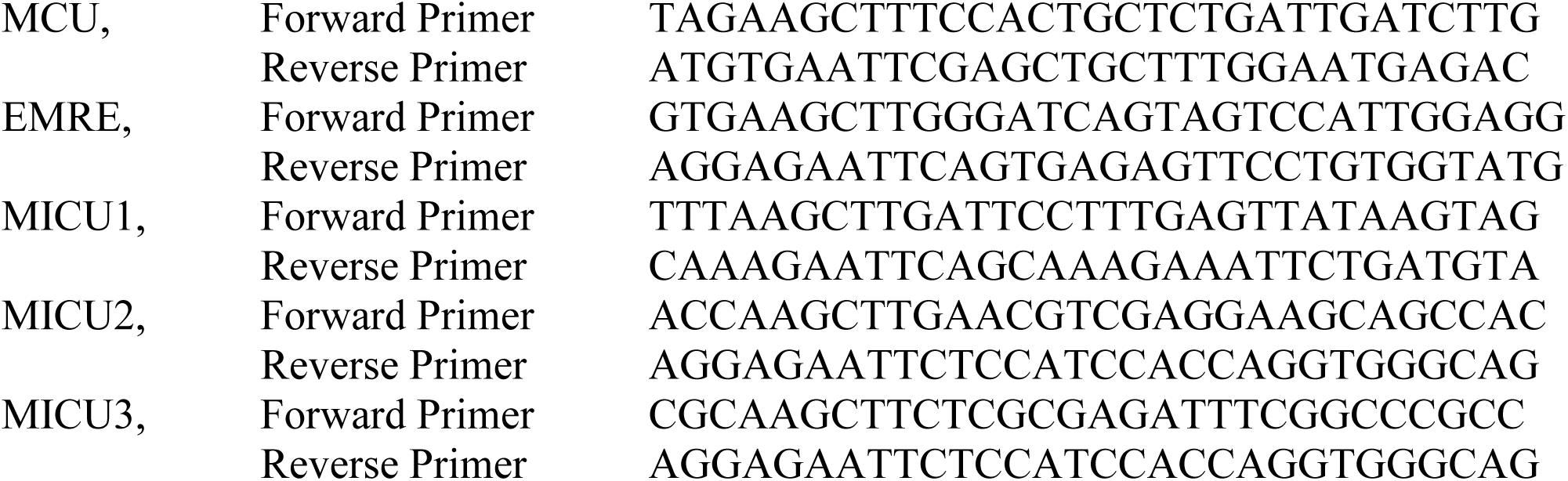

#### Isolation of mitochondria and mitoplasts from MEFs

Mitoplasts were isolated from MEFs using methodology previously described (*41*). Briefly, MEFs were homogenized in ice-cold medium (Initial medium) containing 250 mM sucrose, 10 mM HEPES, 1 mM EGTA, and 0.1% bovine serum albumin (BSA) (pH adjusted to 7.2 with Trizma® base) using a glass grinder with six slow strokes of a Teflon pestle rotating at 280 rpm. The homogenate was centrifuged at 700× g for 10 min to create a pellet of nuclei and unbroken cells. The first nuclear pellet was resuspended in the fresh Initial medium and homogenized again to increase the mitochondrial yield. Mitochondria were collected by centrifugation of the supernatant at 8,500× g for 10 min.

Mitoplasts were produced from mitochondria using a French press. Mitochondria were suspended in a hypertonic solution containing 140 mM sucrose, 440 mM D-mannitol, 5 mM HEPES, and 1 mM EGTA (pH adjusted to 7.2 with Trizma® base) and then subjected to a French press at 1,200–2,000 psi to rupture the outer membrane. Mitoplasts were pelleted at 10,500× g for 15 min and resuspended for storage in 0.5–1 ml of solution containing 750 mM KCl, 100 mM HEPES, and 1 mM EGTA (pH adjusted to 7.2 with Trizma® base). Mitoplasts prepared and stored with this method contained the same amount of auxiliary MICU1 and MICU2 subunits as compared to intact mitochondria (Fig. S4a, see the co-immunoprecipitation section below).

Mitochondria and mitoplasts were prepared at 0–4 °C and stored on ice for up to 5 h. Immediately before the electrophysiological experiments, 15–50 μl of the mitoplast suspension was added to 500 μl solution containing 150 mM KCl, 10 mM HEPES, and 1 mM EGTA (pH adjusted to 7.0 with Trizma® base) plating on 5-mm coverslips pretreated with 0.1% gelatin to reduce mitoplast adhesion.

#### Patch-clamp recording

Whole mitoplast currents were measured as described previously (*41*). Gigaohm seals with mitoplasts were formed in the bath solution containing 150 mM KCl, 10 mM HEPES and 1 mM EGTA, pH 7.2 (adjusted with KOH). Voltage steps of 350–500 mV for 2-8 ms were applied to rupture the IMM and obtain the whole-mitoplast configuration. Typically, pipettes had resistances of 20–40 MΩ, and the access resistance was 35–65 MΩ. The membrane capacitances of mitoplasts range from 0.2 – 0.6 pF.

All indicated voltages are on the matrix side of the IMM (pipette solution), relative to the cytosolic side (bath solution, Fig. S3) (*41*). Currents were normally induced by a voltage ramp from −160 mV to +80 mV (interval between pulses was 5 s) to cover all physiological voltages across the IMM, but other voltage protocols were also used as indicated in the figures. All whole-IMM recordings were performed under continuous perfusion of the bath solution. Currents were normalized per membrane capacitance to obtain current densities (pA/pF). Currents flowing into mitochondria are shown as negative, while those flowing out are positive. Membrane capacitance transients observed upon application of voltage steps were removed from current traces.

Typically, pipettes were filled with one of the following three solutions (*41*) (tonicity was adjusted to ∼350 mmol/kg with sucrose.):

*Solution A* was used to measure *I*_Ca_ and contained: 110 mM Na-gluconate, 40 mM HEPES, 10 mM EGTA and 2 mM MgCl_2_ (pH 7.0 with NaOH)

*Solution B* was used to measure *I*_Na_ or *I*_Mn_ and contained: 110 Na-gluconate, 40 HEPES, 1 EGTA, 5 EDTA and 2 mM NaCl (pH 7.0 with Tris base).

*Solution C* was used to measure outward *I*_Ca_ (the MCU rectification experiments) and contained: 130 mM tetramethylammonium hydroxide (TMA), 100 mM HEPES and 2 mM CaCl_2_ (pH 7.0 with D-gluconic acid)

To measure whole-mitoplast *I*_Ca_, the bath solution was formulated to contain only 150 mM HEPES (pH 7.0 with Tris base, tonicity ∼300 mmol/kg with sucrose) and different dilutions of CaCl_2_ from a 1 M stock (Sigma)(*9*). The control solution contained: 150 mM HEPES, 80 mM sucrose and 1 mM EGTA (pH 7.0 with Tris base, tonicity ∼300 mmol/kg with sucrose). The bath solution used for measuring *I*_Na_ contained: 110 mM Na-gluconate, 40 mM HEPES, 1 mM EGTA and 5 mM EDTA (pH 7.0 with Tris base, tonicity ∼300 mmol/kg with sucrose). The bath solution for measuring inhibition of *I*_Na_ by cytosolic Ca^2+^ contained: 110 mM Na-gluconate, 40 mM HEPES and 10 mM EDTA (pH 8.0 with Tris, tonicity ∼380 mmol/kg with sucrose)) and varying amounts of CaCl_2_ were added to the bath solution to achieve the free [Ca^2+^] calculated using the MaxChelator program (C. Patton, Stanford University).

A rapid exchange of [Ca^2+^]_cyto_ from virtual zero (control solution) to 1 mM was achieved using a commercially available piezo-driven, fast solution exchange system (Warner Instruments, SF-77B perfusion fast step system). It was interfaced with our pClamp acquisition software in order to precisely time the steps during solution change. The timing (τ ∼0.4 ms) for solution exchange was judged by the current changes because of a junction potential difference using solutions with different ionic strengths.

Currents were recorded using an Axopatch 200B amplifier (Molecular Devices). Data acquisition and analyses were performed using PClamp 10 (Molecular Devices) and Origin 9.6 (OriginLab). All data were acquired at 10 kHz and filtered at 1 kHz.

#### Single-channel recordings and analysis

All single-channel data were acquired from inside–out patches excised from isolated mitoplasts(*9*). Patches were excised in a bath solution containing 150 mM KCl, 10 mM HEPES and 1 mM EGTA, pH 7.2 (adjusted with KOH). Recordings were performed under symmetrical conditions (the same bath and pipette solutions): 105 mM CaCl_2_ and 40 mM HEPES, pH 7.0 with Tris base. Signals were sampled at 50 kHz and low-pass filtered at 1 kHz. Fire-polished, borosilicate pipettes (Sutter QF-150-75) coated with Silguard (Dow Corning Corp., Midland, MI) and having a tip resistance of 50–70 MΩ were used for low noise recordings.

To characterize the single-channel conductance and subconductance levels and their occupancy probabilities, we used the MLab version of the QuB software, freely available from the Milescu lab at: https://milesculabs.biology.missouri.edu/QuB_Downloads.html. The data were first resampled at 2.5 kHz and then were idealized with the Baum-Welch and Viterbi algorithms, as implemented in QuB, which classify each point in the data to a conductance level and produce estimates of current amplitudes and occupancy probabilities. The time-averaged single-channel current can be calculated as the product between occupancy probability and current amplitude, summated over all conductance levels (main open state and substates).

#### Time-lapse Ca^2+^ imaging

For imaging experiments, MEFs were plated on collagen type-I-coated glass-bottom 35 mm dishes (P35G-1.5-14-C, Matek), 48–72 h before imaging. Cells were imaged at the interval of 3 s on a Nikon Ti-E microscope using a 40× objective (NA 1.30, oil, CFI Plan Fluor, Nikon), Lambda 421 LED light source (Sutter) and ORCA Flash 4.0 CMOS camera (Hamamatsu Photonics) at room temperature (25°C). The following excitation/emission filter settings were used: 340±13 nm/525±25 and 389±19 nm/510±40 for cytosolic Ca^2+^ imaging using fura-2 (*K*_d_=224 nM) and 480±40 nm nm/525±15 nm for mitochondrially targeted *cepia2* (*CEPIA2mt, K*_d_=160 nM (*46*), cloned into a lentiviral vector). Cells were loaded with 3 μM fura-2 AM (Life Tech., USA) in DMEM/FBS at room temperature for 30 min. After three washes with physiological salt solution (PSS) containing (in mM) 150 NaCl, 4 KCl, 2 CaCl_2_, 1 MgCl_2_, 5.6 glucose and 25 HEPES (pH 7.4), each dish was placed on the stage for imaging. Imaging was performed in PSS within 1 h of dye staining. Baseline fluorescence was taken for 1–2 min after which thapsigargin (Tg) (final [Tg] = 300 nM) was added while imaging was continued for another 10–15 min.

##### Fura-2 Calibration

Baseline measurements were taken, and cells were incubated in PSS (No CaCl_2_) containing 3 mM EGTA, 1 μM ionomycin and 1 μM Tg for 5–10 min. After 2–3 washes with PSS (No CaCl_2_) containing 0.3 mM EGTA, cells were imaged for 5 min (average of last 10 frames was used for calculation) to obtain the R_min_ and F_380max_ values. Finally, PSS containing 10 mM CaCl_2_ (no EGTA), 1 μM ionomycin and 1 μM Tg was added and cells were imaged for 10 min. After the signal reached saturation (∼3 min), the average value from 10 frames was used to calculate R_max_ and F_380min_ values. Using these obtained values, the fura-2 ratio was calibrated by the following equation (*70*):

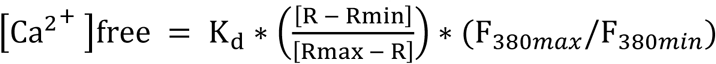

All image analyses were done with ImageJ (NIH). Briefly, mitochondrial and cytosolic regions were manually determined for each cell. The average fluorescence intensity in the regions was measured and the background intensity was subtracted. For analysis of the *cepia2* signal, we normalized the fluorescence intensity by the baseline fluorescence. For analysis of the fura-2 signal, we calculated the fluorescence ratio (F_340_/F_380_ for fura-2).

The time point for increase in mitochondrial [Ca^2+^] (upstroke) was detected using a script written in Python (https://github.com/ishanparanjpe/upstroke) and manually checked afterwards. Briefly, the fluorescence signal was smoothed by applying a second–order zero phase digital Butterworth filter with an optimal cutoff frequency as previously described (*71*). From the smoothed signal, the upstroke frame was defined as the earliest point between the baseline and signal peak that was greater than 80% of the maximal time derivative. The time-point for change in mitochondrial signal was time-matched with the fura-2 reading to determine the threshold [Ca_2+_]_cyto_.

#### Co-immunoprecipitation

Mitochondria or mitoplasts were isolated from MEFs deficient in the MCU subunit but stably expressing Flag-tagged MCU. Mitochondrial fraction from wild type cells (without MCU-FLAG) was used as negative control. Isolated mitoplasts (but not mitochondria) were incubated in 750 mM KCl for 30 min before solubilization. Briefly, 300 μg of protein lysate was solubilized with 500 μl of lysis buffer (50 mM HEPES pH 7.4, 150 mM NaCl, 1 mM EGTA, 0.2% DDM and Halt protease inhibitor cocktail [Thermo Fisher]) for 30 min at 4°C. Lysates were cleared by spinning at 20,000× g for 10 min at 4°C. Cleared lysates were incubated with anti-Flag M2 affinity gel (Sigma A2220) for 2 h at 4°C. Immunoprecipitates were washed with 1 ml of lysis buffer three times and boiled in 20 μl of Laemmli buffer (without β-mercaptoethanol). One-third of the immunoprecipitate was loaded onto a 4–20% gradient SDS-PAGE gel for detection of the indicated proteins by Western blotting. Flow-through fraction was also collected and analyzed in the same gel.

#### Immunoblots

For Western blot analysis, MEFs or isolated mitochondria/mitoplasts were lysed in radioimmunoprecipitation assay (RIPA) buffer (1% IGEPAL^®^, 0.1% sodium dodecyl sulfate, 0.5% sodium deoxycholate, 150 mM NaCl, 1 mM EDTA, 50 mM Tris-HCl (pH 7.4) and a cocktail of proteases inhibitors). Lysates were resolved by SDS-PAGE; transferred to PVDF membrane (Millipore); and probed with anti-MCU (Sigma, HPA016480, 1:2,000), anti-EMRE (Santa Cruz, sc-86337, 1:200), anti-HSP60 (Santa Cruz, sc-1052, 1:3,000), anti-VDAC (Abcam, ab15895, 1:2,000), anti-MICU1 (Cell Signaling Technology, 12524S, 1:2,000), anti-MICU2 (Bethyl, A300-BL19212, 1:500), anti-MICU3 (Sigma, HPA024779, 1:1,000) and anti-TOM20 (Santa Cruz, sc-11415, 1:2,000). Anti-MICU1 antibody produced a non-specific band near its monomeric molecular weight (∼50 kDa), so samples were prepared in Laemmli buffer without β-mercaptoethanol to detect MICU1 homo-or hetero-dimers (∼100 kDa).

#### Statistical analysis

Data are presented as mean ± standard error of the mean (SEM), as specified in the figure legend. Statistical analysis was completed in Excel or Origin 9.6. All experiments were performed in triplicate or more. Statistical significance at an exact *p-value* was determined with the methods as indicated in the corresponding figure legends.

## SUPPLEMENTARY FIGURES

**Fig. S1.**
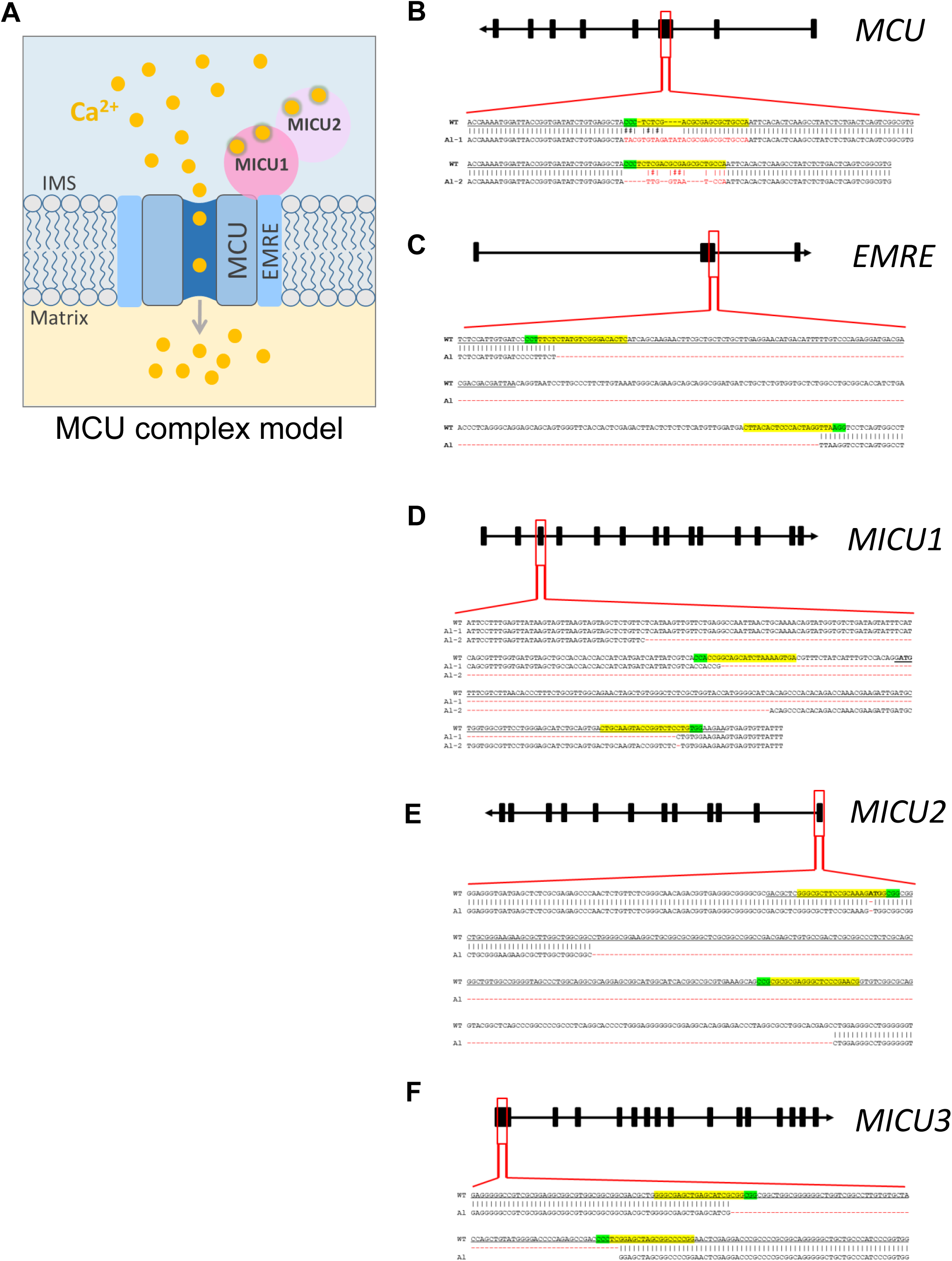
Generation of knockouts for various MCU complex subunits. (**A**) A schematic arrangement of various subunits in the MCU complex. Four MCU and four EMRE subunits form the pore of the MCU complex (only two MCU and two EMRE subunits are shown for simplicity). EMRE also tethers MICU1 subunit to the pore on the cytosolic side of the IMM (i.e., in the mitochondrial intermembrane space, IMS). MICU1 forms homodimers or hetero-dimerizes with MICU2 or MICU3 (not shown). Each MICU subunit has two EF hands that bind cytosolic Ca^2+^. (**B** to **F**) CRISPR-mediated indels in various MCU subunit genes and the resulting mutant alleles. The CRISPR binding sites (for sgRNA) are highlighted in *yellow*, and their PAM sequences are highlighted in *green*. The translational initiation codon (ATG) is shown in *bold* where applicable. (B) Overview of the *MCU* gene and indels in the knockout. A sgRNA was used to target exon 3. The sequence of targeted region in MCU gene is shown; exon 3 is underlined. Targeted sequencing indicates frame-shift indels (*red*) in both alleles (*Al-*1 and *Al-*2). (C) Overview of the *EMRE* gene and truncated region in the knockout. Two sgRNAs were used for CRISPR-Cas9–mediated deletion in the exon-2 (*underlined*) and the flanking region. Targeted sequencing indicates same 259-bp deletion (*red*) in both alleles. (D) Overview of the *MICU1* gene and truncated region in the knockout. Two sgRNAs were used for CRISPR-Cas9– mediated deletion in the exon-3 (*underlined*) and the flanking region. Targeted sequencing indicates that almost all of exon-3 is deleted along with a portion of the flanking region (*red*) in both alleles (*Al-*1 and A*l-*2). (E) Overview of the *MICU2* gene and truncated region in the knockout. Two sgRNAs were used for CRISPR-Cas9–mediated deletion in the exon-1 (*underlined*) and the flanking region. Targeted sequencing indicates that almost all of exon-1 is deleted (*red*) in both alleles. (F) Overview of the *MICU3* gene and truncated region in the knockout. Two sgRNAs were used for CRISPR-Cas9–mediated deletion in the exon-1 (*underlined*). Targeted sequencing indicates a 73-bp deletion in the expected cut area (*red*) in both alleles.

**Fig. S2.**
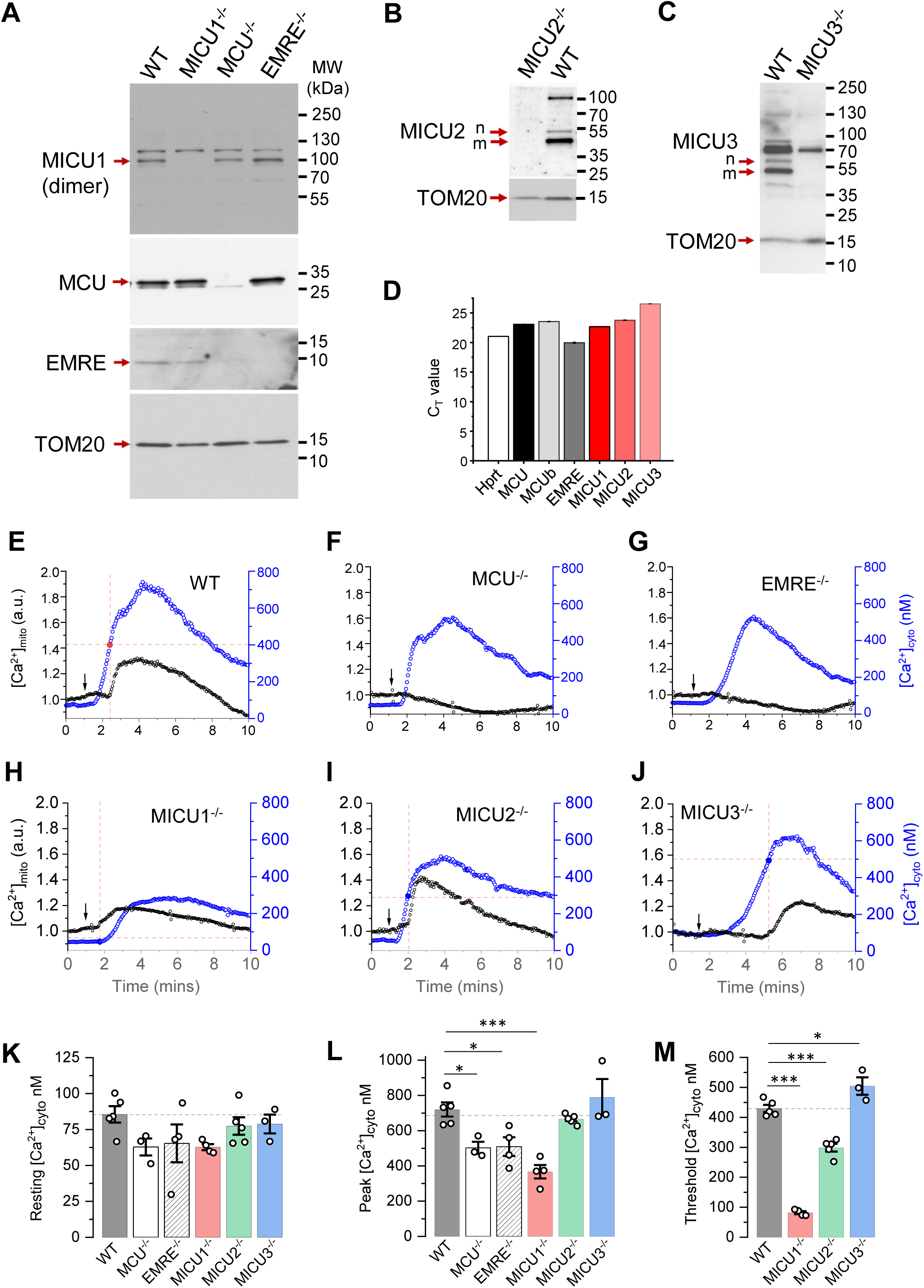
Expression of various MCU complex subunits and [Ca^2+^]_mito_ phenotype in cells deficient for various MCU complex subunits. (**A** to **C**) Western blots show expression of various MCU complex subunits in the respective knockout cells. For MICU1 (**A**), samples were prepared without reducing agent, β-mercaptoethanol. The MICU1 band is near the expected molecular weight (∼100 kDa) for the homo-or hetero-dimer (with MICU2 or 3). Multiple bands were observed with anti-MICU2 (**B**) and anti-MICU3 (**C**) antibodies, which were absent in knockout cell lines. This is likely due to the presence of different oligomeric states of the protein, as well as the mature and nascent (before truncation of the mitochondrial targeting signal) forms of the protein. Arrows mark the mature (m) and nascent (n) proteins near the expected molecular weight. (**D**) PCR showing the mRNA expression of various MCU subunits in *Drp1*^*-/-*^ MEFs. Hprt was used as the reference. (**E** to **J**) Representative [Ca^2+^]_mito_ (*black*, left ordinate) and [Ca^2+^]_cyto_ (*blue*, right ordinate) in an individual cell with *WT* MCU complex, and individual cells with MCU, EMRE, and MICU1−3 knockouts before and after application of 300 nM Tg (arrow). Dashed red lines indicate the [Ca^2+^]_cyto_ at which the [Ca^2+^]_mito_ starts to increase (“[Ca^2+^]_cyto_ threshold”). (**K** to **M**) Resting [Ca^2+^]_cyto_ (K), peak [Ca^2+^]_cyto_ after addition of Tg (L), and [Ca^2+^]_cyto_ threshold for [Ca^2+^]_mito_ elevation (M) in *WT* and indicated knockout cell lines. *WT* (*n* = 5 dishes, total cells = 150); *MCU*^*-/-*^ (*n* = 3 dishes, total cells = 183); *EMRE*^*-/-*^ (*n* = 4 dishes, total cells = 187); *MICU1*^*-/-*^ (*n* = 4 dishes, total cells = 196); *MICU2*^*-/-*^ (*n* = 4 dishes, total cells = 192); and *MICU3*^*-/-*^ (*n* = 3 dishes, total cells = 115). Mean ± SEM; one-way ANOVA with post-hoc Tukey test. \**p*< 0.05; \*\*\**p*< 0.001. Statistics was run on number of dishes.

**Fig. S3.**
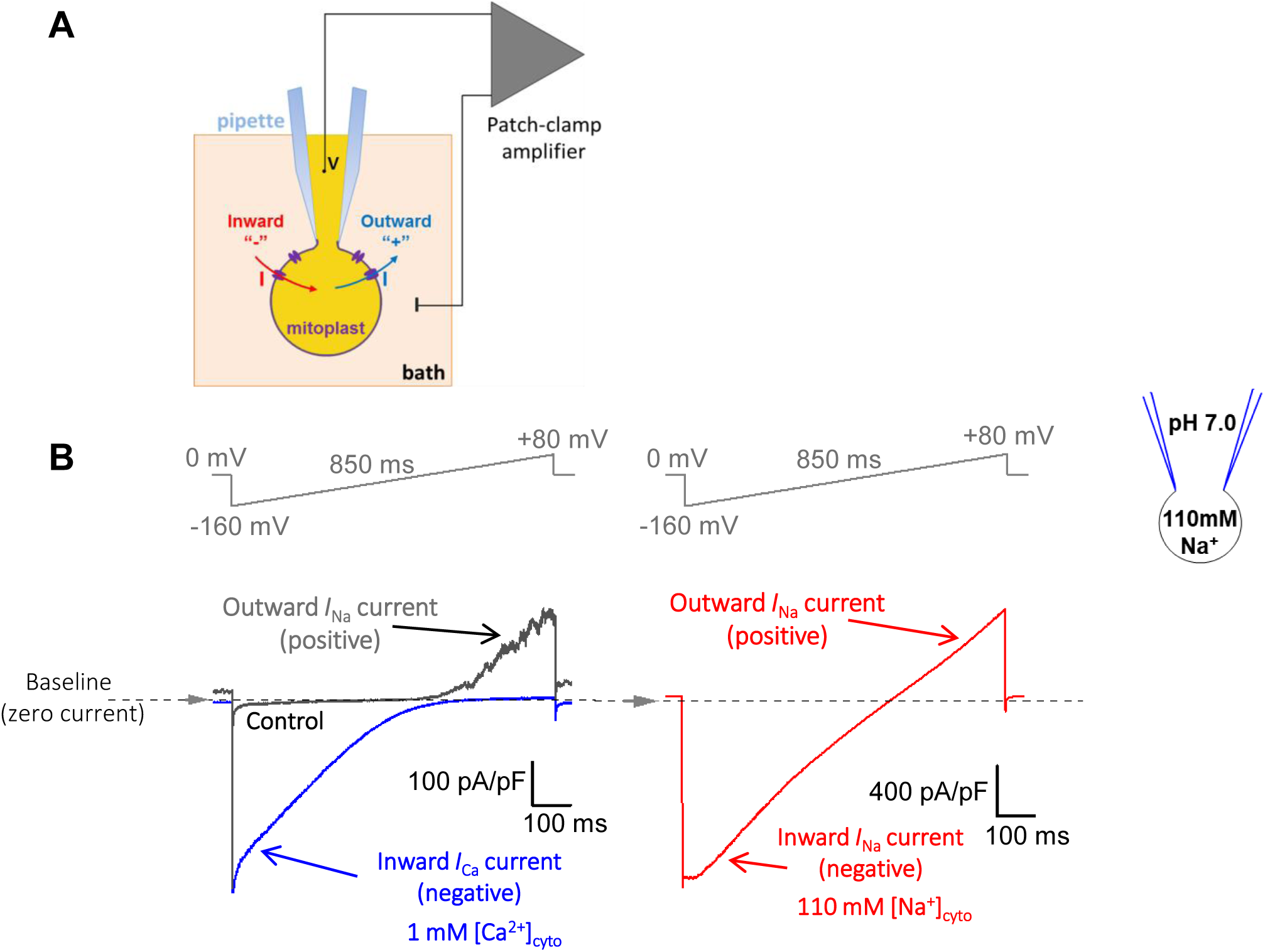
Recording MCU currents across the whole IMM. (**A**) Diagram of patch-clamp recording from a vesicle of the whole IMM (mitoplast). After formation of a gigaohm seal between the patch pipette and the mitoplast, the IMM patch under the pipette is broken by applying short pulses of high voltage (200–500 mV, 2–8 ms), sometimes combined with light suction, to gain access into the mitoplast through the pipette. In this configuration, called the “whole-IMM” configuration, the interior of the mitoplast (mitochondrial matrix) is perfused with the pipette solution. The bath is also perfused to control the experimental solution on the cytosolic side of the IMM. The voltage across the IMM is set to the desired value (V), and the currents (I) are measured using the patch-clamp amplifier. Directions of currents flowing across the IMM: inward currents (flowing into the mitoplast) are negative, while outward currents are positive. (**B**) *Left panel:* Example MCU current traces recorded in the whole-IMM configuration. The voltage protocol used to elicit the currents is shown above. All indicated voltages are within the mitochondrial matrix relative to the bath (cytosol). The voltage of the bath solution is defined to be zero. The zero current level is shown by the dashed line and an arrow. The directions of the currents are indicated as negative (inward) and positive (outward). The MCU current in Ca^2+^-free bath solution (control) is shown in *grey*. The outward current in control is mediated by Na^+^ ions permeating through the MCU channel in the Ca^2+^-free conditions (*I*_Na_, pipette solution contains Na^+^). After application of 1 mM Ca^2+^ on the cytosolic face of the IMM (bath), we observe an inward Ca^2+^ current (*I*_Ca_, *blue*) via MCU, while the outward *I*_Na_ is simultaneously inhibited. *Right panel*, When [Ca^2+^]_cyto_ is brought to virtual zero (1 mM EGTA and 5 mM EDTA) under conditions when both bath and pipette solution contain Na^+^, we observe *I*_Na_ via MCU (*red*) in both inward and outward directions. The current amplitude and time calibration bars are indicated. The current amplitude is normalized per membrane capacitance to facilitate comparison of current amplitudes between mitoplasts of different sizes.

**Fig. S4.**
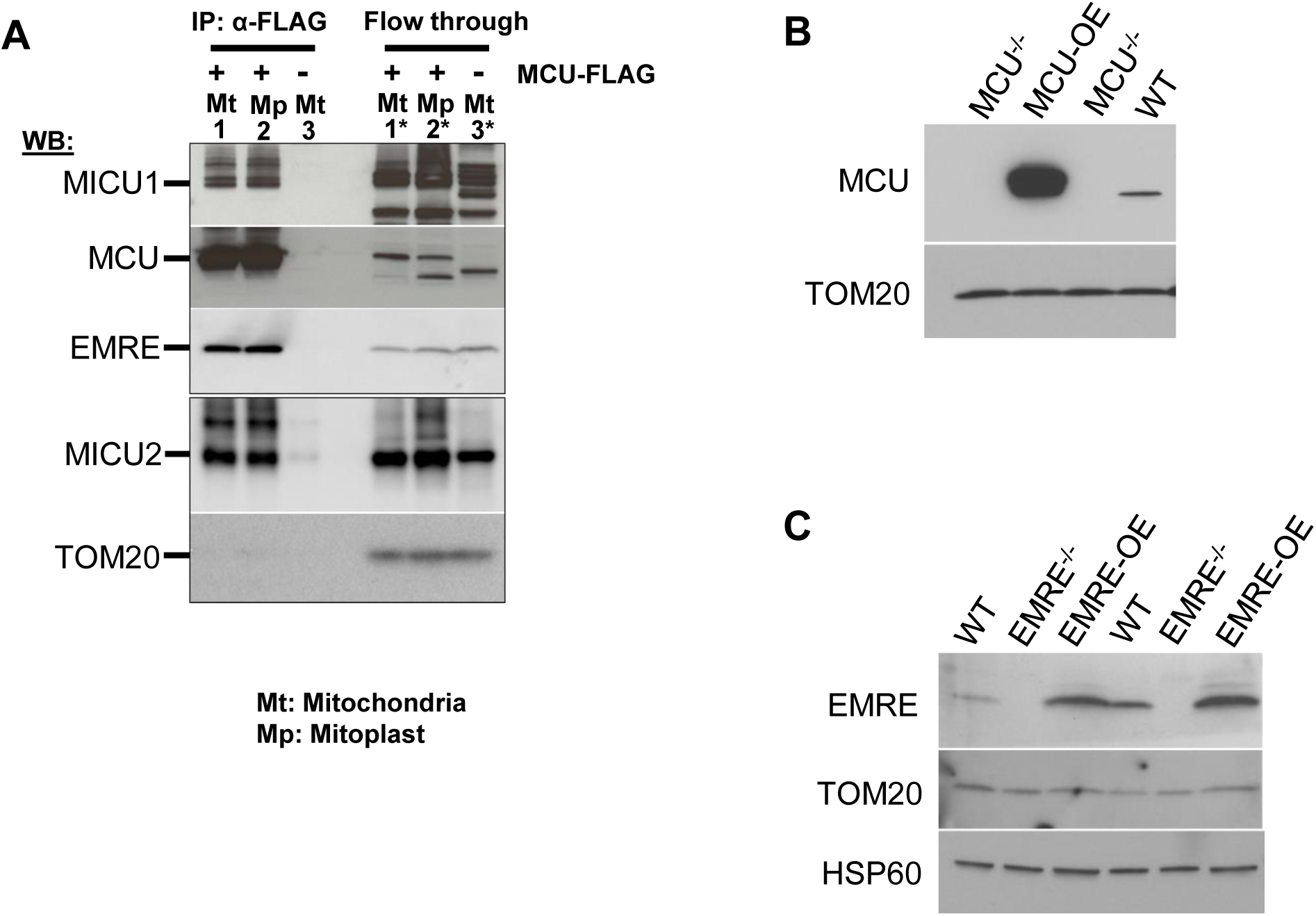
Protein expression of MCU subunits in MEFs and isolated mitoplasts. (**A**) Co-immunoprecipitation of the MCU complex proteins from mitochondrial and mitoplast fractions. Anti-FLAG beads were used to immunoprecipitate MCU-FLAG (expressed in *MCU*^*-/-*^ cells) from mitochondrial and mitoplast fractions. Mitochondria isolated from *WT* cells (No FLAG tag) were used as negative control. Left three lanes are protein-complexes immunoprecipitated with anti-FLAG beads. Lane-1: immunoprecipitate (IP) from MCU-FLAG mitochondrial (Mt) lysate, lane-2: IP from MCU-FLAG mitoplast (Mp) lysate, lane-3: IP from *WT* mitochondrial lysate. Right three lanes correspond to samples from the flow-through fraction after immunoprecipitation. Lane-1*: mitochondrial lysate from MCU-FLAG, lane-2*: mitoplast lysate from MCU-FLAG, lane-3*: mitochondrial lysate from *WT*. Upper (MICU1, MCU and EMRE) and lower (MICU2 and TOM20) boxes are from the same samples run on different gels. (**B**) Western blots of protein lysates from cells with *WT* MCU complex (*WT*), *MCU*^*-/-*^ cells, and *MCU*^*-/-*^ cells overexpressing MCU (MCU-OE) using anti-MCU and anti-TOM20 (the mitochondrial loading control). (**C**) Western blots of protein lysates from *WT* cells, *EMRE*^*-/-*^ cells, and *EMRE*^*-/-*^ cells overexpressing EMRE (EMRE-OE) using anti-EMRE, anti-TOM20 and anti-HSP60 (the mitochondrial loading controls).

**Fig. S5.**
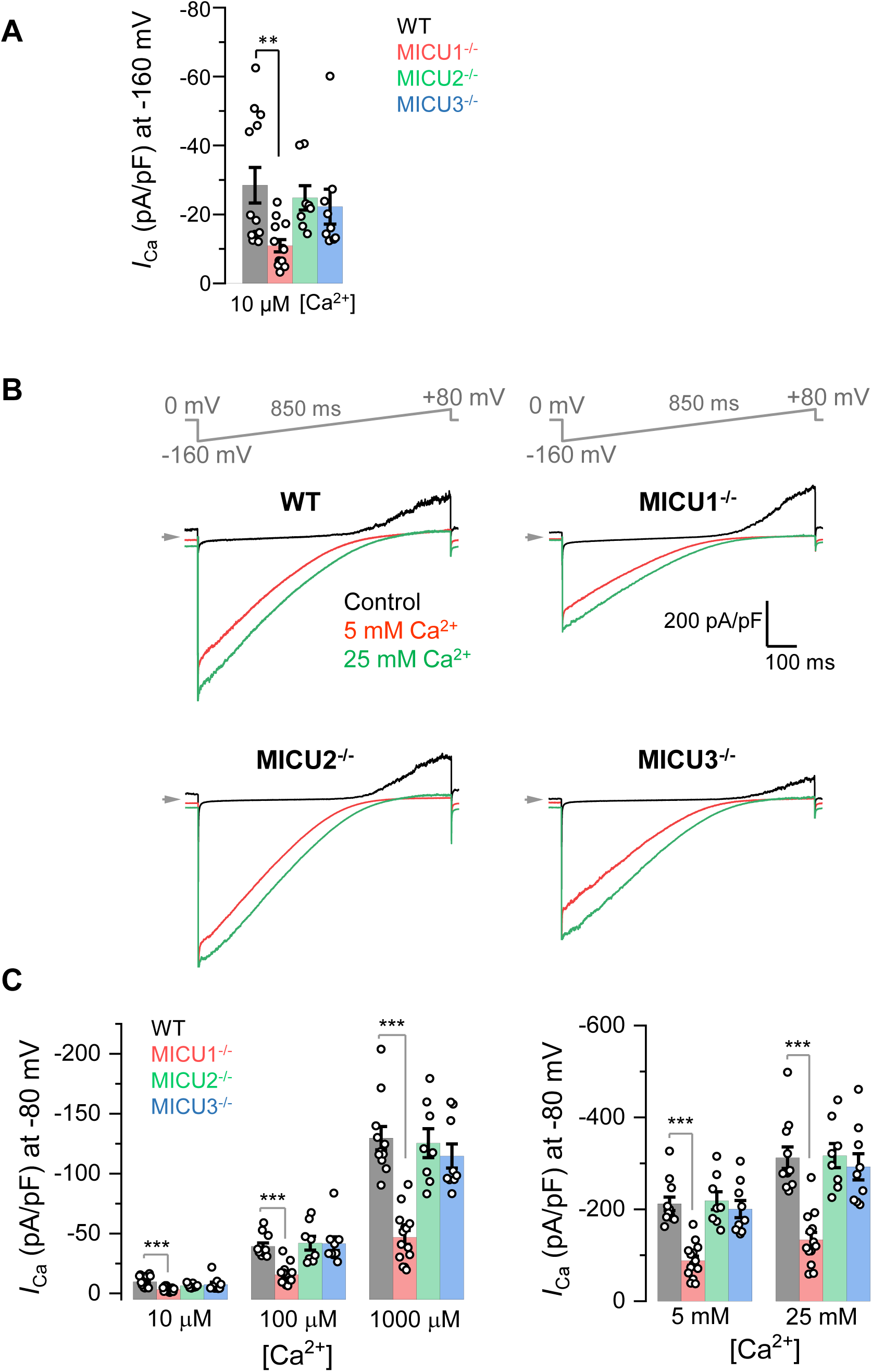
*I*_Ca_ in MICU1−3 knockouts. (**A**) *I*_Ca_ amplitude in *WT* and MICU1-3 knockouts measured at −160 mV using 10 μM [Ca^2+^]_cyto_ with an enlarged Y-axis. Data is same as used in Fig. 1E. Mean ± SEM; one-way ANOVA with post-hoc Tuckey test; \*\**p*< 0.01. (**B**) Representative inward *I*_Ca_ in *WT, MICU1*^*-/-*^, *MICU2*^*-/-*^ and *MICU3*^*-/-*^ mitoplasts exposed to 5 mM, and 25 mM [Ca^2+^]_cyto_. (**C**) *I*_Ca_ amplitude measured at −80 mV in *WT* (*n* = 13), *MICU1*^*-/-*^ (*n* = 14), *MICU2*^*-/-*^ (*n* = 8) and *MICU3*^*-/-*^ (*n* = 9) mitoplasts at 10 μM, 100 μM and 1000 μM [Ca^2+^]_cyto_ (*left*, for *I*_Ca_ traces see Fig. 2A) and at 5 mM and 25 mM [Ca^2+^]_cyto_ (*right*, for *I*_Ca_ traces see Figure 1D). Mean ± SEM; one-way ANOVA with post-hoc Tuckey test; \*\*\**p*< 0.001.

**Fig. S6.**
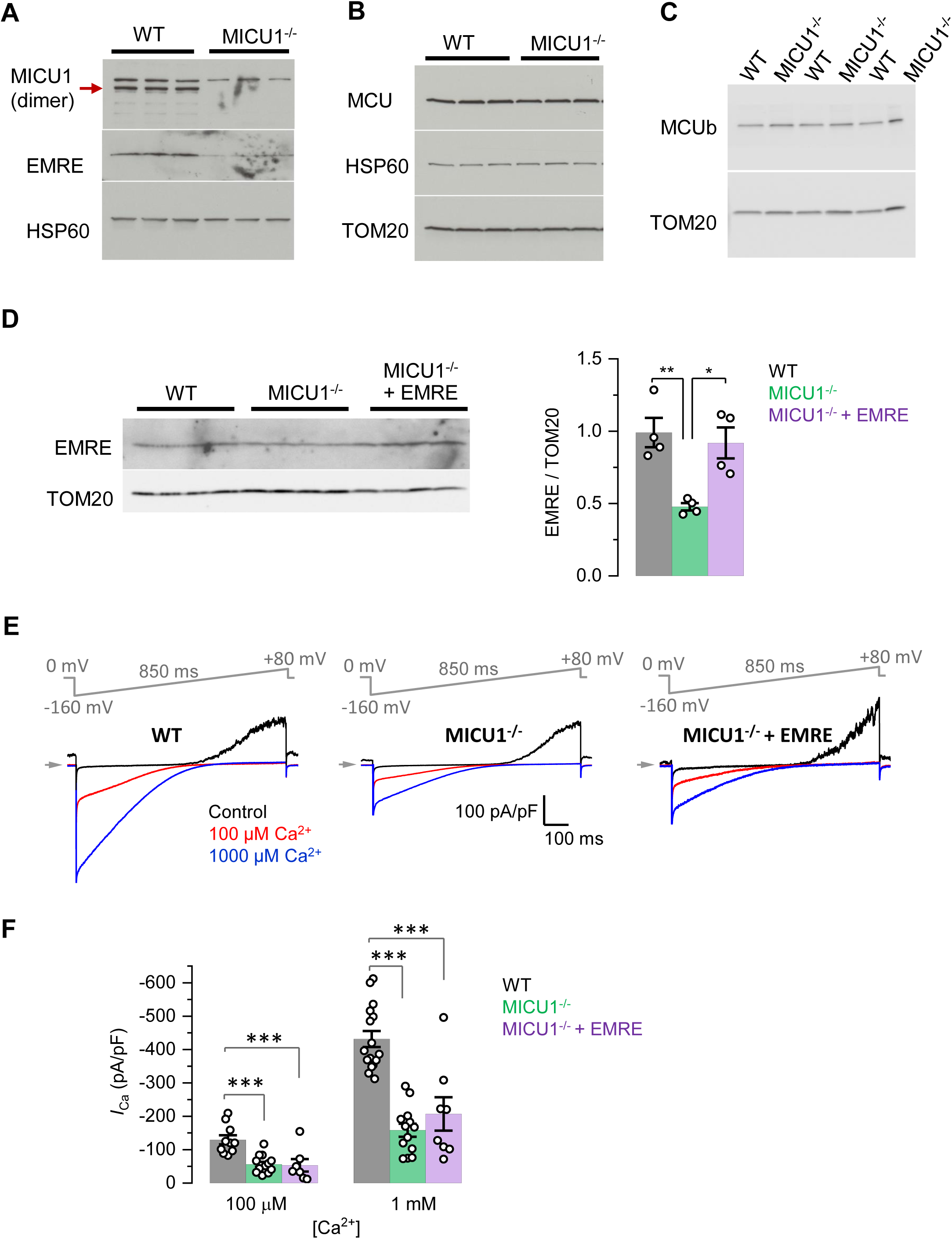
Rescue of EMRE expression in *MICU1*^*-/-*^ does not rescue *I*_Ca_. (**A** to **C**) Western blots showing the expression levels of EMRE (A), MCU (B) and MCUb (C) in cells with *WT* MCU complex and *MICU1*^*-/-*^ (*n* = 3 independent samples each). (**D**) (*Left*) Western blots showing EMRE protein level in *WT* and *MICU1*^*-/-*^ (before and after EMRE overexpression). (*Right*) Graph represents quantification of Western blot (*n* = 4 independent samples each). (**E**) Representative inward *I*_Ca_ in *WT, MICU1*^*-/-*^, and when EMRE was overexpressed in *MICU1*^*-/-*^ (*MICU1*^*-/-*^ + EMRE) upon exposure to 100 μM and 1000 μM [Ca^2+^]_cyto_. (**F**) *I*_Ca_ amplitudes measured at −160 mV in *MICU1*^*-/-*^ overexpressing EMRE (*MICU1*^*-/-*^ + EMRE, *n* = 8) as well as in *MICU1*^*-/-*^ and *WT. WT* and *MICU1*^*-/-*^ data are the same as in Fig. 1E. Mean ± SEM; one-way ANOVA with post-hoc Tuckey test. \**p*< 0.05; \*\**p*< 0.01; \*\*\**p*< 0.001.

**Fig. S7.**
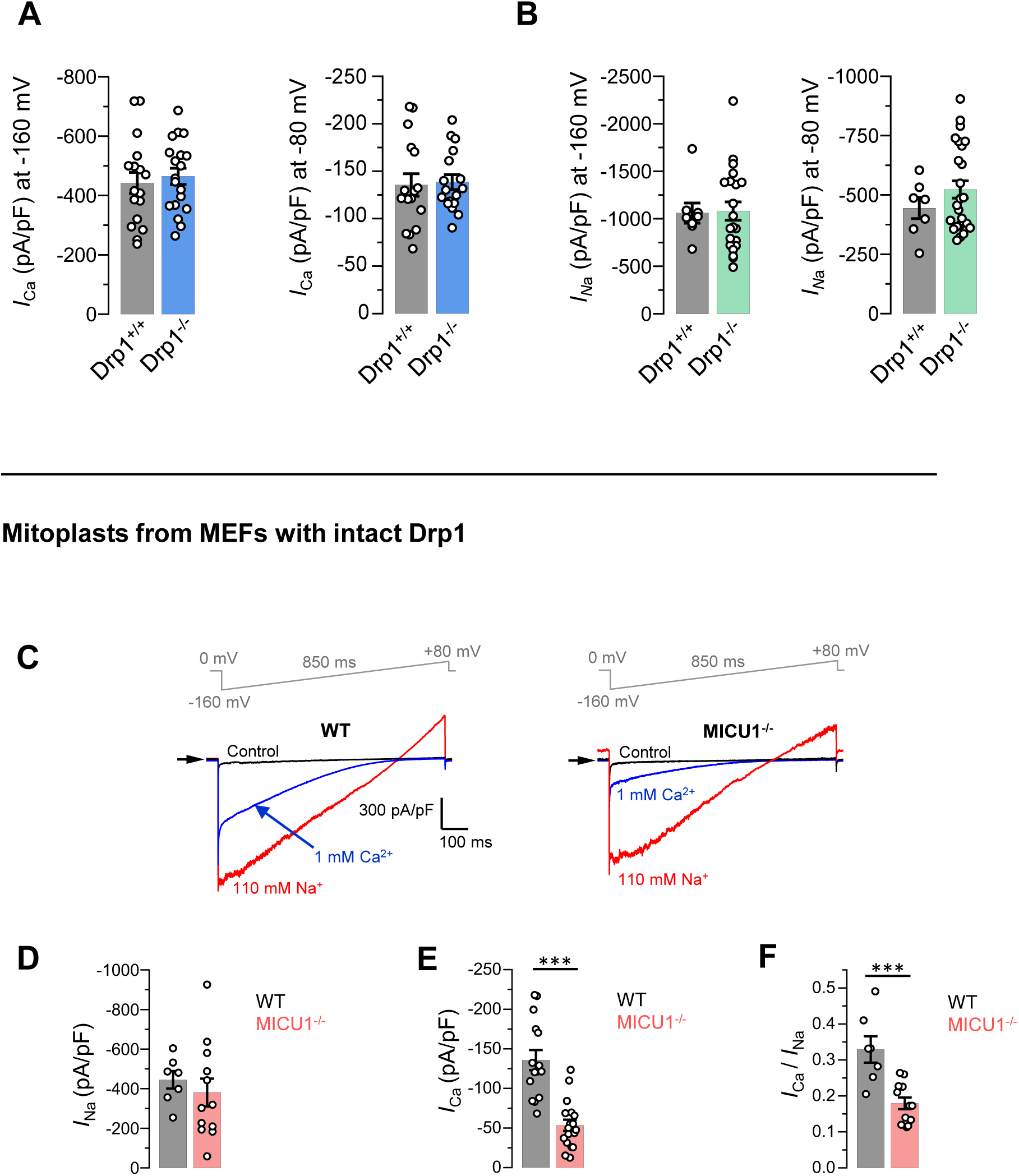
Drp1 does not affect the currents mediated by the MCU complex or their phenotype in *MICU1*^*-/-*^. (**A**) *I*_Ca_ amplitudes at −160 mV (*left*) and −80 mV (*right*) in mitoplasts from MEFs with Drp1 (*Drp*^*+/+*^) and without Drp1 (*Drp1*^*-/-*^). (*n* = 17-19) Mean ± SEM. (**B**) *I*_Na_ amplitudes at −160 mV (*left*) and −80 mV (*right*) in mitoplasts from MEFs with Drp1 (Drp^+/+^, *n* = 8) and without Drp1 (*Drp1*^*-/-*^, *n* = 21). Mean ± SEM. (**C** to **F**) Current phenotypes of *MICU1*^*-/-*^ in mitoplasts isolated from MEFs with an intact Drp1. (C) Representative *I*_Ca_ (*blue*) and *I*_Na_ (*red*) recorded from the *WT* (*n* = 7) and *MICU1*^*-/-*^ (*n* = 12) mitoplasts exposed to 1 mM [Ca^2+^]_cyto_ or 110 mM [Na^+^]_cyto_. Amplitudes of *I*_Na_ (D) and *I*_Ca_ (E) measured at −80 mV. (F) Ratio between *I*_Ca_ and *I*_Na_ measured in the same mitoplast. Mean ± SEM; unpaired t-test, two-tailed; \*\*\**p*< 0.001.

**Fig. S8.**
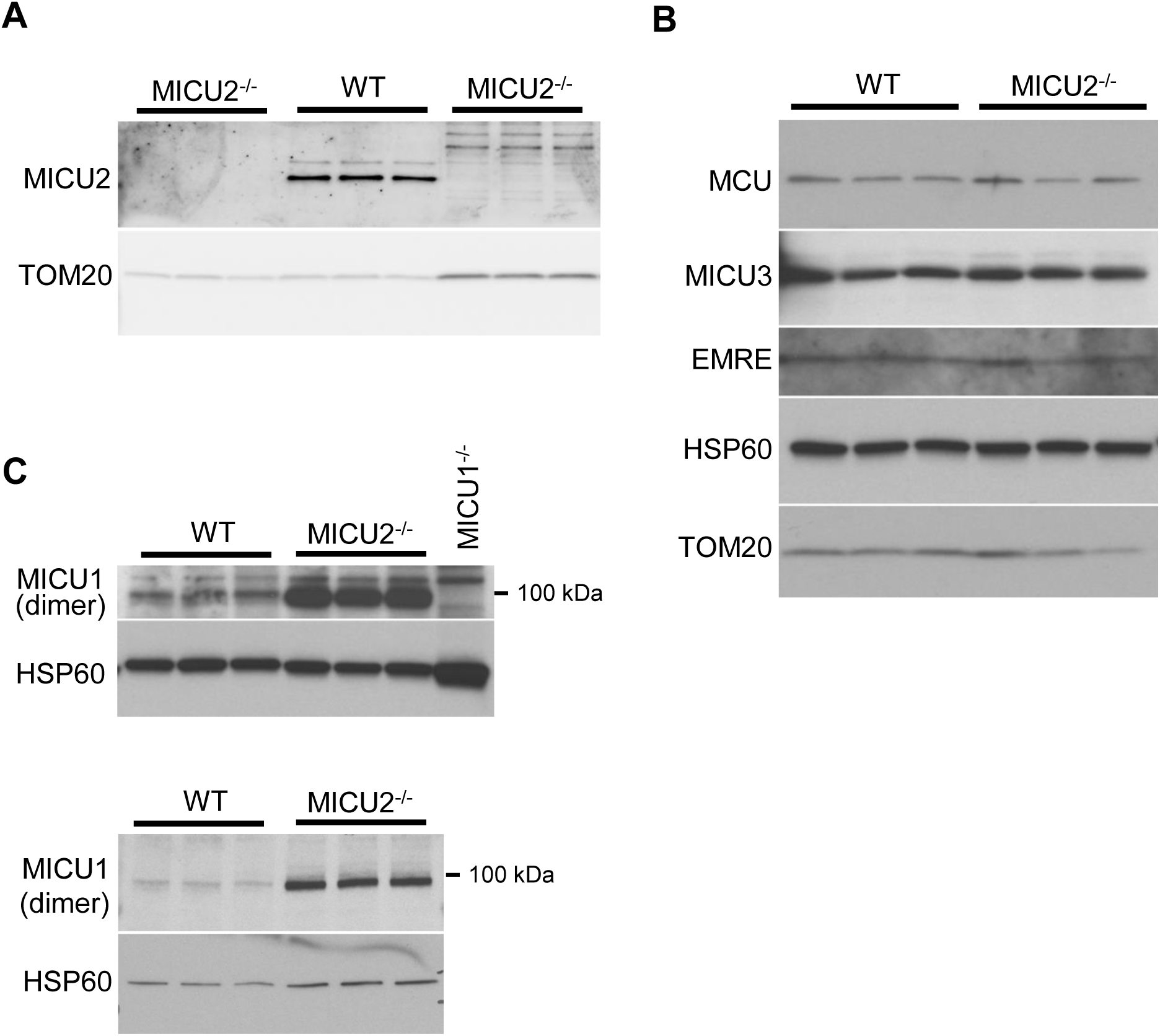
Expression levels of different MCU subunits in *MICU2*^*-/-*^ MEFs and the open probability of MCU at different potentials. (**A** to **C**) Western blots showing the expression levels of MICU2 (A), MCU, EMRE, MICU3 (B) and MICU1 dimers (C) in *WT* and *MICU2*^*-/-*^ cells (*n* = 3−6 independent samples each). For detection of MICU1 dimers, samples were prepared in Laemmli buffer without β-mercaptoethanol.

**Fig. S9.**
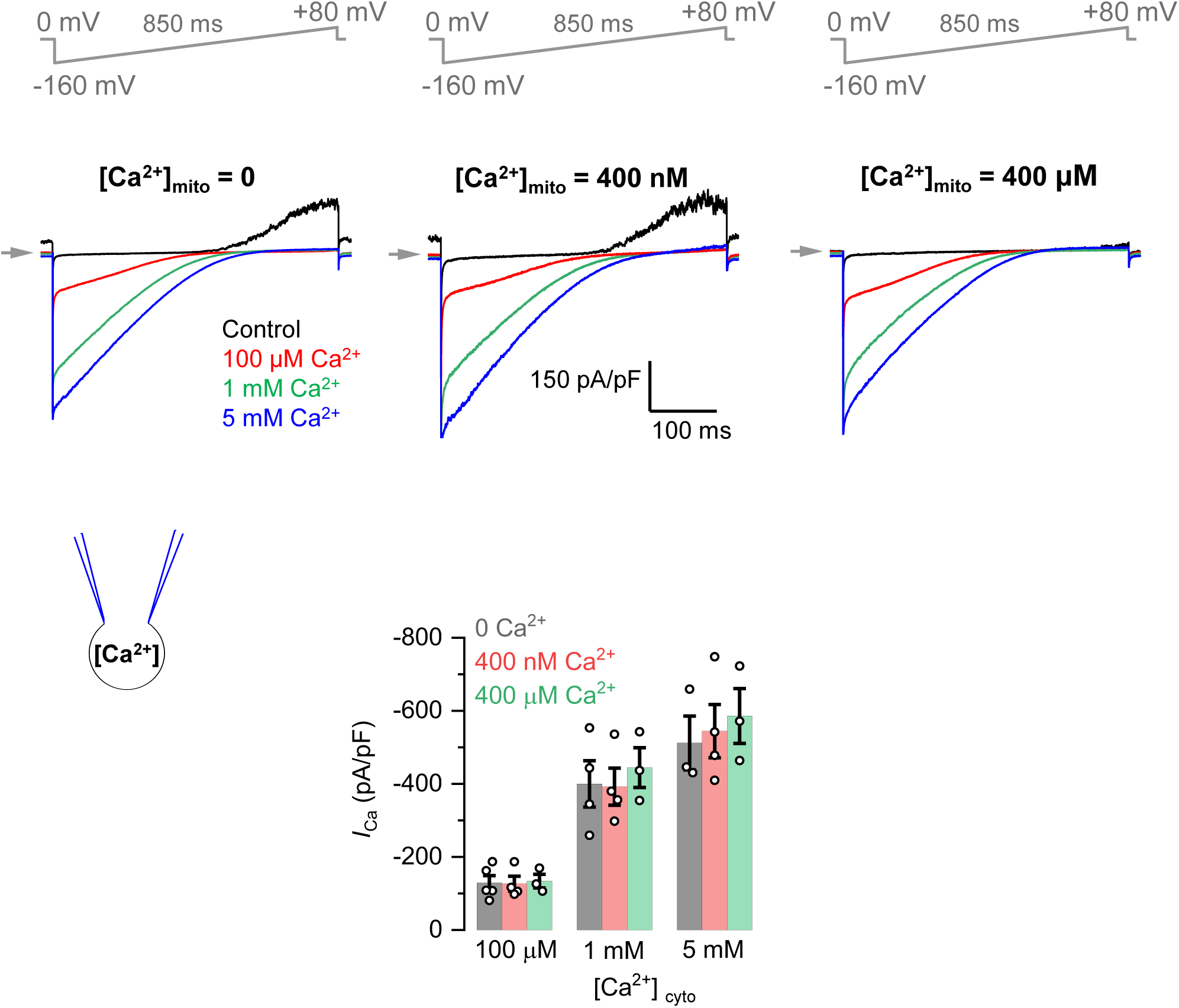
Matrix Ca^2+^ does not inhibit *I*_Ca._ (*Upper Panels*) Inward *I*_Ca_ in the presence of 0 (*left*), 400 nM (*middle*) and 400 μM (*right*) [Ca^2+^]_mito_ (pipette solution). [Ca^2+^]_cyto_ was 100 μM, 1 mM or 5 mM. (*Lower Panel*) *I*_Ca_ amplitudes at 0 (*n*=3-5), 400 nM (*n*=4) or 400 μM (*n*=3) [Ca^2+^]_mito_. *I*_Ca_ was measured at −160 mV and in different [Ca^2+^]_cyto_ as indicated. Mean ± SEM; one-way ANOVA with post-hoc Tuckey test.

**Fig. S10.**
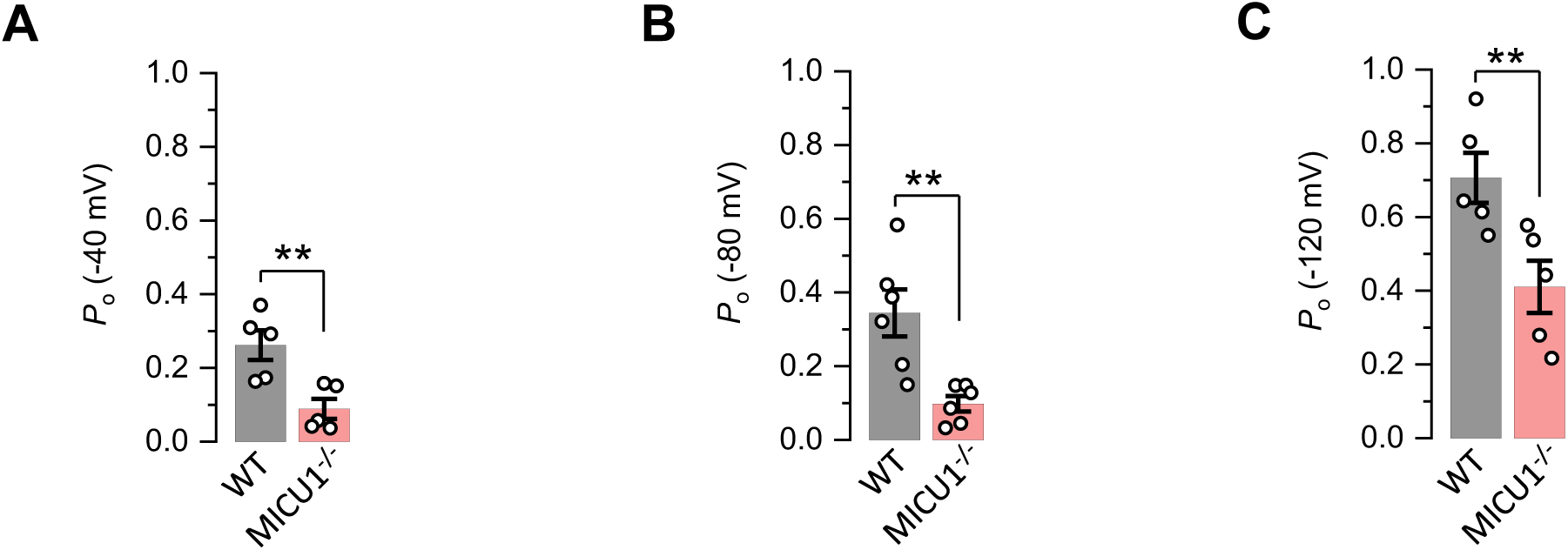
Open probability of the MCU channel in *WT* and *MICU1*^*-/-*^. (**A** to **C**) Open probability of the MCU channel in *WT* (*n*=5-6) and *MICU1*^*-/-*^ (*n*=5) at −40 mV (A) −80 mV (B) and at −120 mV (C). The same *WT* and knockout data were used as in Fig. 4D, but presented to show data distribution. Mean ± SEM; unpaired t-test, two-tailed; \*\**p*< 0.01.

**Table S1.**
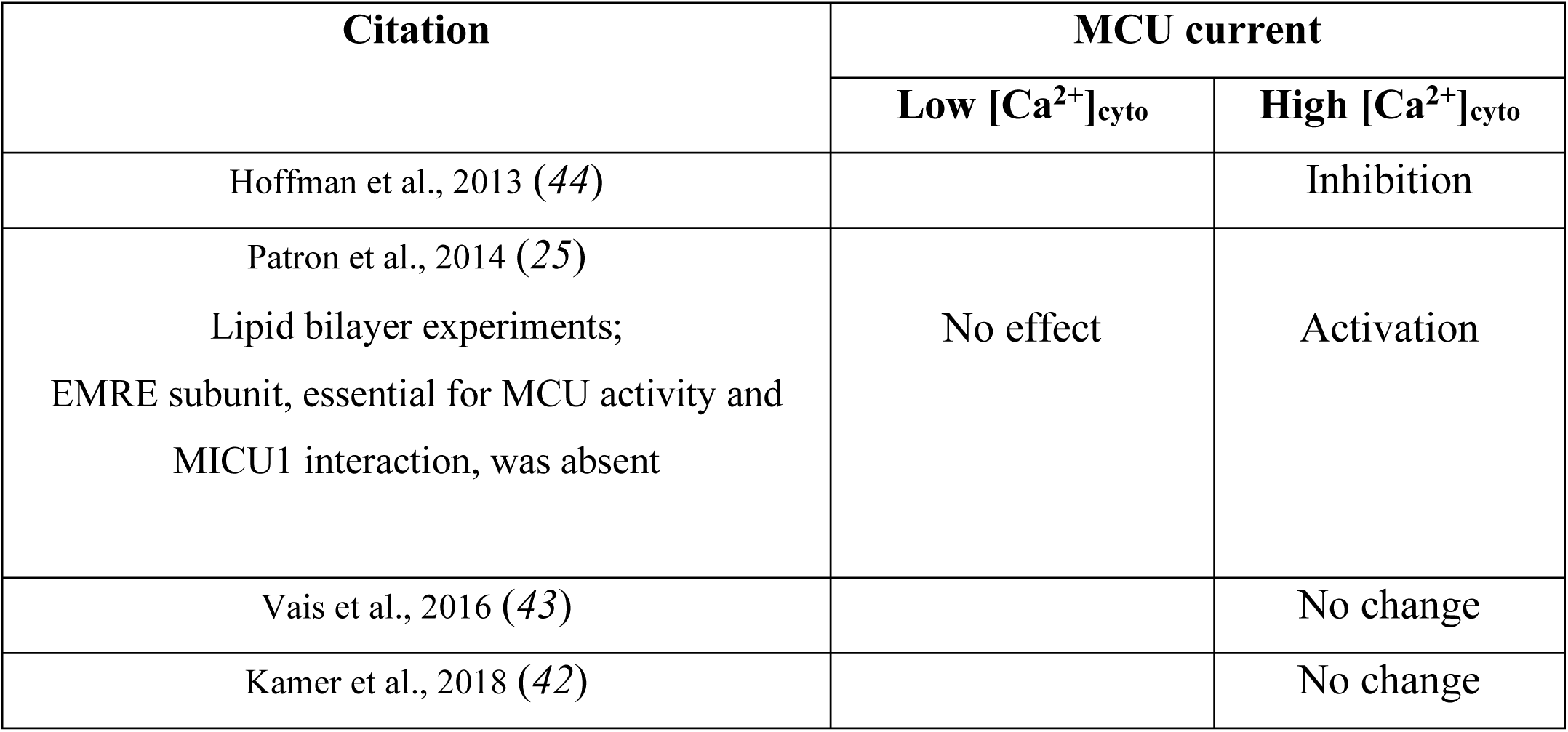
MICU1 effect on MCU as determined by previous electrophysiological experiments.

**Table S2.**
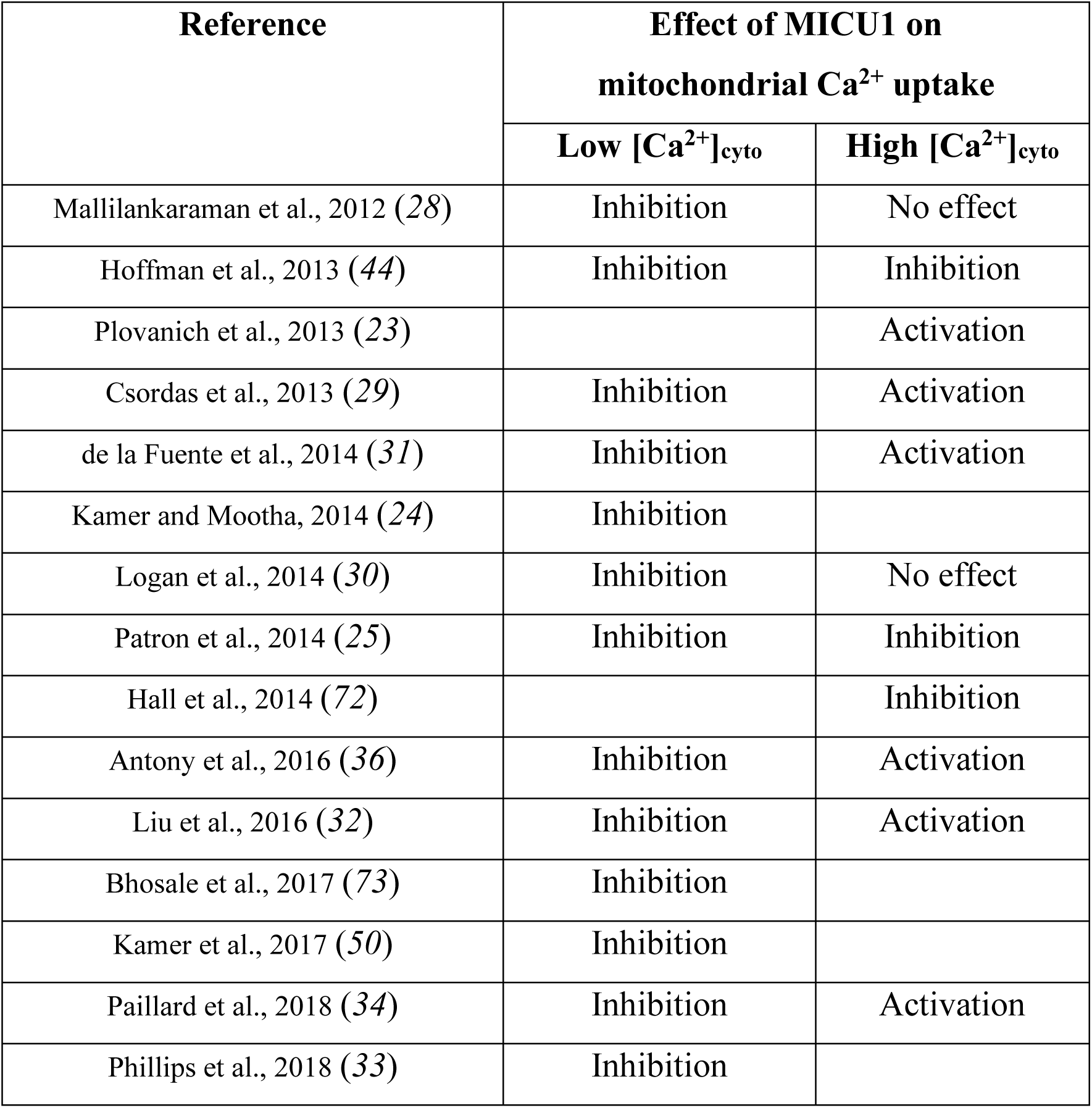
MICU1 effect on MCU as determined by previous Ca^2+^ imaging experiments.

## REFERENCES

1. B. Glancy, R. S. Balaban, Role of mitochondrial Ca2+ in the regulation of cellular energetics. Biochemistry 51, 2959–2973 (2012).

2. M. J. Berridge, M. D. Bootman, H. L. Roderick, Calcium signalling: dynamics, homeostasis and remodelling. Nature reviews 4, 517–529 (2003).

3. P. Bernardi, Mitochondrial transport of cations: channels, exchangers, and permeability transition. Physiol Rev 79, 1127–1155 (1999).

4. T. E. Gunter, D. R. Pfeiffer, Mechanisms by which mitochondria transport calcium. Am J Physiol 258, C755–786 (1990).

5. H. F. Deluca, G. W. Engstrom, Calcium uptake by rat kidney mitochondria. Proc Natl Acad Sci U S A 47, 1744–1750 (1961).

6. L. Tang et al., Structural basis for Ca2+ selectivity of a voltage-gated calcium channel. Nature 505, 56–61 (2014).

7. P. Hess, J. B. Lansman, R. W. Tsien, Calcium channel selectivity for divalent and monovalent cations. Voltage and concentration dependence of single channel current in ventricular heart cells. J Gen Physiol 88, 293–319 (1986).

8. P. Hess, R. W. Tsien, Mechanism of ion permeation through calcium channels. Nature 309, 453–456 (1984).

9. Y. Kirichok, G. Krapivinsky, D. E. Clapham, The mitochondrial calcium uniporter is a highly selective ion channel. Nature 427, 360–364 (2004).

10. F. Fieni, S. B. Lee, Y. N. Jan, Y. Kirichok, Activity of the mitochondrial calcium uniporter varies greatly between tissues. Nat Commun 3, 1317 (2012).

11. N. X. Nguyen et al., Cryo-EM structure of a fungal mitochondrial calcium uniporter. Nature 559, 570–574 (2018).

12. C. Fan et al., X-ray and cryo-EM structures of the mitochondrial calcium uniporter. Nature 559, 575–579 (2018).

13. R. Baradaran, C. Wang, A. F. Siliciano, S. B. Long, Cryo-EM structures of fungal and metazoan mitochondrial calcium uniporters. Nature 559, 580–584 (2018).

14. Y. Wang et al., Structural Mechanism of EMRE-Dependent Gating of the Human Mitochondrial Calcium Uniporter. Cell 177, 1252–1261 e1213 (2019).

15. J. Yoo et al., Cryo-EM structure of a mitochondrial calcium uniporter. Science 361, 506–511 (2018).

16. P. Bernardi, A. Rasola, Calcium and cell death: the mitochondrial connection. Subcell Biochem 45, 481–506 (2007).

17. H. Kroner, Ca2+ions, an allosteric activator of calcium uptake in rat liver mitochondria. Arch Biochem Biophys 251, 525–535 (1986).

18. A. Vinogradov, A. Scarpa, The initial velocities of calcium uptake by rat liver mitochondria. J Biol Chem 248, 5527–5531 (1973).

19. J. M. Baughman et al., Integrative genomics identifies MCU as an essential component of the mitochondrial calcium uniporter. Nature 476, 341–345 (2011).

20. D. De Stefani, A. Raffaello, E. Teardo, I. Szabo, R. Rizzuto, A forty-kilodalton protein of the inner membrane is the mitochondrial calcium uniporter. Nature 476, 336–340 (2011).

21. Y. Sancak et al., EMRE is an essential component of the mitochondrial calcium uniporter complex. Science 342, 1379–1382 (2013).

22. F. Perocchi et al., MICU1 encodes a mitochondrial EF hand protein required for Ca(2+) uptake. Nature 467, 291–296 (2010).

23. M. Plovanich et al., MICU2, a paralog of MICU1, resides within the mitochondrial uniporter complex to regulate calcium handling. PLoS One 8, e55785 (2013).

24. K. J. Kamer, V. K. Mootha, MICU1 and MICU2 play nonredundant roles in the regulation of the mitochondrial calcium uniporter. EMBO Rep 15, 299–307 (2014).

25. M. Patron et al., MICU1 and MICU2 finely tune the mitochondrial Ca2+ uniporter by exerting opposite effects on MCU activity. Mol Cell 53, 726–737 (2014).

26. M. F. Tsai et al., Dual functions of a small regulatory subunit in the mitochondrial calcium uniporter complex. eLife 5, (2016).

27. M. Patron, V. Granatiero, J. Espino, R. Rizzuto, D. De Stefani, MICU3 is a tissue-specific enhancer of mitochondrial calcium uptake. Cell Death Differ 26, 179–195 (2019).

28. K. Mallilankaraman et al., MICU1 is an essential gatekeeper for MCU-mediated mitochondrial Ca(2+) uptake that regulates cell survival. Cell 151, 630–644 (2012).

29. G. Csordas et al., MICU1 controls both the threshold and cooperative activation of the mitochondrial Ca(2)(+) uniporter. Cell Metab 17, 976–987 (2013).

30. C. V. Logan et al., Loss-of-function mutations in MICU1 cause a brain and muscle disorder linked to primary alterations in mitochondrial calcium signaling. Nat Genet 46, 188–193 (2014).

31. S. de la Fuente, J. Matesanz-Isabel, R. I. Fonteriz, M. Montero, J. Alvarez, Dynamics of mitochondrial Ca2+ uptake in MICU1-knockdown cells. Biochem J 458, 33–40 (2014).

32. J. C. Liu et al., MICU1 Serves as a Molecular Gatekeeper to Prevent In Vivo Mitochondrial Calcium Overload. Cell Rep 16, 1561–1573 (2016).

33. C. B. Phillips, C. W. Tsai, M. F. Tsai, The conserved aspartate ring of MCU mediates MICU1 binding and regulation in the mitochondrial calcium uniporter complex. eLife 8, (2019).

34. M. Paillard et al., MICU1 Interacts with the D-Ring of the MCU Pore to Control Its Ca(2+) Flux and Sensitivity to Ru360. Mol Cell 72, 778–785 e773 (2018).

35. J. Matesanz-Isabel et al., Functional roles of MICU1 and MICU2 in mitochondrial Ca(2+) uptake. Biochim Biophys Acta 1858, 1110–1117 (2016).

36. A. N. Antony et al., MICU1 regulation of mitochondrial Ca(2+) uptake dictates survival and tissue regeneration. Nat Commun 7, 10955 (2016).

37. D. Lewis-Smith et al., Homozygous deletion in MICU1 presenting with fatigue and lethargy in childhood. Neurol Genet 2, e59 (2016).

38. S. Musa et al., A Middle Eastern Founder Mutation Expands the Genotypic and Phenotypic Spectrum of Mitochondrial MICU1 Deficiency: A Report of 13 Patients. JIMD Rep 43, 79–83 (2019).

39. G. Bhosale et al., Pathological consequences of MICU1 mutations on mitochondrial calcium signalling and bioenergetics. Biochim Biophys Acta 1864, 1009–1017 (2017).

40. D. G. Nicholls, Mitochondria and calcium signaling. Cell Calcium 38, 311–317 (2005).

41. V. Garg, Y. Y. Kirichok, Patch-Clamp Analysis of the Mitochondrial Calcium Uniporter. Methods Mol Biol 1925, 75–86 (2019).

42. K. J. Kamer et al., MICU1 imparts the mitochondrial uniporter with the ability to discriminate between Ca(2+) and Mn(2+). Proc Natl Acad Sci U S A 115, E7960–E7969 (2018).

43. H. Vais et al., EMRE Is a Matrix Ca(2+) Sensor that Governs Gatekeeping of the Mitochondrial Ca(2+) Uniporter. Cell Rep 14, 403–410 (2016).

44. N. E. Hoffman et al., MICU1 motifs define mitochondrial calcium uniporter binding and activity. Cell Rep 5, 1576–1588 (2013).

45. N. Ishihara et al., Mitochondrial fission factor Drp1 is essential for embryonic development and synapse formation in mice. Nat Cell Biol 11, 958–966 (2009).

46. J. Suzuki et al., Imaging intraorganellar Ca2+ at subcellular resolution using CEPIA. Nat Commun 5, 4153 (2014).

47. X. Pan et al., The physiological role of mitochondrial calcium revealed by mice lacking the mitochondrial calcium uniporter. Nat Cell Biol 15, 1464–1472 (2013).

48. E. Kovacs-Bogdan et al., Reconstitution of the mitochondrial calcium uniporter in yeast. Proc Natl Acad Sci U S A 111, 8985–8990 (2014).

49. A. Raffaello et al., The mitochondrial calcium uniporter is a multimer that can include a dominant-negative pore-forming subunit. Embo J 32, 2362–2376 (2013).

50. K. J. Kamer, Z. Grabarek, V. K. Mootha, High-affinity cooperative Ca(2+) binding by MICU1-MICU2 serves as an on-off switch for the uniporter. EMBO Rep 18, 1397–1411 (2017).

51. P. Bernardi, A. Angrilli, G. F. Azzone, A gated pathway for electrophoretic Na+ fluxes in rat liver mitochondria. Regulation by surface Mg2+. Eur J Biochem 188, 91–97 (1990).

52. J. P. Wehrle, M. Jurkowitz, K. M. Scott, G. P. Brierley, Mg2+ and the permeability of heart mitochondria to monovalent cations. Arch Biochem Biophys 174, 313–323 (1976).

53. C. Petrungaro et al., The Ca(2+)-Dependent Release of the Mia40-Induced MICU1-MICU2 Dimer from MCU Regulates Mitochondrial Ca(2+) Uptake. Cell Metab 22, 721–733 (2015).

54. Y. Xing et al., Dimerization of MICU Proteins Controls Ca(2+) Influx through the Mitochondrial Ca(2+) Uniporter. Cell Rep 26, 1203–1212 e1204 (2019).

55. R. Rizzuto et al., Close contacts with the endoplasmic reticulum as determinants of mitochondrial Ca2+ responses. Science 280, 1763–1766 (1998).

56. E. Neher, Vesicle pools and Ca2+ microdomains: new tools for understanding their roles in neurotransmitter release. Neuron 20, 389–399 (1998).

57. G. C. Faas, S. Raghavachari, J. E. Lisman, I. Mody, Calmodulin as a direct detector of Ca2+ signals. Nat Neurosci 14, 301–304 (2011).

58. P. Bernardi, V. Petronilli, The permeability transition pore as a mitochondrial calcium release channel: a critical appraisal. J Bioenerg Biomembr 28, 131–138 (1996).

59. F. Ichas, L. S. Jouaville, J. P. Mazat, Mitochondria are excitable organelles capable of generating and conveying electrical and calcium signals. Cell 89, 1145–1153 (1997).

60. M. Montero, M. T. Alonso, A. Albillos, J. Garcia-Sancho, J. Alvarez, Mitochondrial Ca(2+)-induced Ca(2+) release mediated by the Ca(2+) uniporter. Mol Biol Cell 12, 63–71 (2001).

61. U. Igbavboa, D. R. Pfeiffer, Regulation of reverse uniport activity in mitochondria by extramitochondrial divalent cations. Dependence on a soluble intermembrane space component. J Biol Chem 266, 4283–4287 (1991).

62. H. Vais et al., EMRE Is a Matrix Ca(2+) Sensor that Governs Gatekeeping of the Mitochondrial Ca(2+) Uniporter. Cell reports 14, 403–410 (2016).

63. T. E. Gunter et al., An analysis of the effects of Mn2+ on oxidative phosphorylation in liver, brain, and heart mitochondria using state 3 oxidation rate assays. Toxicol Appl Pharmacol 249, 65–75 (2010).

64. J. Wettmarshausen et al., MICU1 Confers Protection from MCU-Dependent Manganese Toxicity. Cell Rep 25, 1425–1435 e1427 (2018).

65. B. Hille, Ionic channels of excitable membranes. (Sinauer Associates, Sunderland, Mass., ed. 2nd, 1992), pp. xiii, 607.

66. A. P. Wescott, J. P. Y. Kao, W. J. Lederer, L. Boyman, Voltage-energized calcium-sensitive ATP production by mitochondria. Nature Metabolism 1, 975–984 (2019).

67. M. S. Bertolini, M. A. Chiurillo, N. Lander, A. E. Vercesi, R. Docampo, MICU1 and MICU2 Play an Essential Role in Mitochondrial Ca(2+) Uptake, Growth, and Infectivity of the Human Pathogen Trypanosoma cruzi. MBio 10, (2019).

68. R. Tufi et al., Comprehensive Genetic Characterization of Mitochondrial Ca(2+) Uniporter Components Reveals Their Different Physiological Requirements In Vivo. Cell Rep 27, 1541–1550 e1545 (2019).

69. F. A. Ran et al., Genome engineering using the CRISPR-Cas9 system. Nat Protoc 8, 2281–2308 (2013).

70. G. Grynkiewicz, M. Poenie, R. Y. Tsien, A new generation of Ca2+ indicators with greatly improved fluorescence properties. J Biol Chem 260, 3440–3450 (1985).

71. D. A. Winter, Biomechanics and motor control of human movement. (Wiley, Hoboken, N.J., ed. 4th, 2009), pp. xiv, 370 pages.

72. D. D. Hall, Y. Wu, F. E. Domann, D. R. Spitz, M. E. Anderson, Mitochondrial calcium uniporter activity is dispensable for MDA-MB-231 breast carcinoma cell survival. PLoS One 9, e96866 (2014).

73. G. Bhosale et al., Pathological consequences of MICU1 mutations on mitochondrial calcium signalling and bioenergetics. Biochim Biophys Acta Mol Cell Res 1864, 1009–1017 (2017).

