## Supplementary Materials for "The Mechanism of MICU-Dependent Gating of the Mitochondrial Ca^2+^ Uniporter"

**This PDF file includes:**

Materials and Methods
Figs. S1 to S10
Tables S1 to S2
References

#### MATERIALS AND METHODS

##### *Cell culture and recombinant gene expression*

All mouse embryonic fibroblast (MEF) cells with (*32*) or without Drp1 (*46*), and all knockout clones were grown in low glucose (5.6 mM) Dulbecco's modified Eagle's medium (DMEM) supplemented with 10% FBS, 100 U/ml penicillin, and 100 U/ml streptomycin at 37$^o$C, 5% $CO_2$. Cells were maintained by splitting every 48-72 hours at a ratio of 1:5 to 1:10.

We used third generation lentiviral (bi-cistronic) vectors containing the ORF for gene of interest with or without a selection marker (EGFP, mCherry or puromycin, Supplementary Fig. 1 and 2). The vectors were generated by VectorBuilder, Inc. (Chicago, IL, USA), and their sequences were confirmed independently by the company and by us. Recombinant cDNA expressing cells were enriched using multiple rounds of FACS or antibiotic selection. In some cases, EGFP was targeted to mitochondria (using a mitochondrial targeting sequence from COX8) to identify mitoplasts expressing the recombinant protein of interest during patch clamp experiments.

##### *Quantitative Real-Time PCR Analysis*

qPCR was performed by Syd Labs (Natick, MA, USA). Total RNA was isolated from cells using the RNAeasy Minikit (QIAGEN), and reverse transcribed using the First Strand cDNA Synthesis Kit (Syd Labs). qPCR reactions were performed with the following gene-specific primers (generated by Integrated DNA Technologies):

| | | |
|---|---|---|
| Hprt, | Forward Primer | 5'-GTCCCAGCGTCGTGATTAGC-3' |
| | Reverse Primer | 5'-GTGATGGCCTCCCATCTCCT-3' |
| MCU, | Forward Primer | 5'-AAGGGCTTAGCGAGTCTTGTC-3' |
| | Reverse Primer | 5'- GGGTGCTGGTGTGTTAGTGT -3' |
| MCUb, | Forward Primer | 5'-CCACACCCCAGGTTTTATGTATG-3' |
| | Reverse Primer | 5'-ATGGCAGAGTGAGGGTTACCA-3' |
| EMRE, | Forward Primer | 5'-ATTTTGCCCAAGCCGGTGAA-3' |
| | Reverse Primer | 5'-CCTCAAGCAGAGCAGCGAAG-3' |
| MICU1, | Forward Primer | 5'-CTTAACACCCTTTCTGCGTTGG-3' |
| | Reverse Primer | 5'-AGCATCAATCTTCGTTTGGTCT-3' |
| MICU2, | Forward Primer | 5'-CTCCGCAAACAGCGGTTCAT-3' |
| | Reverse Primer | 5'-TGCCAGCTTCTTGACCAGTG-3' |
| MICU3, | Forward Primer | 5'-GTAAGGTCAGAGCACGCAGAA-3' |
| | Reverse Primer | 5'-TTTCCTGTTGGACGCTGACAA -3' |

cDNA (100 ng, calculated from initial RNA) samples were pre-amplified for 12 cycles using ABsolute qPCR SYBR Green Low ROX Mix (ThermoFisher). qPCR reactions were performed using an Agilent MX3000 (Fluidigm) with 40 cycles of amplification (15 s at 95$^o$C, 5 s at 70$^o$C, and 60 s at 60$^o$C). Ct values were calculated by the Real-Time PCR Analysis Software (Fluidigm). Relative gene expression was determined by the $\Delta$Ct method. Hprt was selected as the reference gene.

##### *Generation of knockout cell lines by the CRISPR/Cas9 method*

Knockout MEF cell lines were generated using the CRISPR/Cas9 method (*70*). All knockouts (except the *MCU*$^{-/-}$ line) were generated by Alstem LLC (Richmond, CA, USA). Either one sgRNA or a pair of two adjacent sgRNAs were used to create a point indel or a truncate indel, respectively (Fig. S1).

| | |
|---|---|
| MCU, | TGGCAGCGCTCGCGTCGAGA GGG |
| EMRE, | GAGTGTCCCGACATAGAGAA AGG |
| | CTTACACTCCCACTAGGTTA AGG |
| MICU1, | TCACTTTTAGATGCTGCCGG TGG |
| | CTGCAAGTACCGGTCTCCTG TGG |
| MICU2, | CGTTCGGGAGCCCTCGCGCG CGG |
| | GGGCGCTTCCGCAAAGATGG CGG |
| MICU3, | GGGCGAGCTGAGCATCGCGG CGG |
| | CCGGGGCCGCTAGCTCCGAG GGG |

MEFs were transfected with the Cas9 gRNA vector (Addgene: PX459) via electroporation (Invitrogen Neon transfection system) using the following parameters: $1\times10^6$ cells and 1 µg of two different gRNA-Cas9 plasmids. Puromycin was used for enrichment of transfected cells, and serial dilution was performed to select single-cell clones. A stable homozygous knockout cell line was confirmed by PCR amplification of the targeted region, cloning into a pUC19 vector, and sequencing showing that either a frameshift or large deletion had occurred in the targeted region of the gene (Fig. S1). All knockout clones were further validated by Western blotting (Fig. S2). The primers used for amplification of genomic sites and cloning into pUC19 sequencing vector were as follows:

| | | |
|---|---|---|
| MCU, | Forward Primer | TAGAAGCTTTCCACTGCTCTGATTGATCTTG |
| | Reverse Primer | ATGTGAATTCGAGCTGCTTTGGAATGAGAC |
| EMRE, | Forward Primer | GTGAAGCTTGGGATCAGTAGTCCATTGGAGG |
| | Reverse Primer | AGGAGAATTCAGTGAGAGTTCCTGTGGTATG |
| MICU1, | Forward Primer | TTTAAGCTTGATTCCTTTGAGTTATAAGTAG |
| | Reverse Primer | CAAAGAATTCAGCAAAGAAATTCTGATGTA |
| MICU2, | Forward Primer | ACCAAGCTTGAACGTCGAGGAAGCAGCCAC |
| | Reverse Primer | AGGAGAATTCTCCATCCACCAGGTGGGCAG |
| MICU3, | Forward Primer | CGCAAGCTTCTCGCGAGATTTCGGCCCGCC |
| | Reverse Primer | AGGAGAATTCTCCATCCACCAGGTGGGCAG |

$$[\mathrm{Ca}^{2+}]\mathrm{free} = \mathrm{K_d} * \left(\frac{[\mathrm{R} - \mathrm{Rmin}]}{[\mathrm{Rmax} - \mathrm{R}]}\right) * (\mathrm{F}_{380max}/\mathrm{F}_{380min})$$

All image analyses were done with ImageJ (NIH). Briefly, mitochondrial and cytosolic regions were manually determined for each cell. The average fluorescence intensity in the regions was measured and the background intensity was subtracted. For analysis of the *cepia2* signal, we normalized the fluorescence intensity by the baseline fluorescence. For analysis of the fura-2 signal, we calculated the fluorescence ratio ($F_{340}/F_{380}$ for fura-2).

#### SUPPLEMENTARY FIGURES

**Fig. S1. Generation of knockouts for various MCU complex subunits.** (**A**) A schematic arrangement of various subunits in the MCU complex. Four MCU and four EMRE subunits form the pore of the MCU complex (only two MCU and two EMRE subunits are shown for simplicity). EMRE also tethers MICU1 subunit to the pore on the cytosolic side of the IMM (i.e., in the mitochondrial intermembrane space, IMS). MICU1 forms homodimers or hetero-dimerizes with MICU2 or MICU3 (not shown). Each MICU subunit has two EF hands that bind cytosolic $Ca^{2+}$. (**B** to **F**) CRISPR-mediated indels in various MCU subunit genes and the resulting mutant alleles. The CRISPR binding sites (for sgRNA) are highlighted in *yellow*, and their PAM sequences are highlighted in *green*. The translational initiation codon (ATG) is shown in *bold* where applicable. (B) Overview of the *MCU* gene and indels in the knockout. A sgRNA was used to target exon 3. The sequence of targeted region in MCU gene is shown; exon 3 is underlined. Targeted sequencing indicates frame-shift indels (*red*) in both alleles (*Al*-1 and *Al*-2). (C) Overview of the *EMRE* gene and truncated region in the knockout. Two sgRNAs were used for CRISPR-Cas9–mediated deletion in the exon-2 (*underlined*) and the flanking region. Targeted sequencing indicates same 259-bp deletion (*red*) in both alleles. (D) Overview of the *MICU1* gene and truncated region in the knockout. Two sgRNAs were used for CRISPR-Cas9–mediated deletion in the exon-3 (*underlined*) and the flanking region. Targeted sequencing indicates that almost all of exon-3 is deleted along with a portion of the flanking region (*red*) in both alleles (*Al*-1 and A*l*-2). (E) Overview of the *MICU2* gene and truncated region in the knockout. Two sgRNAs were used for CRISPR-Cas9–mediated deletion in the exon-1 (*underlined*) and the flanking region. Targeted sequencing indicates that almost all of exon-1 is deleted (*red*) in both alleles. (F) Overview of the *MICU3* gene and truncated region in the knockout. Two sgRNAs were used for CRISPR-Cas9–mediated deletion in the exon-1 (*underlined*). Targeted sequencing indicates a 73-bp deletion in the expected cut area (*red*) in both alleles.

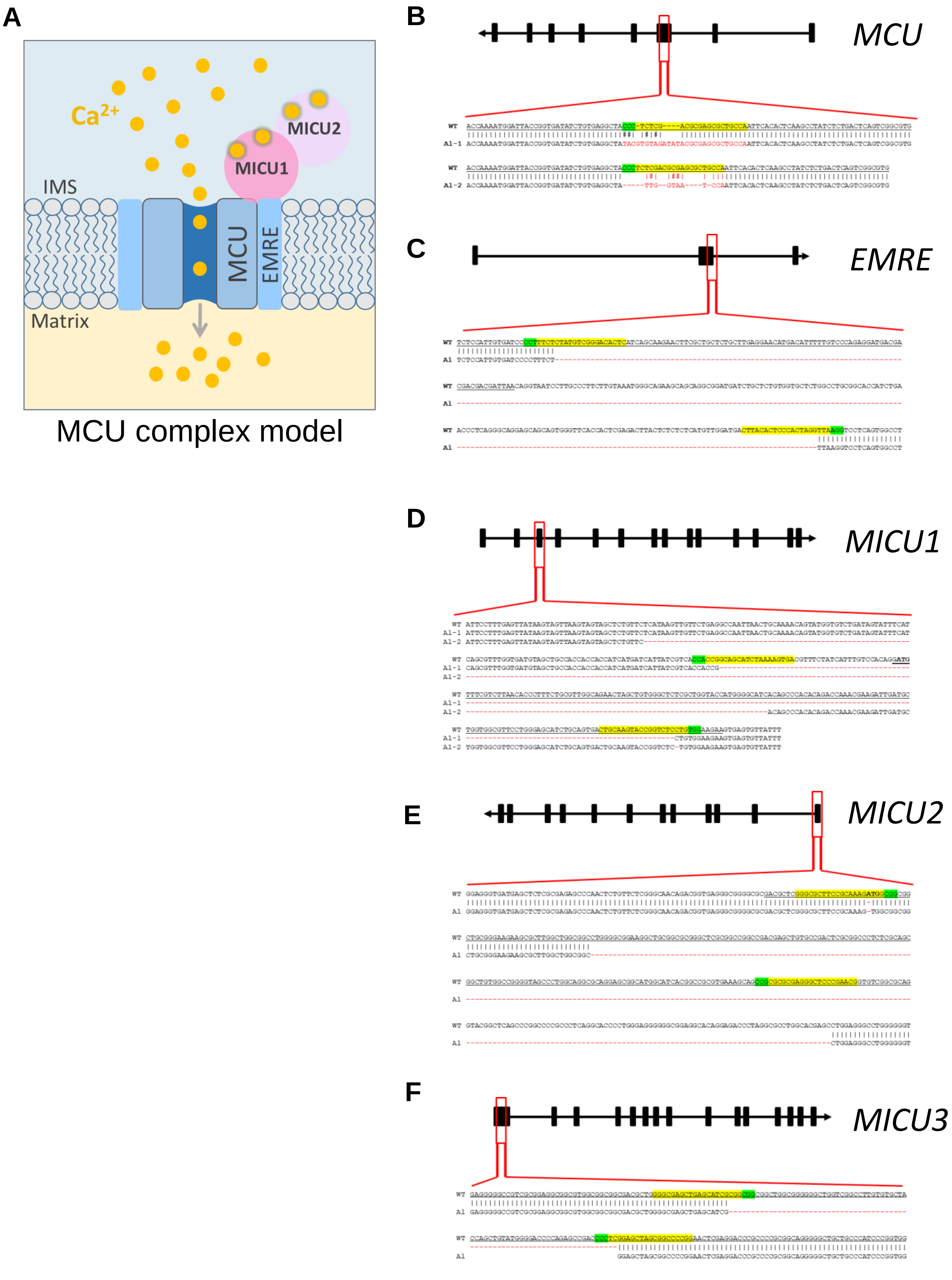
A
Ca2+
MICU2
MICU1
IMS
MCU
EMRE
Matrix
MCU complex model
B
MCU
C
EMRE
D
MICU1
E
MICU2
F
MICU3



**Fig. S1**

**Fig. S2. Expression of various MCU complex subunits and $[Ca^{2+}]_{mito}$ phenotype in cells deficient for various MCU complex subunits.** (**A** to **C**) Western blots show expression of various MCU complex subunits in the respective knockout cells. For MICU1 (**A**), samples were prepared without reducing agent, β-mercaptoethanol. The MICU1 band is near the expected molecular weight (~100 kDa) for the homo- or hetero-dimer (with MICU2 or 3). Multiple bands were observed with anti-MICU2 (**B**) and anti-MICU3 (**C**) antibodies, which were absent in knockout cell lines. This is likely due to the presence of different oligomeric states of the protein, as well as the mature and nascent (before truncation of the mitochondrial targeting signal) forms of the protein. Arrows mark the mature (m) and nascent (n) proteins near the expected molecular weight. (**D**) PCR showing the mRNA expression of various MCU subunits in *Drp1*$^{-/-}$ MEFs. Hprt was used as the reference. (**E** to **J**) Representative $[Ca^{2+}]_{mito}$ (*black*, left ordinate) and $[Ca^{2+}]_{cyto}$ (*blue*, right ordinate) in an individual cell with *WT* MCU complex, and individual cells with MCU, EMRE, and MICU1–3 knockouts before and after application of 300 nM Tg (arrow). Dashed red lines indicate the $[Ca^{2+}]_{cyto}$ at which the $[Ca^{2+}]_{mito}$ starts to increase ("$[Ca^{2+}]_{cyto}$ threshold"). (**K** to **M**) Resting $[Ca^{2+}]_{cyto}$ (K), peak $[Ca^{2+}]_{cyto}$ after addition of Tg (L), and $[Ca^{2+}]_{cyto}$ threshold for $[Ca^{2+}]_{mito}$ elevation (M) in *WT* and indicated knockout cell lines. *WT* (*n* = 5 dishes, total cells = 150); *MCU*$^{-/-}$ (*n* = 3 dishes, total cells = 183); *EMRE*$^{-/-}$ (*n* = 4 dishes, total cells = 187); *MICU1*$^{-/-}$ (*n* = 4 dishes, total cells = 196); *MICU2*$^{-/-}$ (*n* = 4 dishes, total cells = 192); and *MICU3*$^{-/-}$ (*n* = 3 dishes, total cells = 115). Mean ± SEM; one-way ANOVA with post-hoc Tukey test. *$p$< 0.05; ***$p$< 0.001. Statistics was run on number of dishes.

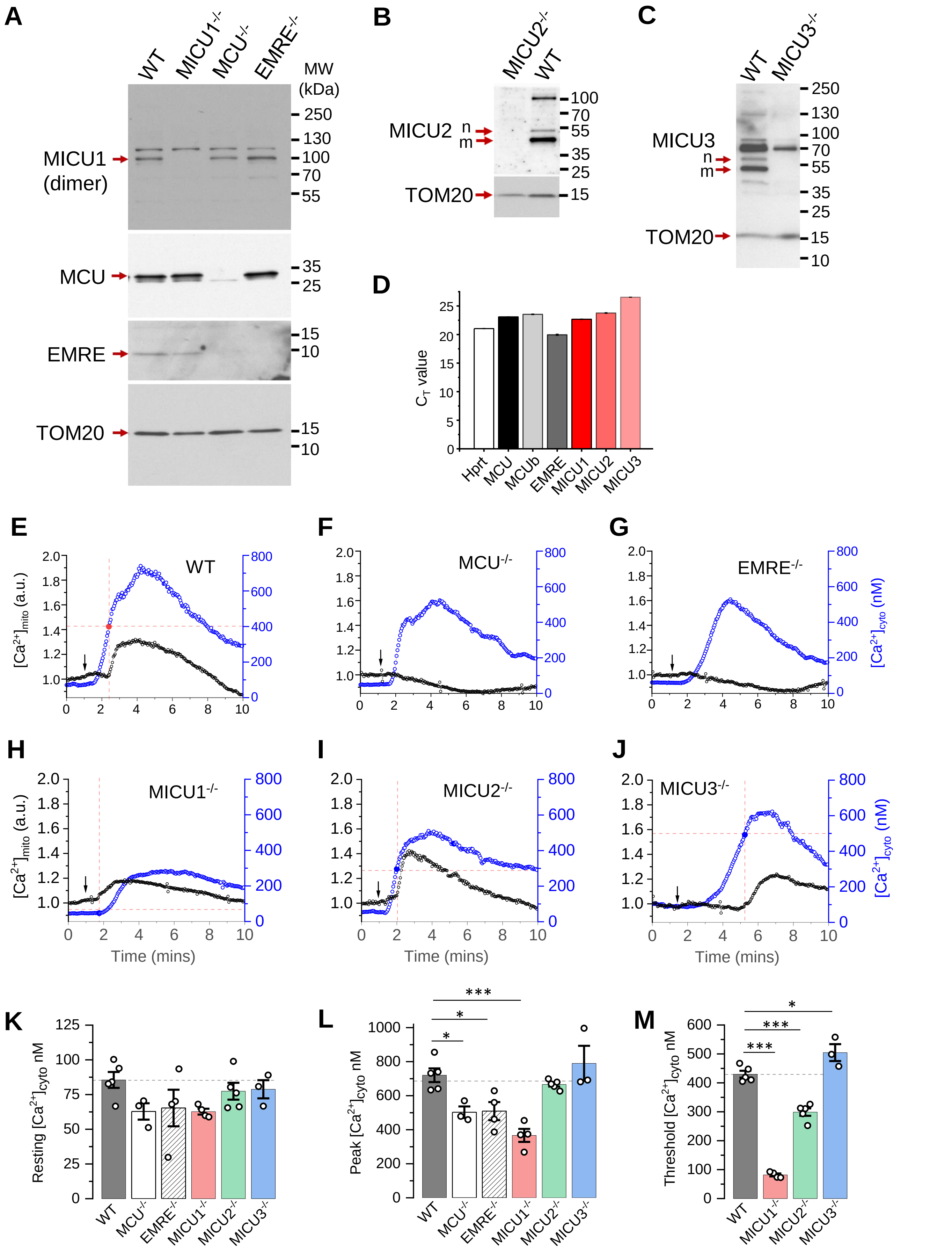

Fig. S2

**Fig. S3. Recording MCU currents across the whole IMM.** (**A**) Diagram of patch-clamp recording from a vesicle of the whole IMM (mitoplast). After formation of a gigaohm seal between the patch pipette and the mitoplast, the IMM patch under the pipette is broken by applying short pulses of high voltage (200–500 mV, 2–8 ms), sometimes combined with light suction, to gain access into the mitoplast through the pipette. In this configuration, called the "whole-IMM" configuration, the interior of the mitoplast (mitochondrial matrix) is perfused with the pipette solution. The bath is also perfused to control the experimental solution on the cytosolic side of the IMM. The voltage across the IMM is set to the desired value (V), and the currents (I) are measured using the patch-clamp amplifier. Directions of currents flowing across the IMM: inward currents (flowing into the mitoplast) are negative, while outward currents are positive. (**B**) *Left panel:* Example MCU current traces recorded in the whole-IMM configuration. The voltage protocol used to elicit the currents is shown above. All indicated voltages are within the mitochondrial matrix relative to the bath (cytosol). The voltage of the bath solution is defined to be zero. The zero current level is shown by the dashed line and an arrow. The directions of the currents are indicated as negative (inward) and positive (outward). The MCU current in $Ca^{2+}$-free bath solution (control) is shown in *grey*. The outward current in control is mediated by $Na^+$ ions permeating through the MCU channel in the $Ca^{2+}$-free conditions ($I_{Na}$, pipette solution contains $Na^+$). After application of 1 mM $Ca^{2+}$ on the cytosolic face of the IMM (bath), we observe an inward $Ca^{2+}$ current ($I_{Ca}$, *blue*) via MCU, while the outward $I_{Na}$ is simultaneously inhibited. *Right panel*, When $[Ca^{2+}]_{cyto}$ is brought to virtual zero (1 mM EGTA and 5 mM EDTA) under conditions when both bath and pipette solution contain $Na^+$, we observe $I_{Na}$ via MCU (*red*) in both inward and outward directions. The current amplitude and time calibration bars are indicated. The current amplitude is normalized per membrane capacitance to facilitate comparison of current amplitudes between mitoplasts of different sizes.

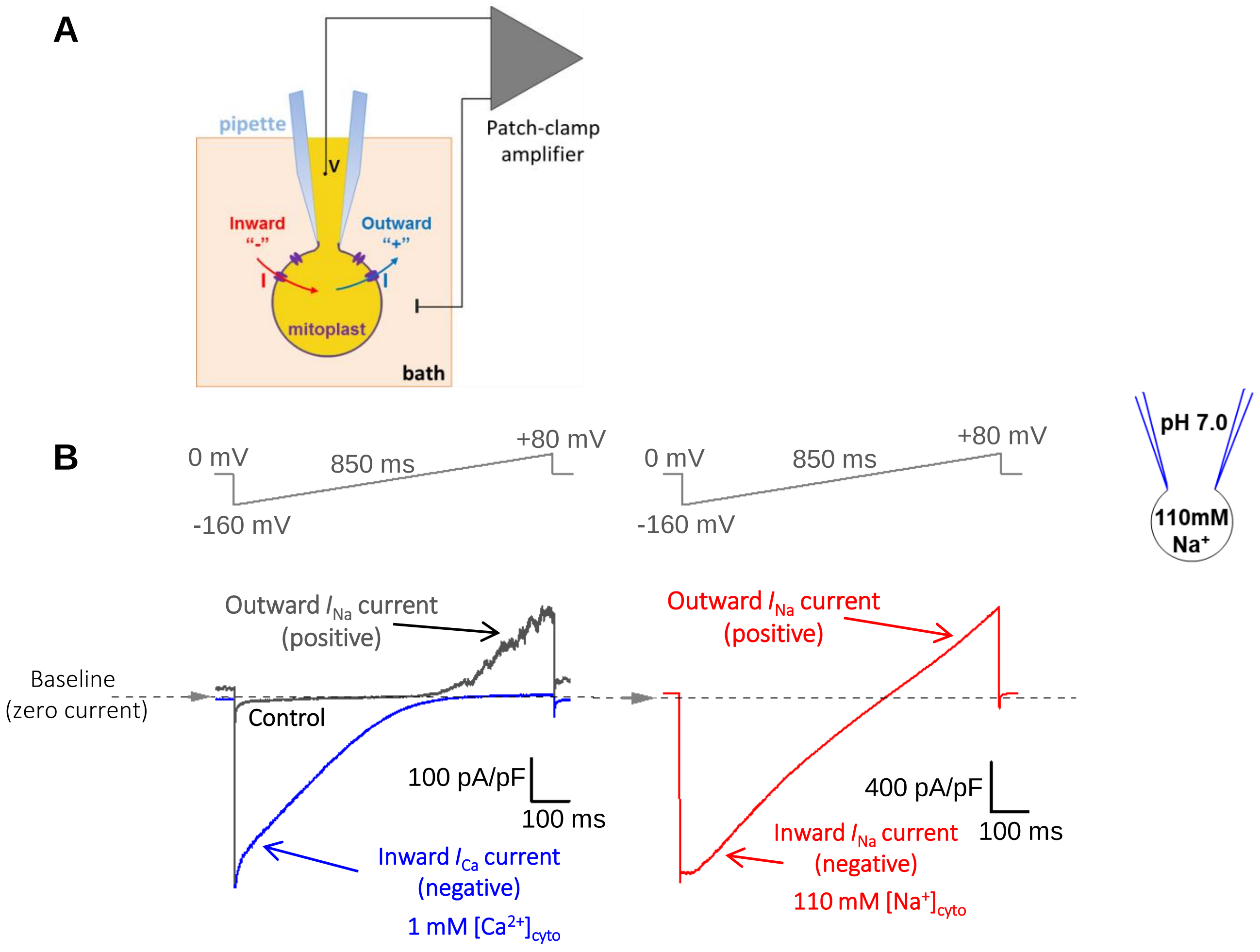

A
pipette
V
Patch-clamp
amplifier
Inward
"-"
Outward
"+"
mitoplast
bath
B
0 mV
850 ms
+80 mV
-160 mV
0 mV
850 ms
+80 mV
-160 mV
pH 7.0
110mM
Na+
Outward $I_{Na}$ current
(positive)
Baseline
(zero current)
Control
100 pA/pF
100 ms
Inward $I_{Ca}$ current
(negative)
1 mM $[Ca^{2+}]_{cyto}$
Outward $I_{Na}$ current
(positive)
400 pA/pF
100 ms
Inward $I_{Na}$ current
(negative)
110 mM $[Na^{+}]_{cyto}$




**Fig. S3**

**Fig. S4. Protein expression of MCU subunits in MEFs and isolated mitoplasts.** (**A**) Co-immunoprecipitation of the MCU complex proteins from mitochondrial and mitoplast fractions. Anti-FLAG beads were used to immunoprecipitate MCU-FLAG (expressed in *MCU*$^{-/-}$ cells) from mitochondrial and mitoplast fractions. Mitochondria isolated from *WT* cells (No FLAG tag) were used as negative control. Left three lanes are protein-complexes immunoprecipitated with anti-FLAG beads. Lane-1: immunoprecipitate (IP) from MCU-FLAG mitochondrial (Mt) lysate, lane-2: IP from MCU-FLAG mitoplast (Mp) lysate, lane-3: IP from *WT* mitochondrial lysate. Right three lanes correspond to samples from the flow-through fraction after immunoprecipitation. Lane-1*: mitochondrial lysate from MCU-FLAG, lane-2*: mitoplast lysate from MCU-FLAG, lane-3*: mitochondrial lysate from *WT*. Upper (MICU1, MCU and EMRE) and lower (MICU2 and TOM20) boxes are from the same samples run on different gels. (**B**) Western blots of protein lysates from cells with *WT* MCU complex (*WT*), *MCU*$^{-/-}$ cells, and *MCU*$^{-/-}$ cells overexpressing MCU (MCU-OE) using anti-MCU and anti-TOM20 (the mitochondrial loading control). (**C**) Western blots of protein lysates from *WT* cells, *EMRE*$^{-/-}$ cells, and *EMRE*$^{-/-}$ cells overexpressing EMRE (EMRE-OE) using anti-EMRE, anti-TOM20 and anti-HSP60 (the mitochondrial loading controls).

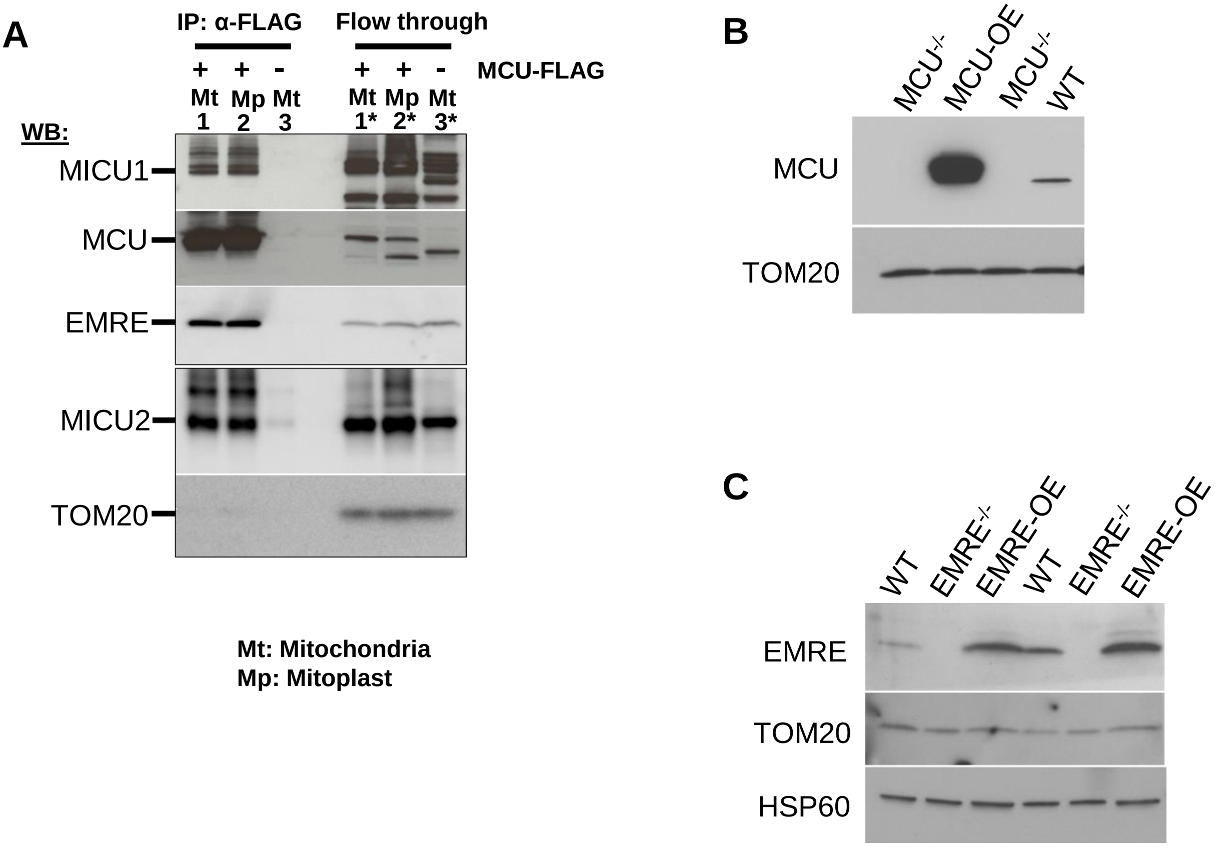
A
IP: α-FLAG
Flow through
+ + - + + -
MCU-FLAG
Mt 1
Mp 2
Mt 3
Mt 1*
Mp 2*
Mt 3*
WB:
MICU1
MCU
EMRE
MICU2
TOM20
Mt: Mitochondria
Mp: Mitoplast
B
MCU-/-
MCU-OE
MCU-/-
WT
MCU
TOM20
C
WT
EMRE-/-
EMRE-OE
WT
EMRE-/-
EMRE-OE
EMRE
TOM20
HSP60



### Fig. S4

**Fig. S5. $I_{Ca}$ in MICU1–3 knockouts.** (**A**) $I_{Ca}$ amplitude in *WT* and MICU1-3 knockouts measured at -160 mV using 10 μM $[Ca^{2+}]_{cyto}$ with an enlarged Y-axis. Data is same as used in Fig. 1E. Mean ± SEM; one-way ANOVA with post-hoc Tuckey test; $^{**}p < 0.01$. (**B**) Representative inward $I_{Ca}$ in *WT*, *MICU1*$^{-/-}$, *MICU2*$^{-/-}$ and *MICU3*$^{-/-}$ mitoplasts exposed to 5 mM, and 25 mM $[Ca^{2+}]_{cyto}$. (**C**) $I_{Ca}$ amplitude measured at -80 mV in *WT* ($n$ = 13), *MICU1*$^{-/-}$ ($n$ = 14), *MICU2*$^{-/-}$ ($n$ = 8) and *MICU3*$^{-/-}$ ($n$ = 9) mitoplasts at 10 μM, 100 μM and 1000 μM $[Ca^{2+}]_{cyto}$ (*left*, for $I_{Ca}$ traces see Fig. 2A) and at 5 mM and 25 mM $[Ca^{2+}]_{cyto}$ (*right*, for $I_{Ca}$ traces see Figure 1D). Mean ± SEM; one-way ANOVA with post-hoc Tuckey test; $^{***}p < 0.001$.

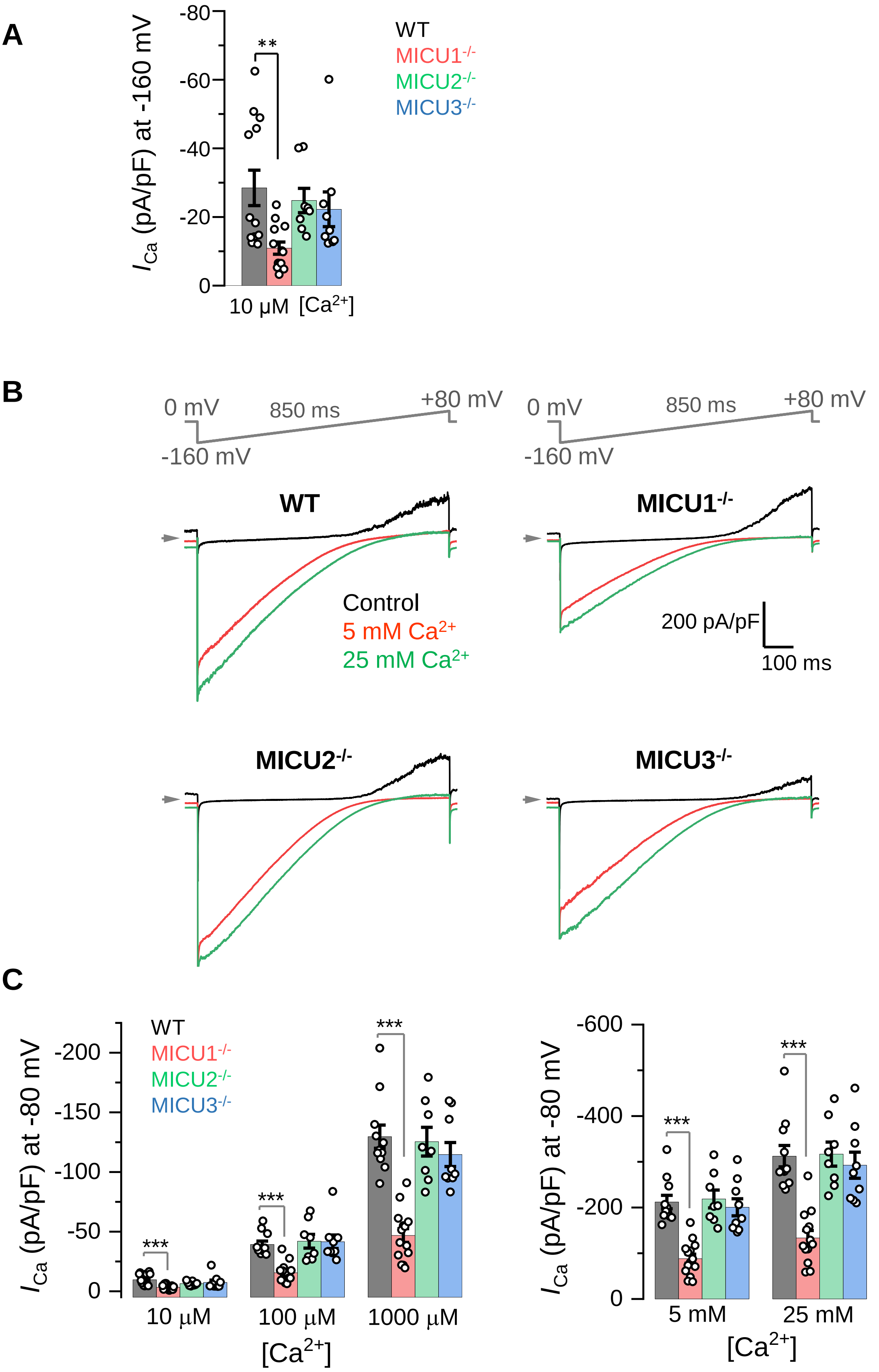
A
$I_{Ca}$ (pA/pF) at -160 mV
-80
-60
-40
-20
0
**
10 µM [Ca2+]
WT
MICU1-/-
MICU2-/-
MICU3-/-
B
0 mV
850 ms
+80 mV
-160 mV
WT
MICU1-/-
Control
5 mM Ca2+
25 mM Ca2+
200 pA/pF
100 ms
MICU2-/-
MICU3-/-
C
$I_{Ca}$ (pA/pF) at -80 mV
-200
-150
-100
-50
0
***
10 µM
100 µM
1000 µM
[Ca2+]
-600
-400
-200
0
5 mM
25 mM
[Ca2+]

**Fig. S5**

**Fig. S6. Rescue of EMRE expression in *MICU1*$^{-/-}$ does not rescue $I_{Ca}$.** (**A** to **C**) Western blots showing the expression levels of EMRE (A), MCU (B) and MCUb (C) in cells with *WT* MCU complex and *MICU1*$^{-/-}$ ($n$ = 3 independent samples each). (**D**) (*Left*) Western blots showing EMRE protein level in *WT* and *MICU1*$^{-/-}$ (before and after EMRE overexpression). (*Right*) Graph represents quantification of Western blot ($n$ = 4 independent samples each). (**E**) Representative inward $I_{Ca}$ in *WT*, *MICU1*$^{-/-}$, and when EMRE was overexpressed in *MICU1*$^{-/-}$ (*MICU1*$^{-/-}$ + EMRE) upon exposure to 100 μM and 1000 μM $[Ca^{2+}]_{cyto}$. (**F**) $I_{Ca}$ amplitudes measured at -160 mV in *MICU1*$^{-/-}$ overexpressing EMRE (*MICU1*$^{-/-}$ + EMRE, $n$ = 8) as well as in *MICU1*$^{-/-}$ and *WT*. *WT* and *MICU1*$^{-/-}$ data are the same as in Fig. 1E. Mean ± SEM; one-way ANOVA with post-hoc Tuckey test. *$p< 0.05$; **$p< 0.01$; ***$p< 0.001$.

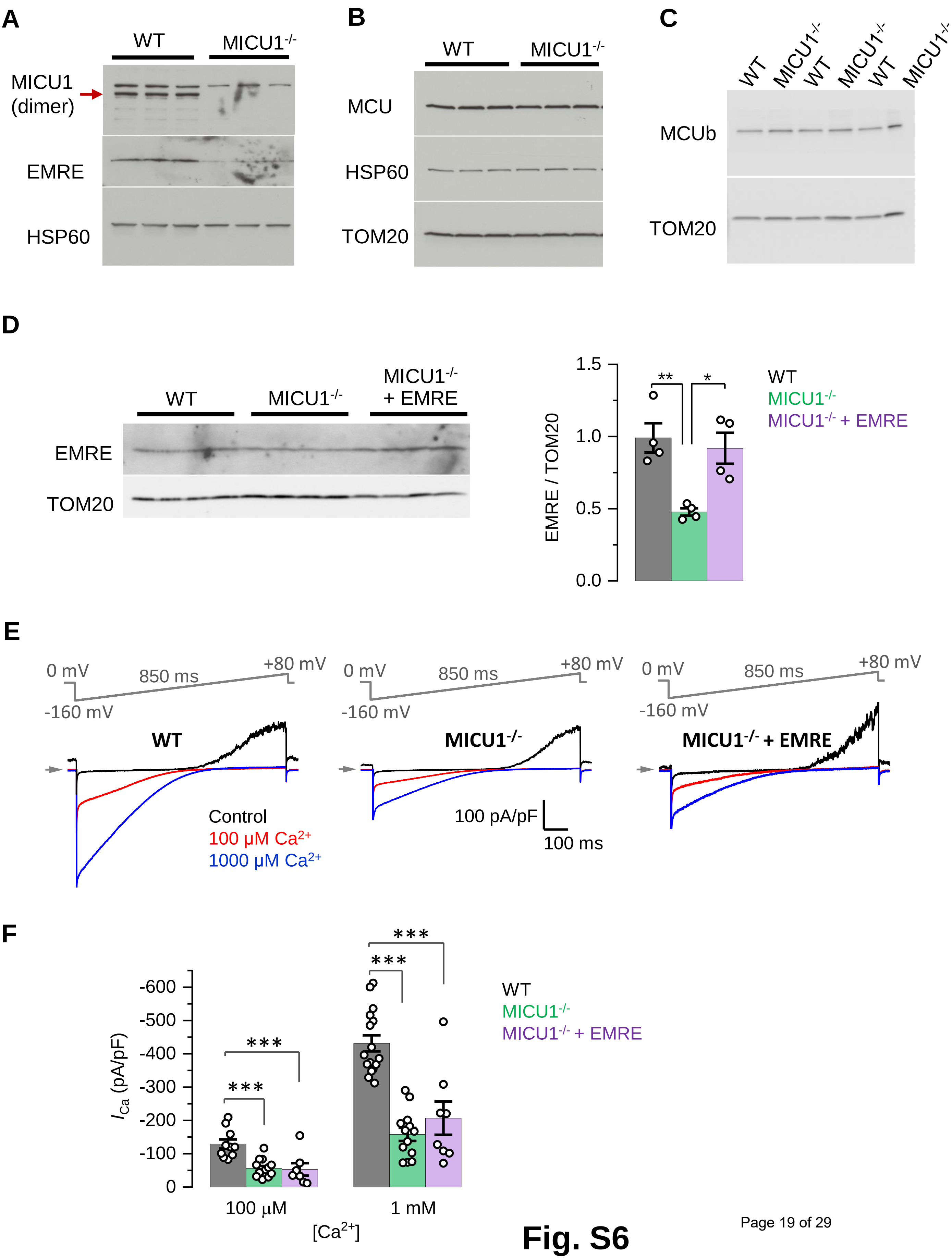
A
WT
MICU1-/-
MICU1
(dimer)
EMRE
HSP60
B
WT
MICU1-/-
MCU
HSP60
TOM20
C
WT
MICU1-/-
WT
MICU1-/-
WT
MICU1-/-
MCUb
TOM20
D
WT
MICU1-/-
MICU1-/- + EMRE
EMRE
TOM20
EMRE / TOM20
1.5
1.0
0.5
0.0
**
*
WT
MICU1-/-
MICU1-/- + EMRE
E
0 mV
850 ms
+80 mV
-160 mV
WT
MICU1-/-
MICU1-/- + EMRE
Control
100 μM Ca2+
1000 μM Ca2+
100 pA/pF
100 ms
F
ICa (pA/pF)
-600
-500
-400
-300
-200
-100
0
***
***
***
***
100 μM
1 mM
[Ca2+]
WT
MICU1-/-
MICU1-/- + EMRE

Fig. S6

**Fig. S7. Drp1 does not affect the currents mediated by the MCU complex or their phenotype in *MICU1*$^{-/-}$.** (**A**) $I_{Ca}$ amplitudes at -160 mV (*left*) and -80 mV (*right*) in mitoplasts from MEFs with Drp1 (*Drp*$^{+/+}$) and without Drp1 (*Drp1*$^{-/-}$). (*n* = 17-19) Mean ± SEM. (**B**) $I_{Na}$ amplitudes at -160 mV (*left*) and -80 mV (*right*) in mitoplasts from MEFs with Drp1 (Drp$^{+/+}$, *n* = 8) and without Drp1 (*Drp1*$^{-/-}$, *n* = 21). Mean ± SEM. (**C** to **F**) Current phenotypes of *MICU1*$^{-/-}$ in mitoplasts isolated from MEFs with an intact Drp1. (C) Representative $I_{Ca}$ (*blue*) and $I_{Na}$ (*red*) recorded from the *WT* (*n* = 7) and *MICU1*$^{-/-}$ (*n* = 12) mitoplasts exposed to 1 mM $[Ca^{2+}]_{cyto}$ or 110 mM $[Na^{+}]_{cyto}$. Amplitudes of $I_{Na}$ (D) and $I_{Ca}$ (E) measured at -80 mV. (F) Ratio between $I_{Ca}$ and $I_{Na}$ measured in the same mitoplast. Mean ± SEM; unpaired t-test, two-tailed; ***$p < 0.001$.

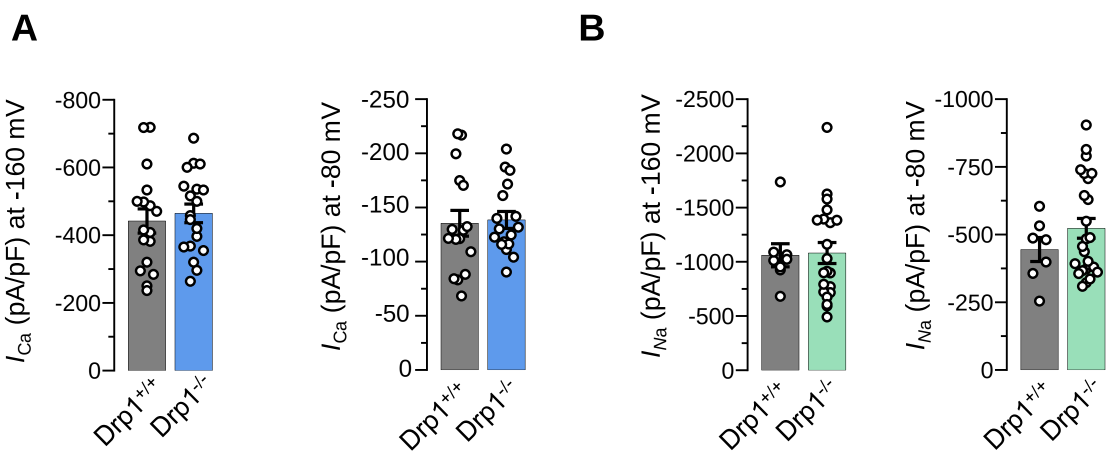


#### Mitoplasts from MEFs with intact Drp1

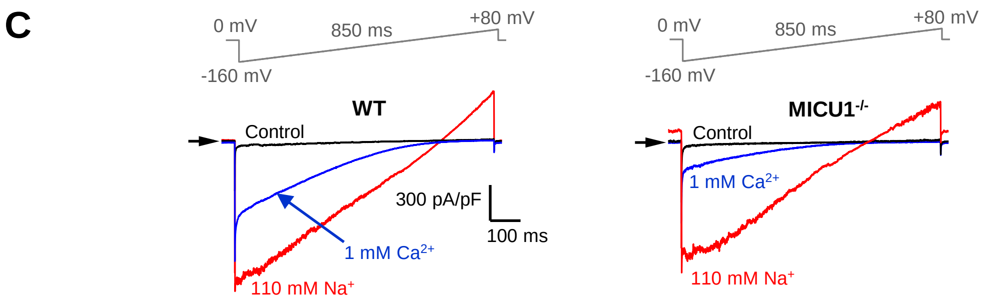


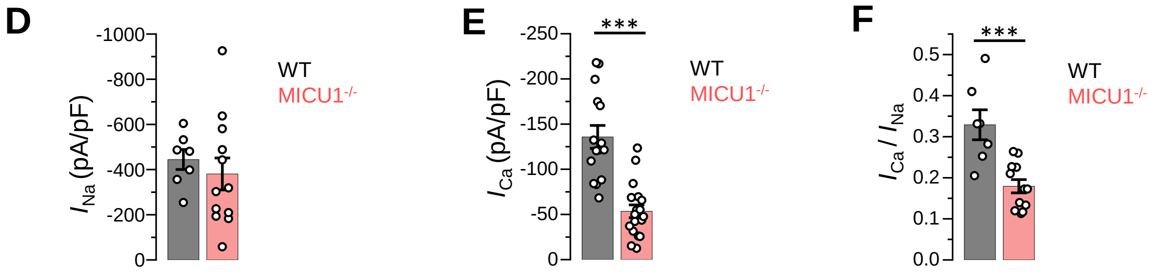




**Fig. S7**

**Fig. S8. Expression levels of different MCU subunits in *MICU2$^{-/-}$* MEFs and the open probability of MCU at different potentials.** (**A** to **C**) Western blots showing the expression levels of MICU2 (A), MCU, EMRE, MICU3 (B) and MICU1 dimers (C) in *WT* and *MICU2$^{-/-}$* cells ($n$ = 3–6 independent samples each). For detection of MICU1 dimers, samples were prepared in Laemmli buffer without β-mercaptoethanol.

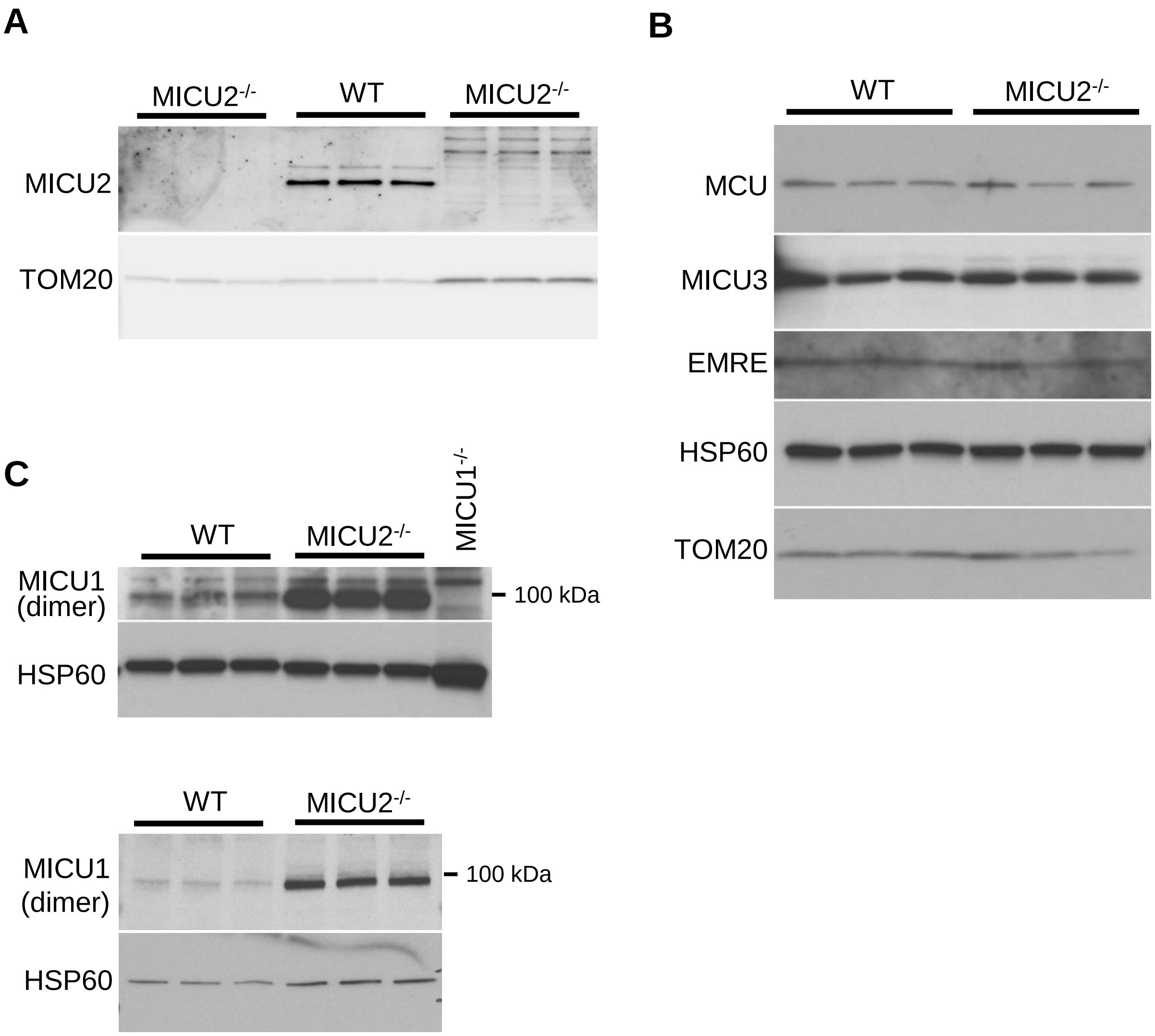
A
MICU2-/-
WT
MICU2-/-
MICU2
TOM20
B
WT
MICU2-/-
MCU
MICU3
EMRE
HSP60
TOM20
C
WT
MICU2-/-
MICU1-/-
MICU1 (dimer)
100 kDa
HSP60
WT
MICU2-/-
MICU1 (dimer)
100 kDa
HSP60



**Fig. S8**

**Fig. S9. Matrix $Ca^{2+}$ does not inhibit $I_{Ca}$.** (*Upper Panels*) Inward $I_{Ca}$ in the presence of 0 (*left*), 400 nM (*middle*) and 400 μM (*right*) $[Ca^{2+}]_{mito}$ (pipette solution). $[Ca^{2+}]_{cyto}$ was 100 μM, 1 mM or 5 mM. (*Lower Panel*) $I_{Ca}$ amplitudes at 0 (*n*=3-5), 400 nM (*n*=4) or 400 μM (*n*=3) $[Ca^{2+}]_{mito}$. $I_{Ca}$ was measured at -160 mV and in different $[Ca^{2+}]_{cyto}$ as indicated. Mean ± SEM; one-way ANOVA with post-hoc Tuckey test.

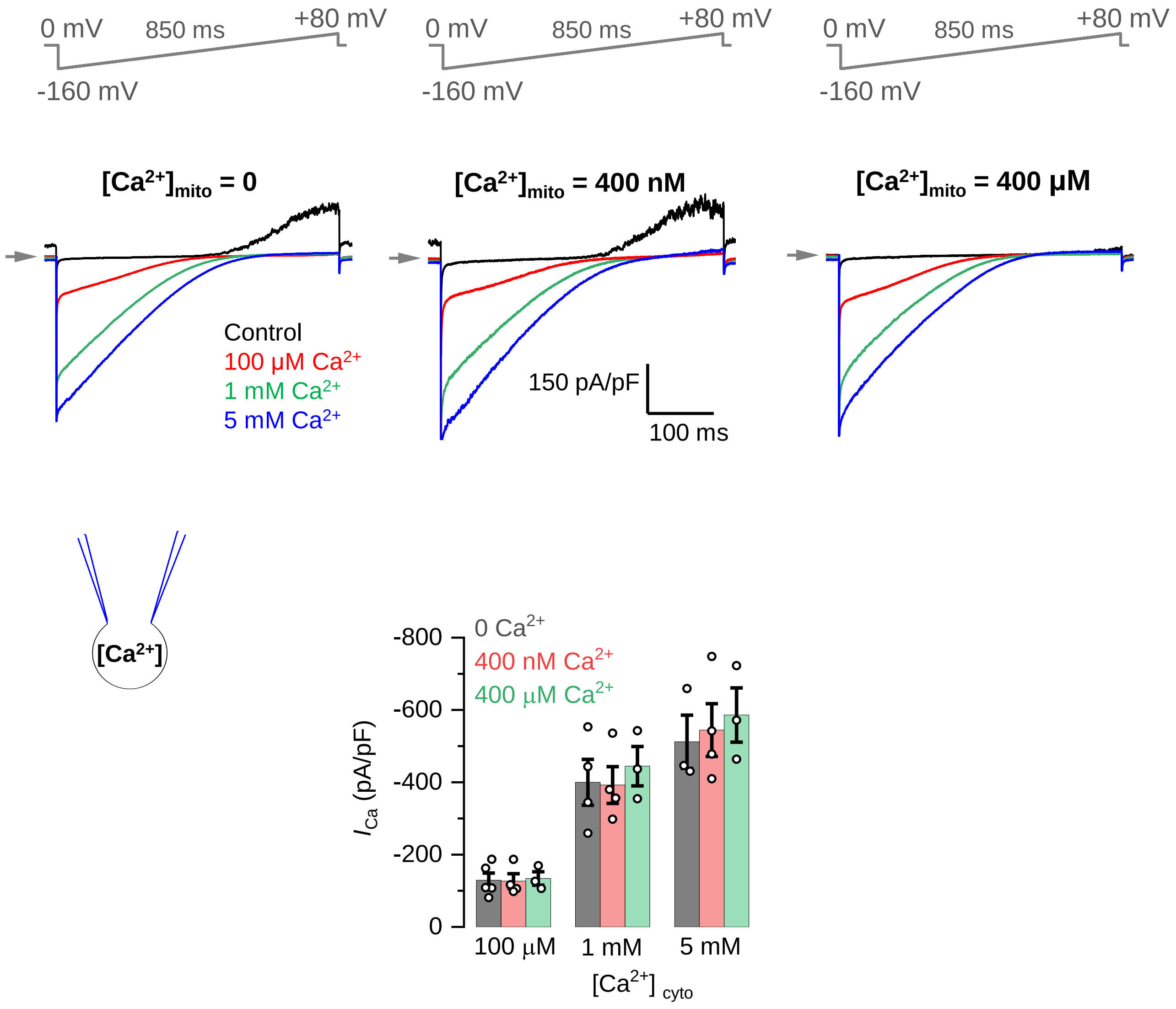
0 mV
850 ms
+80 mV
-160 mV
[Ca2+]mito = 0
[Ca2+]mito = 400 nM
[Ca2+]mito = 400 µM
Control
100 µM Ca2+
1 mM Ca2+
5 mM Ca2+
150 pA/pF
100 ms
[Ca2+]
0 Ca2+
400 nM Ca2+
400 µM Ca2+
ICa (pA/pF)
-800
-600
-400
-200
0
100 µM
1 mM
5 mM
[Ca2+] cyto



**Fig. S9**

**Fig. S10. Open probability of the MCU channel in *WT* and *MICU1*$^{-/-}$.** (**A** to **C**) Open probability of the MCU channel in *WT* ($n$=5-6) and *MICU1*$^{-/-}$ ($n$=5) at -40 mV (A) -80 mV (B) and at -120 mV (C). The same *WT* and knockout data were used as in Fig. 4D, but presented to show data distribution. Mean ± SEM; unpaired t-test, two-tailed; $^{**}p < 0.01$.

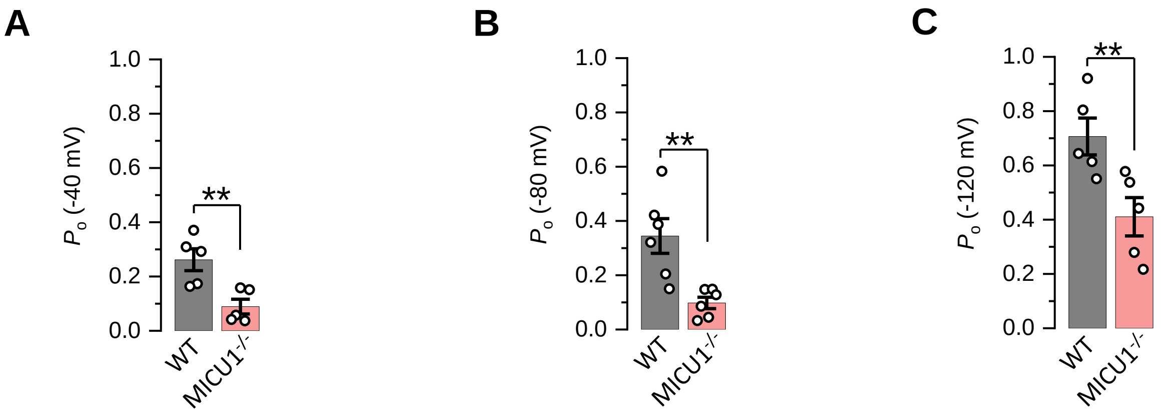


### Fig. S10

**Table S1. MICU1 effect on MCU as determined by previous electrophysiological experiments.**

| Citation | MCU current | |
|---|---|---|
| | **Low $[Ca^{2+}]_{cyto}$** | **High $[Ca^{2+}]_{cyto}$** |
| Hoffman et al., 2013 (*44*) | | Inhibition |
| Patron et al., 2014 (*25*)<br>Lipid bilayer experiments;<br>EMRE subunit, essential for MCU activity and MICU1 interaction, was absent | No effect | Activation |
| Vais et al., 2016 (*43*) | | No change |
| Kamer et al., 2018 (*42*) | | No change |

**Table S2. MICU1 effect on MCU as determined by previous $Ca^{2+}$ imaging experiments.**

| Reference | Effect of MICU1 on mitochondrial $Ca^{2+}$ uptake | |
|---|---|---|
| | Low $[Ca^{2+}]_{cyto}$ | High $[Ca^{2+}]_{cyto}$ |
| Mallilankaraman et al., 2012 (*28*) | Inhibition | No effect |
| Hoffman et al., 2013 (*44*) | Inhibition | Inhibition |
| Plovanich et al., 2013 (*23*) | | Activation |
| Csordas et al., 2013 (*29*) | Inhibition | Activation |
| de la Fuente et al., 2014 (*31*) | Inhibition | Activation |
| Kamer and Mootha, 2014 (*24*) | Inhibition | |
| Logan et al., 2014 (*30*) | Inhibition | No effect |
| Patron et al., 2014 (*25*) | Inhibition | Inhibition |
| Hall et al., 2014 (*73*) | | Inhibition |
| Antony et al., 2016 (*36*) | Inhibition | Activation |
| Liu et al., 2016 (*32*) | Inhibition | Activation |
| Bhosale et al., 2017 (*74*) | Inhibition | |
| Kamer et al., 2017 (*51*) | Inhibition | |
| Paillard et al., 2018 (*34*) | Inhibition | Activation |
| Phillips et al., 2018 (*33*) | Inhibition | |
